# Seasonal influenza: Modelling approaches to capture immunity propagation

**DOI:** 10.1101/637074

**Authors:** Edward M. Hill, Stavros Petrou, Simon de Lusignan, Ivelina Yonova, Matt J. Keeling

**Affiliations:** Zeeman Institute: Systems Biology and Infectious Disease Epidemiology Research (SBIDER), University of Warwick, Coventry, CV4 7AL, United Kingdom; Mathematics Institute, University of Warwick, Coventry, CV4 7AL, United Kingdom; Warwick Clinical Trials Unit, Warwick Medical School, University of Warwick, Coventry, CV4 7AL, United Kingdom; Department of Clinical and Experimental Medicine, University of Surrey, Guildford, GU2 7XH, United Kingdom; Royal College of General Practitioners, London, NW1 2FB, United Kingdom; School of Life Sciences, University of Warwick, Coventry, CV4 7AL, United Kingdom

## Abstract

Seasonal influenza poses serious problems for global public health, being a significant contributor to morbidity and mortality. In England, there has been a long-standing national vaccination programme, with vaccination of at-risk groups and children offering partial protection against infection. Transmission models have been a fundamental component of analysis, informing the efficient use of limited resources. However, these models generally treat each season and each strain circulating within that season in isolation. Here, we amalgamate multiple data sources to calibrate a susceptible-latent-infected-recovered type transmission model for seasonal influenza, incorporating the four main strains and mechanisms linking prior season epidemiological outcomes to immunity at the beginning of the following season. Data pertaining to nine influenza seasons, starting with the 2009/10 season, informed our estimates for epidemiological processes, virological sample positivity, vaccine uptake and efficacy attributes, and general practitioner influenza-like-illness consultations as reported by the Royal College of General Practitioners (RCGP) Research and Surveillance Centre (RSC). We performed parameter inference via approximate Bayesian computation to assess strain transmissibility, dependence of present season influenza immunity on prior protection, and variability in the influenza case ascertainment across seasons. This produced reasonable agreement between model and data on the annual strain composition. Parameter fits indicated that the propagation of immunity from one season to the next is weaker if vaccine derived, compared to natural immunity from infection. Projecting the dynamics forward in time suggests that while historic immunity plays an important role in determining annual strain composition, the variability in vaccine efficacy hampers our ability to make long-term predictions.

## Introduction

As a significant contributor to global morbidity and mortality, seasonal influenza is an ongoing public health concern. Worldwide, these annual epidemics are estimated to result in about three to five million cases of severe illness, and about 290,000 to 650,000 respiratory deaths [1]. In England, seasonal influenza inflicts a stark burden on the health system during winter periods, being linked with approximately 10% of all respiratory hospital admissions and deaths [2].

Influenza vaccination can offer some protection against seasonal influenza infection for the individual, while contributing to reduced risk of ongoing transmission via establishment of herd immunity [3, 4]. Influenza vaccines are designed to protect against three or four different influenza viruses; two influenza A viruses (an A(H1N1)pdm09 subtype and A(H3N2) subtype) and either one or two influenza B viruses (covering one or both of the B/Yamagata and B/Victoria lineages). In 2013, 40% of countries worldwide recommended influenza vaccination in their national immunisation programmes, although vaccine uptake varies [5–7].

For England (and elsewhere), the need to deploy updated vaccines on an annual basis means influenza vaccination programmes are costly. Yet, predictions of vaccine impact are difficult due to stochasticity in the annual strain composition, the potential misalignment between vaccine and the dominant co-circulating strains and the interaction between multiple seasons. Analysis informing the efficient use of limited resources is therefore vital, with the use of quality-assured analytical models advocated [8].

The previous ten years have seen the fruitful development of influenza transmission models tied to available real-word and experimental data sources [9–15]. Furthermore, combining parameterised transmission models with health economic evaluations permits assessments of changes to vaccination programmes, providing evidence to inform vaccine policy decisions [16, 17]. An eminent study by Baguelin *et al.* [12] connected virological data from England and Wales to a deterministic epidemiological model within a Bayesian inference framework. The transmission model was subsequently interfaced with a health economic analysis model, leading to the recommendation of introducing a paediatric seasonal influenza vaccination programme in the United Kingdom [17]. Subsequent studies have analysed the effect of mass paediatric influenza vaccination on existing influenza vaccination programmes in England and Wales [18], evaluated cost-effectiveness of quadrivalent vaccines versus trivalent vaccines [19], and cost effectiveness of high-dose and adjuvanted vaccine options in the elderly [20].

Prior modelling studies have typically treated each season and each strain circulating within that season independently. Though there are instances of model conceptualisations incorporating waning immunity for natural infection and vaccination, these assumed one generic influenza virus [13], or treated one or both of the influenza A and influenza B types as sole entities rather than explicitly distinguishing between the two common influenza A subtypes and two co-circulating influenza B lineages [9, 11, 16]. Here, we present a multi-strain, non-age structured, susceptible-latent-infected-recovered (SEIR) type transmission model for influenza incorporating a mechanism to link prior season epidemiological outcomes to immunity at the beginning of the following season. Incorporation of a mechanism for the building and propagation of immunity facilitates investigation of the impact of exposure in the previous influenza season, through natural infection or vaccination, on the disease transmission dynamics and overall disease burden in subsequent years. Accordingly, we sought insights to aid understanding of the longer-term dynamics of the influenza virus and its interaction with immunity at the population level.

In this study, we examine the contribution of the differing sources of immunity propagation between years on seasonal influenza transmission dynamics. To this end, we amalgamate multiple sources of epidemiological and vaccine data for England covering the last decade, and fit model outcomes to the available longitudinal data of seasonal rates of general practice (GP) consultations for influenza-like-illness (ILI), scaled by virological surveillance information. We demonstrate that natural infection plays a more prominent role in propagation of immunity to the next influenza season compared to residual vaccine immunity. We conclude by inspecting forward projections under disparate vaccine efficacy assumptions to determine long-term patterns of seasonal influenza infection.

## Methods

### Historical data

Throughout, we define a complete epidemiological influenza season to run from week 36 to week 35 of the subsequent calendar year. We chose week 36 (which typically corresponds to the first full week in September) as the start week to match the start of the epidemic with the reopening of schools.

### Consultations in General Practices

Since January 1967, the Royal College of General Practitioners (RCGP) Research and Surveillance Centre (RSC) has monitored the activity of acute respiratory infections in GPs. The Weekly Return reports the number of persons in the monitored RCGP RSC network practices registered population consulting for ILI (a medical diagnosis of possible influenza with a set of common symptoms, cough and measured or reported fever ≥ 38°C, with onset within the last 10 days [21]).

Importantly, the dataset is nationally representative both demographically and spatially; the demographics of the sample population closely resembling the country as a whole, with the RCGP RSC network of practices spread across England in order to reflect the distribution of the population. In May 2017, the RCGP RSC network included 1,835,211 patients (approximately 3.3% of the total population in England) from 174 practices [22].

For the influenza seasons 2009/2010 until 2017/2018 inclusive, RCGP RSC provided weekly, age-stratified records containing the size of the monitored RCGP RSC population, the number of individuals in the monitored population consulting for ILI, and ILI rates per 100,000. These data were aggregated across ages to produce weekly ILI rates per 100,000 for the entire sample population. We then summed the weekly ILI rates (weeks 36 through to week 35 in the following calendar year) to produce season-by-season ILI rates per 100,000. Expanded information pertaining to the data extraction is provided in Section 1.1 of the Supporting Information.

### Respiratory Virus RCGP RSC Surveillance

To complement syndromic surveillance, a subset of GPs in the RCGP RSC submit respiratory samples from patients presenting in primary care with an ILI. These respiratory samples undergo virological testing to ascertain presence or absence of influenza viruses. Given not all patients reporting ILI are infected with an influenza virus, these virological sample positivity data yield the fraction of ILI reported primary care consultations that were attributable to influenza.

We collected relevant data from figures within Public Health England (PHE) annual influenza reports (for the 2009/10 to 2017/18 influenza seasons inclusive) [23, 24], and PHE Weekly National Influenza reports (for the 2013/14 influenza season onwards) [25]. (For full details, see Section 1.2 of the Supporting Information.) By scaling the longitudinal data of GP consultations for ILI by the virological surveillance information we garnered an overall estimate of ILI GP consultations attributable to influenza.

### Circulating strain composition

We derived seasonal rates of primary care consultations attributable to each of the two influenza A subtypes (A(H1N1)pdm09 and A(H3N2)) and the two influenza B lineages (B/Victoria and B/Yamagata) per 100,000 population. We computed the circulating strain distribution in each influenza season using publicly available data from FluNet [26], a global web-based tool for influenza virological surveillance, using data for the United Kingdom (additional details are provided in Section 1.3 of the Supporting Information).

### ILI GP consultations attributable to influenza

We applied the circulating strain composition quantities to the overall estimate of ILI GP consultations attributable to influenza, disaggregating the estimate for all influenza strains of interest into strain-specific quantities:

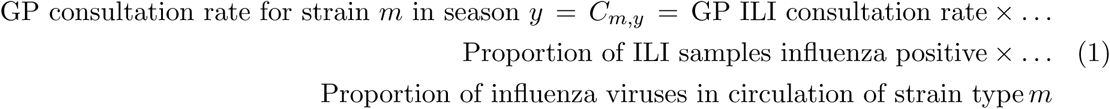

These computations resulted in seasonal rates of GP consultations attributable for the two influenza A subtypes and two influenza B lineages per 100,000 population (Fig. 1(a)). Uncertainty distributions for these values were obtained via bootstrapping given the sample sizes for the three components.

**Fig. 1:**
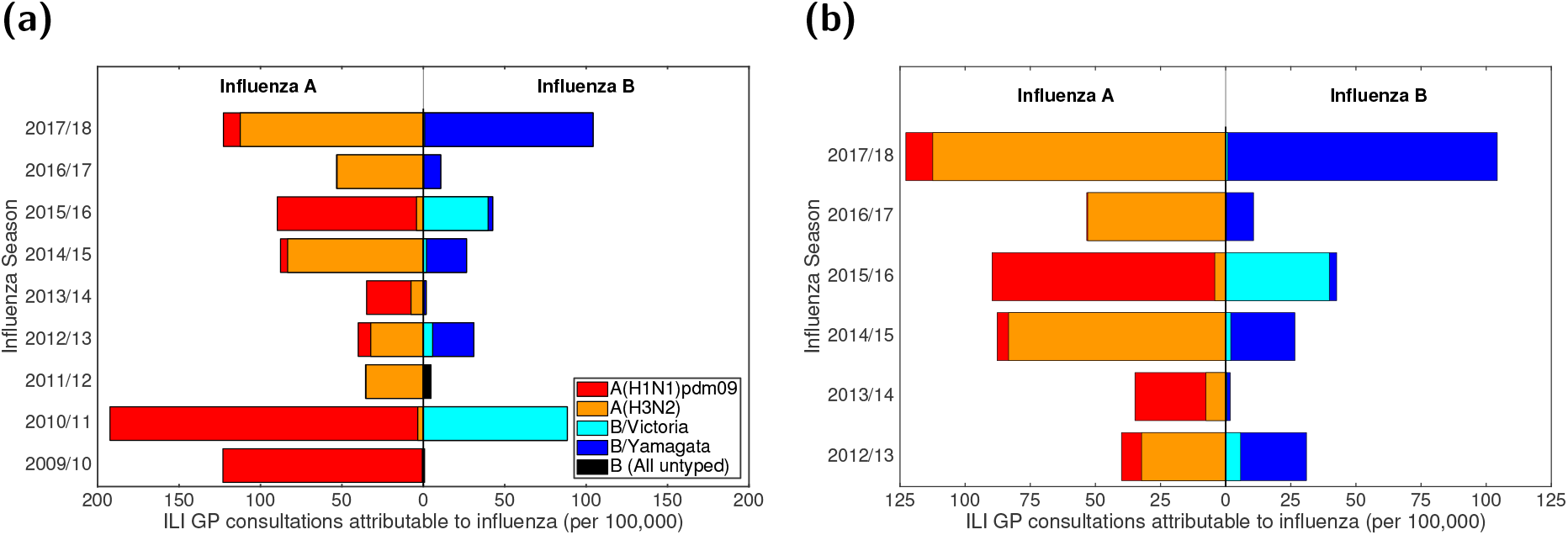
Empirical, strain-stratified data for ILI GP consultations attributable to influenza per 100,000 population. In both panels, stacked horizontal bars in the left-half depict the cumulative total of ILI GP consultations attributable to type A influenza per 100,000 population (red shading denoting the A(H1N1)pdm09 subtype, orange shading the A(H3N2) subtype). In an equivalent manner, on the right-half of each panel stacked horizontal bars present similar data for type B influenza (cyan shading denoting the B/Victoria lineage, dark blue shading the B/Yamagata lineage). **(a)** Influenza seasons 2009/10 to 2017/18 (inclusive). In the 2009/10 and 2011/12 influenza seasons, all samples identified as influenza B were untyped (represented by black filled bars). **(b)** Influenza seasons 2012/13 to 2017/18 (inclusive), the time span for which we performed parameter inference.

### Vaccine uptake and efficacy

For the 2009/10 influenza season, we acquired vaccine uptake profiles from survey results on H1N1 vaccine uptake amongst patient groups in primary care [27]. For the 2010/11 influenza season onward we collated vaccine uptake information from PHE official statistics [24, 25].

We took age adjusted vaccine efficacy estimates for each historical influenza season (2009/2010-2017/2018 inclusive) from publications detailing end-of-season seasonal influenza vaccine effectiveness in the United Kingdom [28–34]. In seasons where equivalent publications were not available, we used mid-season or provisional end-of-season age adjusted vaccine efficacy estimates from PHE reports [35, 36].

There has been substantial variation in vaccine efficacy between seasons and also within seasons across strains, with estimates spanning 23-92% (Table 1). Further, there have been two instances of a mismatch between the vaccine and circulating viruses causing the vaccine to be ineffective (i.e. 0% efficacy) against a particular strain group (B/Yamagata in the 2010/11 influenza season, A(H3N2) in the 2017/18 influenza season). As a sole exception, for the 2009/10 influenza season we used the pandemic vaccine (rather than the seasonal influenza vaccine efficacy) to inform efficacy against A(H1N1)pdm09, with the effectiveness against all other strain types set to zero [28].

**Table 1:**
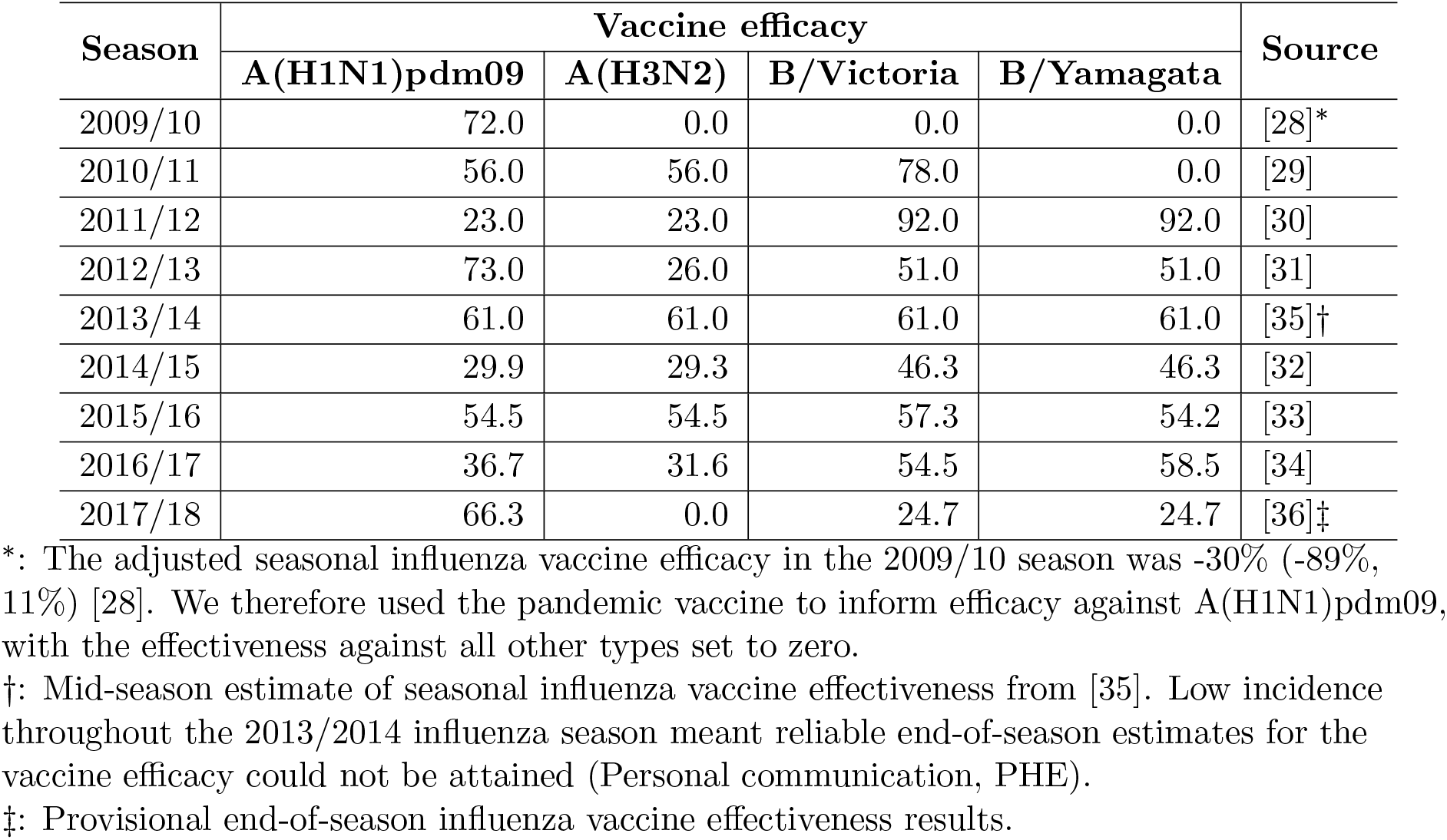
Age adjusted, influenza vaccine efficacy point estimates (by season and strain type) All estimates are presented as percentages. The empirical adjusted influenza vaccine efficacy estimates by season and strain type (presented in Table S1) did not provide individual vaccine efficacy estimates for each influenza A subtype and influenza B lineage. We therefore implemented a series of assumptions to produce the strain-specific, vaccine efficacy point estimates used within our study (described in Section 1.5 of the Supporting Information)).

The empirical data did not provide individual vaccine efficacy estimates for each influenza A subtype and influenza B lineage (Table S1). We therefore used a set of simplifying assumptions to produce the strain-specific, vaccine efficacy point estimates used within our study (Table 1). The time-varying vaccine uptake data, together with the season-specific information on vaccine efficacy, act as forcing for the within-season epidemiological model (details are described in Sections 1.4-1.5 of the Supporting Information).

### Modelling overview

Our motivation was to study the role of immunity propagation in seasonal influenza transmission dynamics by developing and applying a modelling framework incorporating multiple mechanisms linking prior season infection (or vaccination) to immunity in subsequent seasons.

Influenza transmission dynamics are highly complex, with many temporal and structural heterogeneities that will influence the precise pattern of recorded infection. Consequently, our aim was to capture the general trends in the data, such as the pattern of high and low infection levels, rather than the precise values for each influenza season. We, therefore, deliberately chose a parsimonious mechanistic modelling framework, without age-structure, to highlight the impact of immunity propagation.

Following the visualisation convention used in [12], we display a directed acyclic graph representing the model structure and incorporation of the data streams (Fig. 2). The model takes the form of a deterministic continuous-time set of ordinary differential equations (ODEs), which determines the within-season epidemiological dynamics, and a discrete-time map, informing the propagation of immunity from one season to the next.

**Fig. 2:**
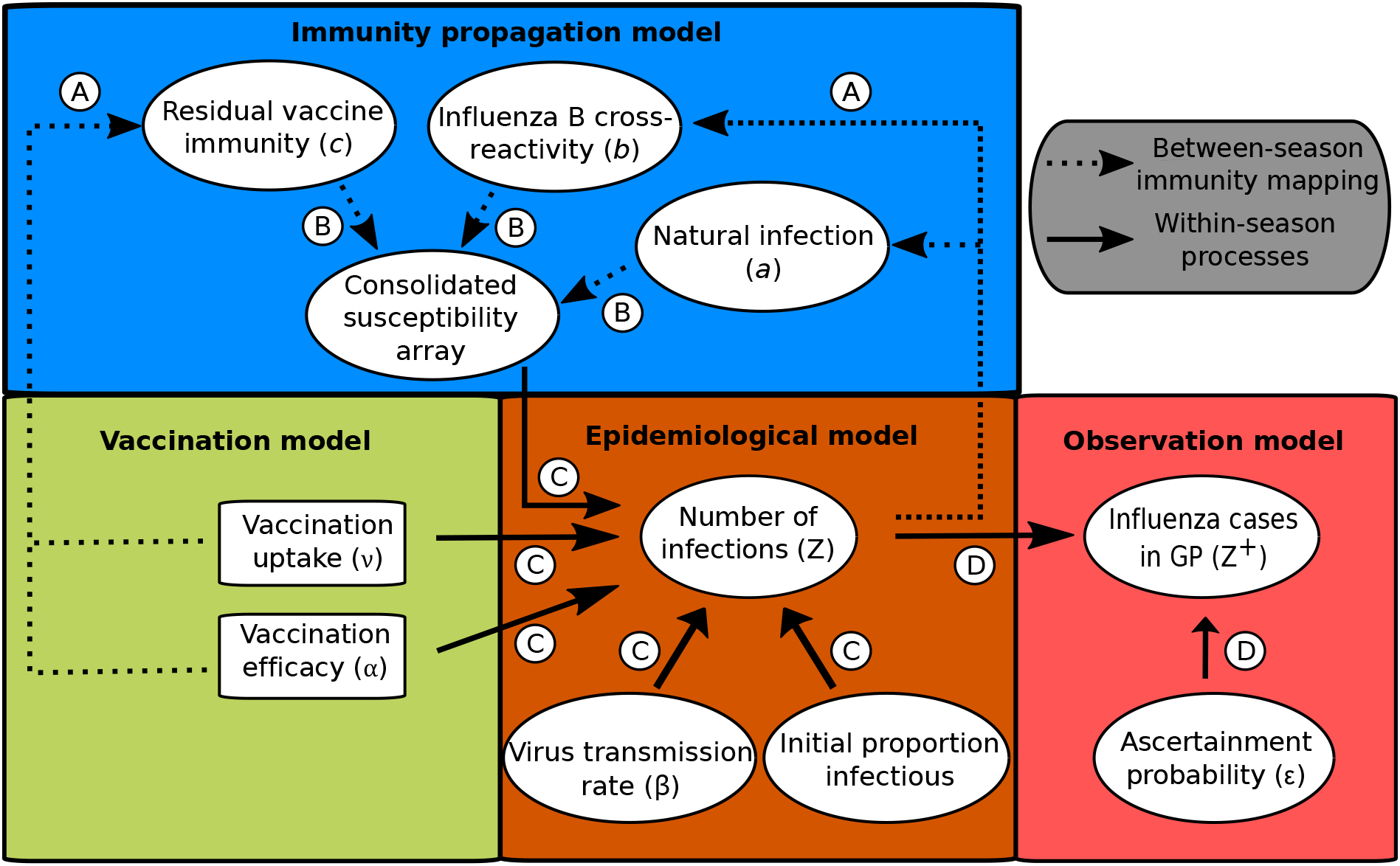
Schematic showing the links between the vaccination, immunity propagation, epidemiological and observation model components. We adopt the visualisation conventions of [12], with ellipses indicating variables, and rectangles indicating data. Dotted arrows indicate relationships between prior season epidemiological outcomes and immunity propagation factors. Solid arrows indicate within-season processes. Circled capitalised letters indicate the relationships connecting the variables or data involved. These relationships are: process A, propagation of immunity as a result of exposure to influenza virus in the previous influenza season (through natural infection or vaccination); process B, modulation of current influenza season virus susceptibility; process C, estimation of influenza case load via the SEIR model of transmission; process D, ascertainment of cases through ILI recording at GP.

Our model captures the four strains targeted by the quadrivalent seasonal influenza vaccine, namely, two influenza A subtypes, A(H1N1)pdm09 and A(H3N2), and two influenza B lineages, B/Victoria and B/Yamagata. The multi-strain model construction permits exploration of cross-reactive immunity mechanisms and their impact on the disease dynamics. Multiple infections per influenza season and/or co-infection events were not permitted. In other words, it was presumed that individuals may only be infected by one strain of influenza virus per season, analogous to natural infection eliciting short-term cross immunity to all other strain types [37].

Next, we summarise, in turn, the vaccination, immunity propagation, epidemiological and observation model components.

### Vaccination model

Epidemiological states were compartmentalised based on present season vaccine status (indexed by *N* for non-vaccinated or *V* for vaccinated). We obtained weekly time-varying vaccine uptake rates, *ν*, at the population level by computing a weighted average of the individual age group uptake values, thereby accounting for the population distribution in each given influenza season. We assumed the rate of vaccination *ν* to be constant over each weekly period. Vaccine efficacy depended upon the extent to which the strains in the vaccine matched the circulating strain in that influenza season (Table 1).

We assumed a ‘leaky’ vaccine, offering partial protection to every vaccinated individual, thus acting to reduce the overall susceptibility of the given group receiving vaccination. The ‘leaky’ vaccine action contrasts with an ‘all-or-nothing’ vaccine assumption, which assumes complete protection to a subset of the vaccinated individuals but no protection in the remainder of vaccinated individuals [38]. Explicitly, under a ‘leaky’ vaccine action with a vaccine efficacy against strain *m* of *α_m_*, the relative susceptibility of the vaccinated group towards strain *m* is 1 *− α_m_*. Additionally, we assumed those administered vaccines had unmodified transmission (i.e. those receiving the ‘leaky’ vaccine were equally as infectious as those who did not). Vaccination is modelled irrespective of infection status (susceptible, latent, infected and recovered individuals), as vaccine history impacts the immunity levels for next season.

### Immunity propagation model

A novel aspect of our model framework was the potential for exposure to influenza virus in the previous influenza season, through natural infection or vaccination, to modulate current influenza season susceptibility. Accordingly, we tracked immunity derived from natural infection and vaccination separately (Fig. 2, process A), requiring ten distinct exposure history groupings. Three susceptibility modifying factors of interest were: (i) modified susceptibility to strain *m* given infection by a strain *m* type virus the previous season, denoted *a*; (ii) carry over cross-reactivity protection between influenza B lineages, denoted *b* (to account for infection with one influenza B virus lineage being potentially beneficial in protecting against subsequent infection with either influenza B virus lineage [39]); (iii) residual strain-specific protection carried over from the prior season influenza vaccine, denoted *c_m_*. We mandated that 0 *< a*,*b*,*c_m_ <* 1. We let *f* (*h, m*) denote, for those in exposure history group *h*, the susceptibility to strain *m*. The collection of ten exposure history groupings and associated strain-specific susceptibilities were consolidated into a single susceptibility array (Fig. 2, process B; Fig. 3).

**Fig. 3:**
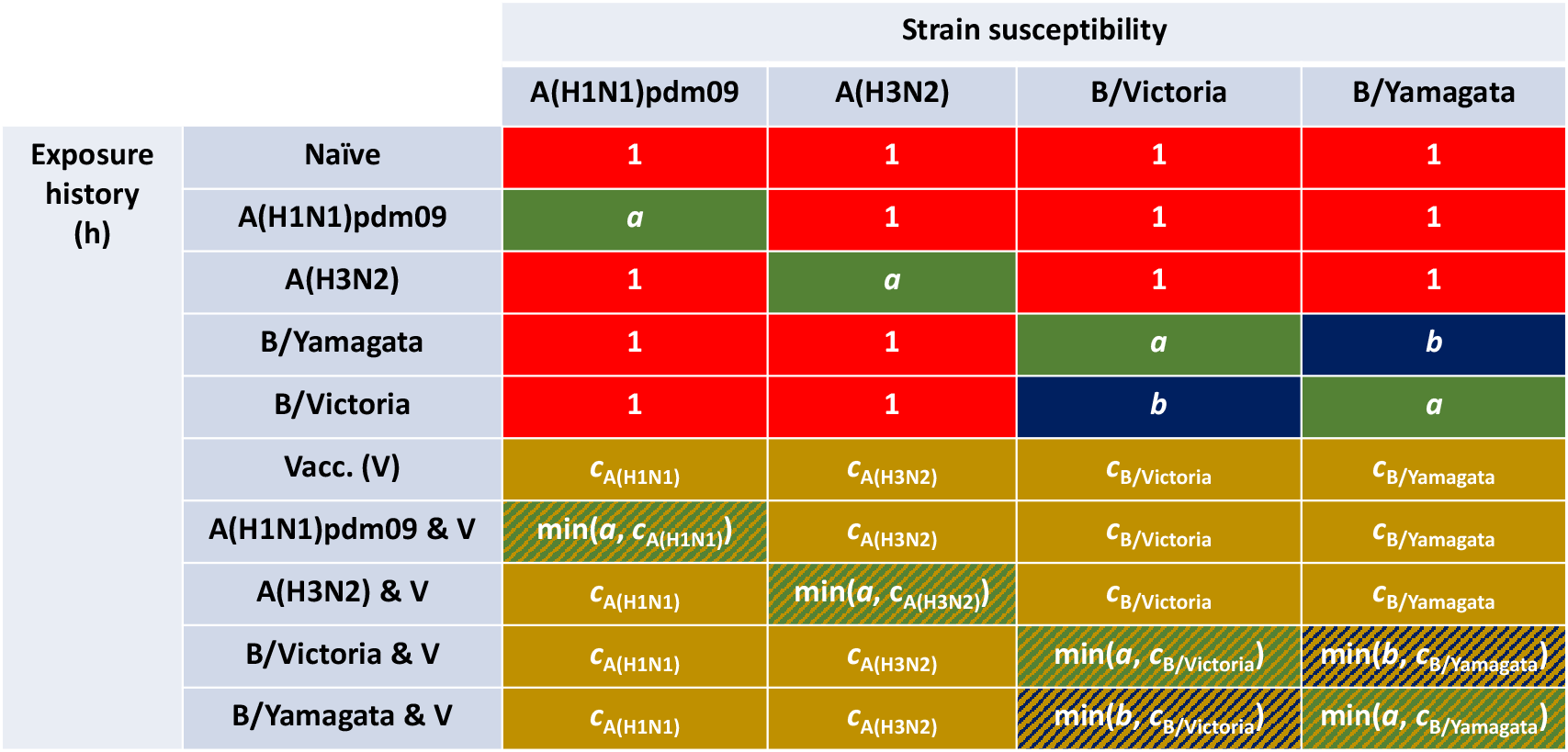
Infographic presenting the interaction between exposure history and susceptibility. The interaction between exposure history *h* and susceptibility to strain *m* in the current season, *f* (*h, m*), was classified into ten distinct groups: One group for the naive (uninfected and not vaccinated, row 1); one group per strain, infected but not vaccinated (rows 2-5); one group for those vaccinated and experiencing no natural infection (row 6); one group per strain for being infected and vaccinated (rows 7-10). We let *a* denote modified susceptibility to strain *m* given infection by a strain *m* type virus the previous season (dark green shading), *b* modified susceptibility due to cross-reactivity between type B influenza lineages (dark blue shading), and *c_m_* the change in susceptibility to strain *m* given vaccination in the previous season (gold shading). Unmodified susceptibilities retained a value of 1 (red shading). We enforced 0 *< a*,*b*,*c_m_<* 1.

We assumed the immunity propagation due to vaccination in the previous season (*c_m_*) to be associated to the strain-specific vaccine efficacy in the previous season 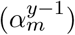. In particular, we introduce a linear scaling factor *ξ*∈(0, 1), which reduces the level of vaccine derived immunity between seasons: 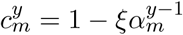.

On a related note, we applied a further assumption in order to parameterise susceptibility values amongst exposure groups capturing those both vaccinated and naturally infected in the prior season (Fig. 3: rows 7-10). For this collection of exposure histories, susceptibility to a subset of strains may conceivably be collectively modified via natural infection and vaccination immunity propagation pathways. In these instances, we treated the immunity propagation mechanisms independently, with the modified susceptibility set by the dominant immunity propagation entity (i.e. min(*a, c_m_*), min(*b, c_m_*)). In other words, we took a pessimistic stance by assuming no boosting of the immunity propagation response as a result of dual influenza virus exposure (from both natural infection and vaccination).

All three propagation parameters (*a, b, ξ*) were inferred from epidemiological data. A mathematical description of the immunity mapping between seasons is given in Section 2.2 of the Supporting Information.

In light of there being uncertainty around the precise time scale for which immunity to seasonal influenza viruses may be retained, with individual- and population-level models having estimated infection acquired immunity to wane over a timescale of two to ten years [40–42]), we also considered an extended variant of the immunity propagation model component.

In the model extension, we fit an additional parameter (*δ*) representing the proportion of those who began the influenza season in an exposure history group linked to natural infection (Fig. 3: rows 2-5, 7-10), and who were also unexposed to influenza virus during the current season (who at the end of of the current influenza season remained susceptible), that retained their pre-existing natural infection acquired immunity. Those keeping immunity arising from natural infection were mapped to the relevant prior infection exposure history group dependent upon current season vaccination status. The remaining proportion (1 *− δ*) transitioned in the same manner as in the original model, reverting to either the naive exposure history group or vaccinated only exposure history group (Figure S20). For clarity, we did not introduce a mechanism to confer vaccine-induced immunity beyond one influenza season (i.e. vaccine-induced immunity could be retained for, at most, a single additional influenza season).

With the more complex immunity structure, we did not gain noticeable improvements in correspondence of model outputs with the data (relative to fits with a model using the simpler immunity propagation setup). Therefore, we do not discuss any further here the alternative immunity propagation structure, instead giving a complete description of the extended model and associated outcomes in Section 5 of the Supporting Information.

### Epidemiological model

The within-season transmission dynamics were performed by an ODE epidemiological model based on SEIR-type dynamics. At the start of each season we passed into the epidemiological model the vaccination attributes (uptake and efficacy), immunity considerations, virus transmission rates and the initial proportion of the population infectious per strain. These collection of epidemiological inputs act as forcing parameters (Fig. 2, process C).

The epidemiological model is a deterministic, non-age, multi-strain structured compartmental based model capturing demography, influenza infection status (with susceptible-latent-infected-recovered, SEIR, dynamics) and vaccine uptake. For influenza, it is commonplace to also consider a risk-stratified population, divided into individuals at low or high risk of complications associated with influenza. However, it is health outcomes, rather than epidemiological processes, that are contingent upon risk status [12] and therefore this additional structure can be ignored for the purposes of this model. We assumed disease transmission to be frequency-dependent, such that fundamental quantities (like the basic reproductive ratio, *R*_0_) are unaffected by changes to population size.

We assumed exponentially-distributed latent and infectious periods (arising from rates of latency loss and recovery, respectively, being constant). Loss of latency rates *γ*_1*,m*_ were strain-dependent. From Lessler *et al.* [43], we chose rates corresponding to average latent periods of 1.4 days and 0.6 days for influenza A and influenza B associated strains respectively. Following Cauchemez *et al.* [44], we set the rate of loss of infectiousness *γ*_2_ = 1/3.8, corresponding to an average infectious period of 3.8 days, independent of strain.

We set the mortality rate based upon a gender-averaged life expectancy for England, approximated from male and female specific statistics from Office for National Statistics data [45] (see Table 2).

**Table 2:**
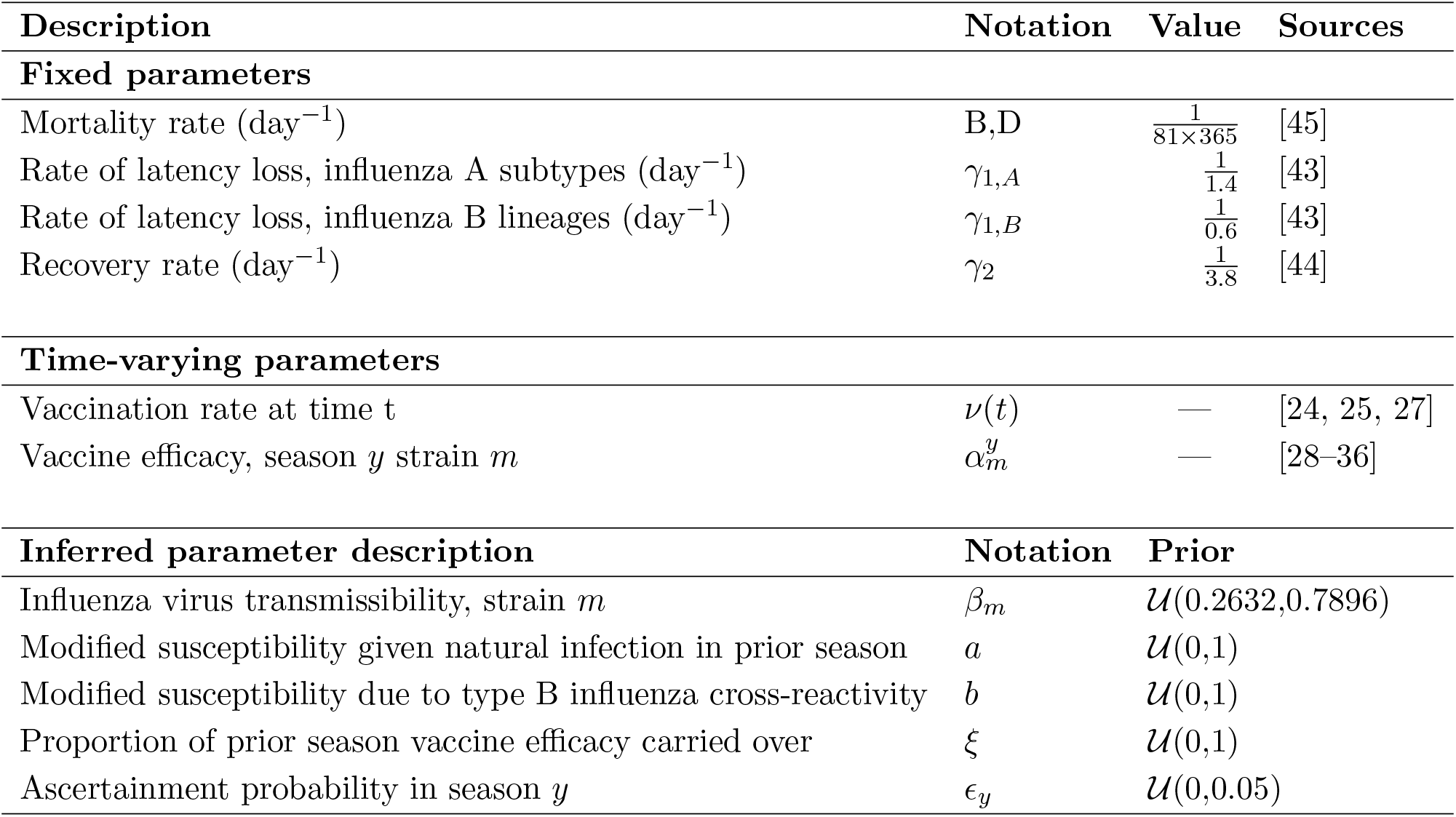
Overview of parameters in the model.

The ODE equations of the SEIR epidemiological model are given in Section 2.3 of the Supporting Information. The strain specific force of infection, *λ_m_*, satisfies

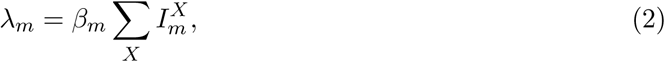

where *β_m_* is the transmission rate for strain *m*, implicitly comprising the contact rate and the transmissibility of the virus (the probability that a contact between an infectious person and a susceptible person leads to transmission), and the superscript *X* ∈ {*N, V*} denotes vaccination status (indexed by *N* for non-vaccinated, *V* for vaccinated).

For use in our observation model, we tracked the incidence *Z_m_*(*y*) of new strain *m* influenza infections in season *y* as a rate per 100,000 population:

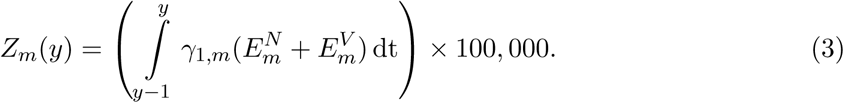

### Observation model

As discussed above, individuals consulting a GP for ILI do not necessarily directly correspond with individuals infected with one of the circulating influenza strains. To address this discrepancy, the final segment of the mathematical model linked the GP consultation data with the number of infections due to circulating influenza viruses in the population (Fig. 2, process D).

Our interest resided in the number of ascertainable cases, where we assumed that each individual infected by the strain of influenza under consideration had a probability *E* of being ascertainable, i.e. going to the GP, being recorded as having ILI, and having a detectable influenza viral load. We assumed the ascertainment probability was season-specific (but strain agnostic) to account for a range of influences such as climate or co-circulating infections. We obtained model estimates of the proportion of individuals experiencing cases of ascertainable influenza (of strain type *m*) in season *y* through scaling the incidence *Z_m_*(*y*) by the ascertainment probability in influenza season *y*, *ɛ_y_*. Consequently, the ascertainable influenza cases in season *y*, *Z_m_*^+^(*y*), obeys:

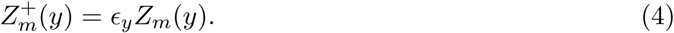

The parameter *ɛ_y_* is inferred by comparing the total predicted incidence in a given year to the calculated GP consultation rate (Eq. (1)).

### Parameter inference

To realise a model capable of generating influenza-attributed ILI GP consultations estimates that resemble the empirical data (Eq. (1)), we sought to fit the following collection of model parameters (Table 2): transmissibility of each influenza virus strain (*β_m_*), modified susceptibility to strain *m* given infection by a strain *m* type virus the previous season (*a*), carry over cross-reactivity protection between influenza B lineages (*b*), residual protection carried over from the prior season influenza vaccine (*ξ*), and an ascertainment probability per influenza season (*ɛ_y_*). One may notice the ascertainment probabilities were the sole group of parameters that could vary between influenza seasons, and we acknowledge that one may feasibly apply temporal (influenza season) dependencies to the transmission rates and immunity propagation parameters. Nevertheless, despite the the integration of additional parameters potentially improving the fit to the data, we would be running the risk of the model containing more parameters than can be justified by the data (overfitting). Instead, our intention was to extract a signal across the considered time frame, if present, identifying the most prominent transmission rates (comparing across the four influenza viruses) and immunity propagation mechanisms.

For inferring posterior parameter distributions, we employed an adaptive-population Monte Carlo approximate Bayesian computation (ABC) algorithm [46] combined with a local perturbation kernel (multivariate normal kernel with optimal local covariance matrix) [47]. Prior distributions were uniform for all parameters (Table 2).

Simulations began in the 2009/2010 influenza season and ran for a specified number of seasons with one parameter set. For the 2009/2010 influenza season, only the A(H1N1)pdm09 strain was present in our simulation (in accordance with the strain composition data where proportions of samples for the remaining three strains were very low, Fig. 1(a)), with the initial proportion of the population infected being 1 *×* 10*^−^*^5^ (one per 100,000). In all subsequent influenza seasons, with all four influenza virus strains being present, the initial proportion of the population infected by each strain was 2.5 *×* 10*^−^*^6^. We fit to the 2012/13 season onward, which omitted the influenza seasons (2009/10 and 2011/12) that had no influenza B lineage typing data (Fig. 1). This selected time frame contained a regime in which the dominant circulating influenza A subtype generally switched annually between H1N1(2009) and H3N2, while for type B influenza the Yamagata lineage incidence typically exceeded Victoria lineage incidence.

The summary statistics for our ABC procedure were comprised of two parts. The first component was a within-season temporal profile check; the peak in influenza infection (combining those in latent and infectious states) could not occur after February in any season. The second component defined the metric to measure the correspondence of the model predicted influenza-attributed ILI GP consultations versus the observed data. We worked with a metric akin to Poisson deviance. Compared to a sum of squared residuals error metric, a measure akin to Poisson deviance penalises with greater severity poor fits to data points of small magnitude. Accounting for season *y* and strain stratification *m*, we defined our deviance measure to be

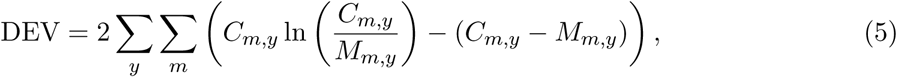

with *C_m,y_* the observed value for strain *m* in season *y*, and *M_m,y_* the model estimate for strain *m* in season *y*.

We amassed 10,000 particles representing a sample from the posterior distribution (see Section 3 of the Supporting Information for expanded details on the parameter estimation methodology).

### Forward simulations

We used the parameter set returning the lowest error (in the inference procedure) to explore potential strain dynamics up to the 2029/30 influenza season. Vaccination attributes (uptake and efficacy) for the 2009/10 to 2017/18 influenza seasons were taken from the observed data. For the forecasted seasons (2018/19 influenza season to the end of each simulation), we studied four scenarios arising from different hypotheses concerning future vaccine efficacy: (i) randomly sampled from all previous vaccine efficacy values (1,000 simulation replicates); (ii) ‘expected’ scenario (single simulation), strain-specific vaccine efficacies set at the median attained efficacy across 2010/11-2017/18 seasonal influenza vaccines (note, efficacy estimates from 2009/10 influenza season were not included as for 2009/10 we used pandemic influenza vaccine efficacy estimates); (iii) pessimistic scenario (single simulation), strain-specific vaccine efficacies set at the minimum attained efficacy across 2010/11-2017/18 seasonal influenza vaccines; and (iv) optimistic scenario (single simulation), strain-specific vaccine efficacies set at the maximum attained efficacy across 2010/11-2017/18 seasonal influenza vaccines. In addition, vaccine uptake in all projected influenza seasons matched that of the 2017/18 influenza season.

### Simulation and software specifics

All model simulations began at week 36 of the epidemiological season. Inference and model simulations were performed in Julia v0.6, with the system of ODEs solved numerically with the package DifferentialEquations v2.2.1. Data processing and production of figures were carried out in Matlab R2019a.

## Results

### Parameter inference

We invoked the adaptive-population Monte Carlo ABC algorithm, accumulating 10,000 parameter sets (representing a sample from the posterior distribution) to determine credible values of the model parameters (Table 3). Samples were obtained after completion of 600 generations of the adaptive-population Monte Carlo ABC scheme, at which point the generation-by-generation tolerance level updates were minor (Fig. 4(a)). Accordingly, additional simulations would only marginally change the posterior distributions. The model parameters were well defined, with Gaussian-shaped histograms (Fig. 4(b)). All parameter sets adhered to the mandatory temporal criteria of peak infection in each influenza season occurring prior to March (Figure S5).

**Fig. 4:**
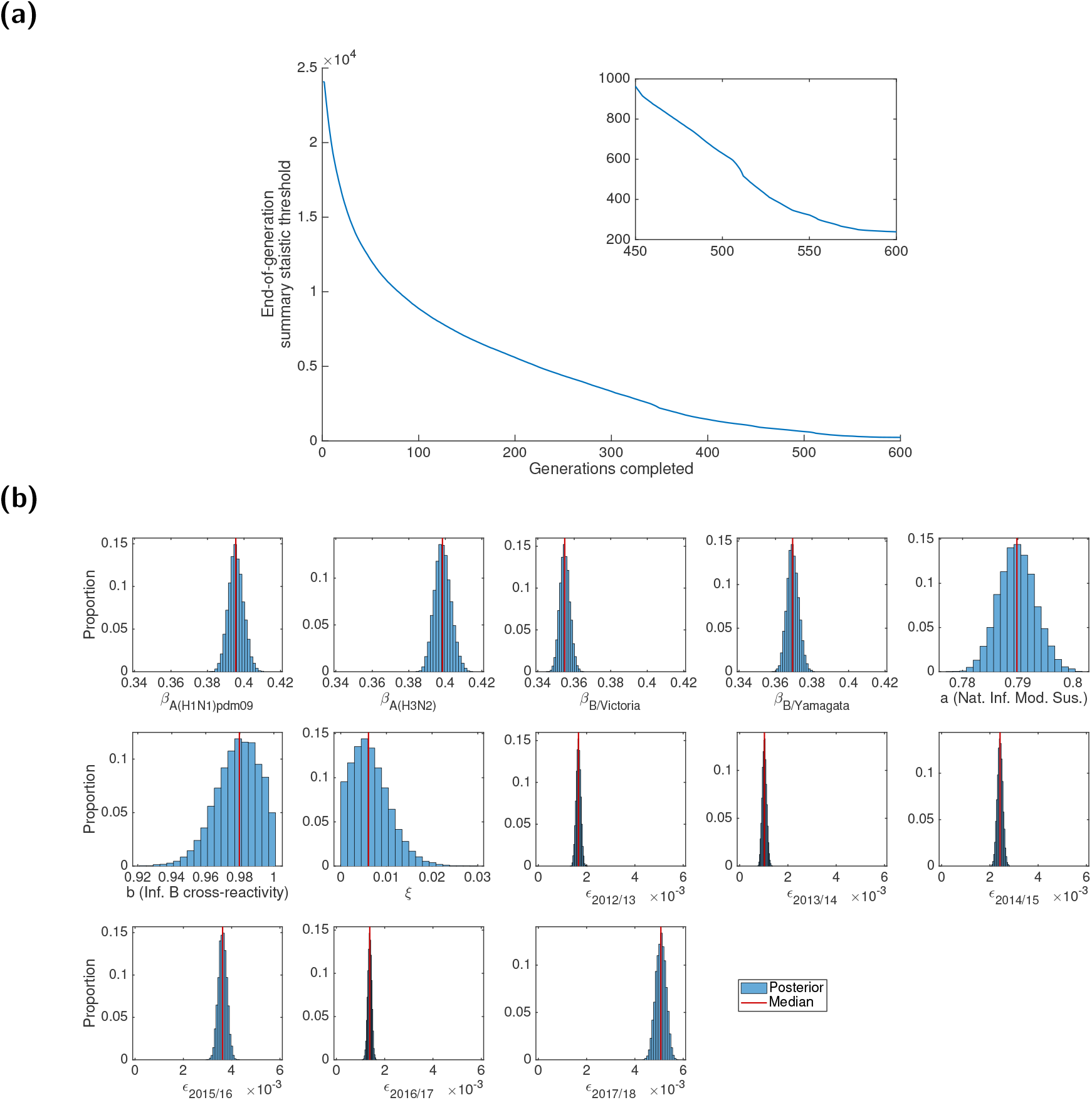
Results of the ABC scheme, fitting to the empirical data. **(a)** Summary metric threshold value upon completion of each generation of the inference scheme. The inset panel displays the latter quarter of generations. **(b)** Inferred parameter distributions estimated from 10,000 retained samples following completion of 600 generations of the inference scheme. Vertical red lines indicate the (non-weighted) median values for the model constants estimated from the inference procedure. Particularly noteworthy outcomes include: transmissibility of type A viruses exceeding type B viruses; prior season influenza B cross-reactivity and vaccine carry over had little impact on present season susceptibility; the highest ascertainment probability occurred in the 2017/18 influenza season.

**Table 3:**
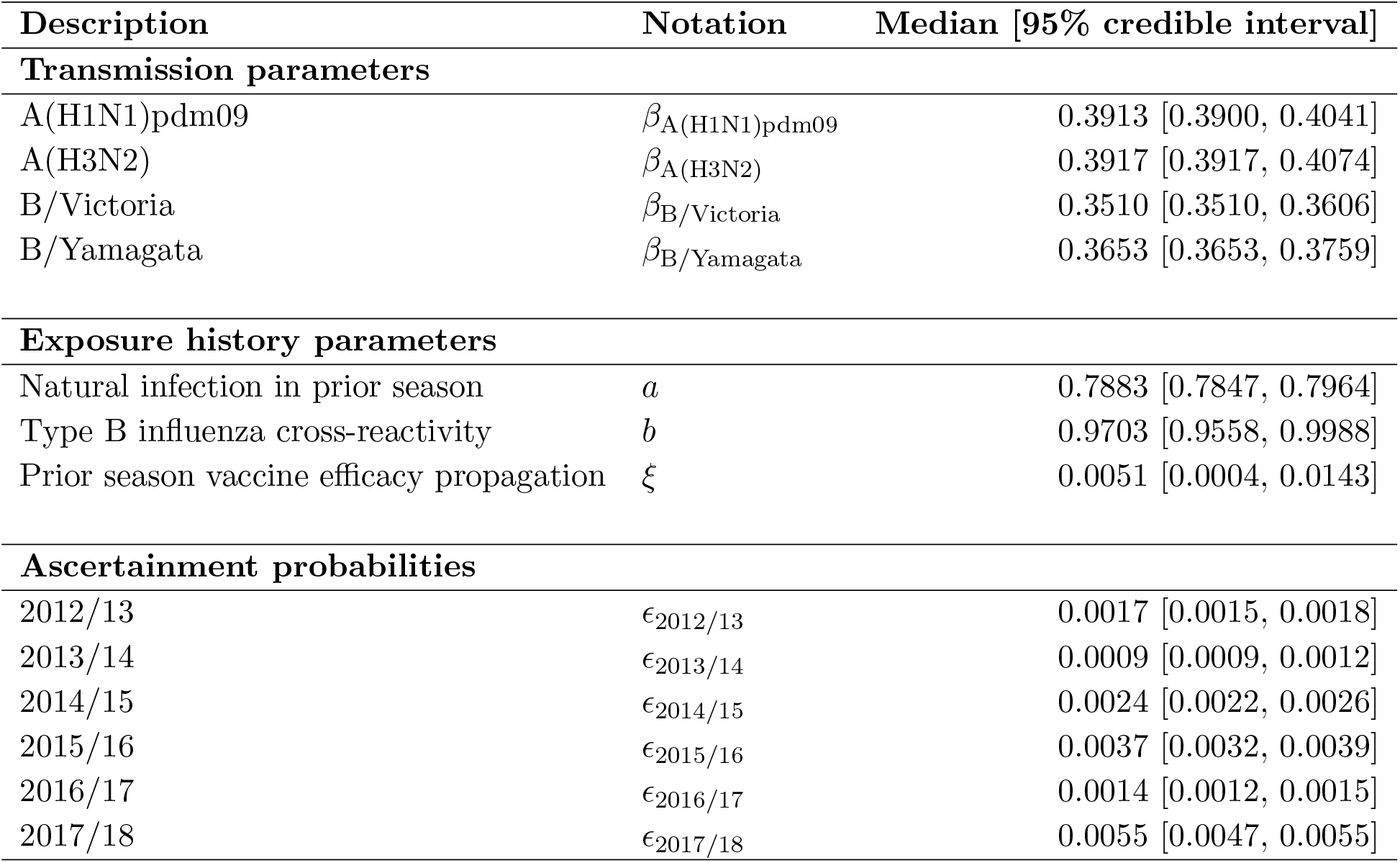
Values (posterior weighted quantiles) for the transmission, exposure history and ascertainment probability parameters inferred fitting to the empirical data. Numbers inside brackets indicate 95% credible intervals. All values are given to 4 d.p.

| Description | Notation | Median [95% credible interval] |
| --- | --- | --- |
| **Transmission parameters** |  |  |
| A(H1N1)pdm09 | $\beta_{A(H1N1)pdm09}$ | 0.3913 [0.3900, 0.4041] |
| A(H3N2) | $\beta_{A(H3N2)}$ | 0.3917 [0.3917, 0.4074] |
| B/Victoria | $\beta_{B/Victoria}$ | 0.3510 [0.3510, 0.3606] |
| B/Yamagata | $\beta_{B/Yamagata}$ | 0.3653 [0.3653, 0.3759] |
| **Exposure history parameters** |  |  |
| Natural infection in prior season | $a$ | 0.7883 [0.7847, 0.7964] |
| Type B influenza cross-reactivity | $b$ | 0.9703 [0.9558, 0.9988] |
| Prior season vaccine efficacy propagation | $\xi$ | 0.0051 [0.0004, 0.0143] |
| **Ascertainment probabilities** |  |  |
| 2012/13 | $\epsilon_{2012/13}$ | 0.0017 [0.0015, 0.0018] |
| 2013/14 | $\epsilon_{2013/14}$ | 0.0009 [0.0009, 0.0012] |
| 2014/15 | $\epsilon_{2014/15}$ | 0.0024 [0.0022, 0.0026] |
| 2015/16 | $\epsilon_{2015/16}$ | 0.0037 [0.0032, 0.0039] |
| 2016/17 | $\epsilon_{2016/17}$ | 0.0014 [0.0012, 0.0015] |
| 2017/18 | $\epsilon_{2017/18}$ | 0.0055 [0.0047, 0.0055] |

Inspecting the attained posterior distributions for virus transmissibility, estimates for the two influenza A subtypes are similar; both are larger than the corresponding estimates for the two type B lineages. However, the B/Yamagata transmissibility estimates exceeded those for B/Victoria, although the values are relatively close.

Amongst the three exposure history parameters (*a*, *b*, *ξ*), the parameter having a notable impact on the transmission dynamics was the modification to susceptibility due to natural infection in the previous influenza season (*a*). If infected by a given strain last season, the inferred weighted median estimate of 0.7883 corresponds to an approximate 21% reduction in susceptibility to the current season variant of that strain type. Strikingly, there was little carry over of prior season vaccine efficacy (*ξ*), with the majority of the posterior distribution massed near 0. Similarly, being previously infected by one of the type B influenza virus conferred little immunity against the remaining type B lineage virus in the next season.

Inferred ascertainment probabilities (*ε*), across all seasons, were typically within the range of 0.001-0.006. The highest ascertainment probabilities were found for the recent 2017/18 influenza season, attaining a median of 0.005; but this still suggests that only one in two hundred infections was reported to their GP and correctly identified as influenza.

We verified model robustness by performing parameter fits using two subsets of the historical influenza season data, spanning the time periods 2012/13-2015/16 and 2012/13-2016/17 respectively. Fitting to these differing influenza season ranges, outcomes generally exhibited qualitative consistency; in particular, amongst the three mechanisms for immunity propagation, current season susceptibility was modulated greatest by natural infection in the previous influenza season. The only notable quantitative differences when fitting to the two shortened time frames (compared to the distributions inferred when fitting to the complete time period) were elevated transmissibility levels (*β*), counteracted by an enlarged effect of prior infection (*a*) and vaccination propagation (*ξ*); further details are given in Section 4.2 of the Supporting Information.

In addition, we assessed the parameter identifiability competency of our ABC fitting scheme via generation of synthetic data from known parameters. Employing our inference scheme to fit the model to the synthetic data, we ably recovered the parameter values from which the synthetic data had been generated, giving extra confidence to our results (see Section 4.3 of the Supporting Information).

### Evaluating model fit

To assess the goodness-of-fit between our model and the available data, we performed 1,000 independent simulations using parameter sets drawn from the ABC inference procedure. Therefore, although the model is deterministic, we generated variability in epidemic composition due to the posterior distribution for the underlying parameters. The incidence of each of the four influenza types over each season was converted to attributed GP consultations (per 100,000), with both incidence and GP consultation outputs compared to the data (Fig. 5). We generally obtained a reasonable fit to the data, both in terms of the severity of infection each year (as controlled by the ascertainment *ϵ_y_*) but also in terms of type and subtype composition (which is largely driven by epidemiological dynamics and propagation of immunity).

**Fig. 5:**
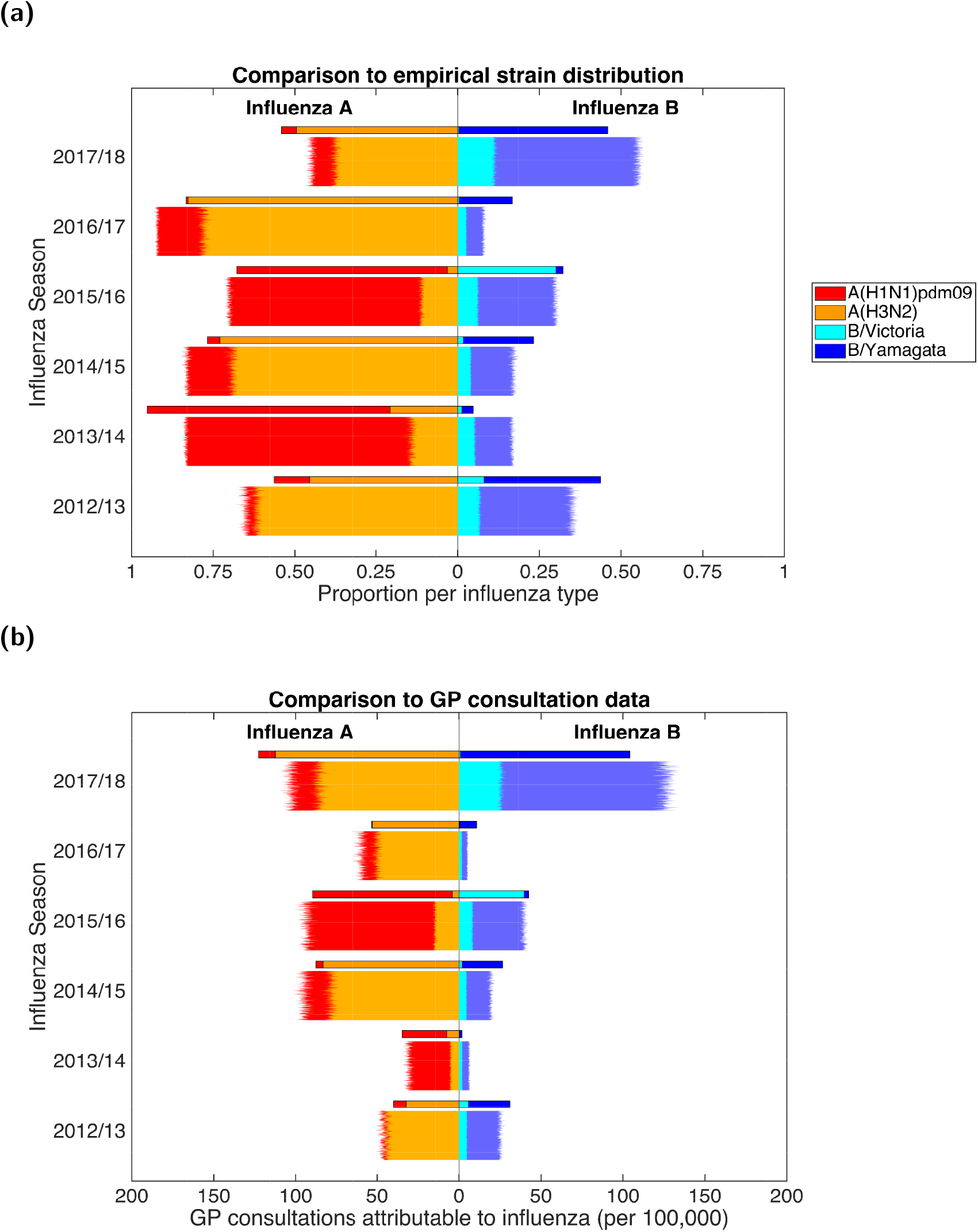
Posterior predictive distributions for strain composition and influenza positive GP consultations per 100,000 population. Stratified by influenza season, we present back-to-back stacked bars per simulation replicate, with 1,000 replicates performed, each using a distinct parameter set representing a sample from the posterior distribution. Each influenza season is topped out by a thicker stacked horizontal bar plot, corresponding to the strain-stratified point estimates for the empirical data. **(a)** Posterior predictive distributions for circulating influenza virus subtype/lineage composition. **(b)** Posterior predictive distributions for influenza positive GP consultations per 100,000 population. In both panels, the left side depicts data pertaining to type A influenza viruses (red shading denoting the A(H1N1)pdm09 subtype, orange shading the A(H3N2) subtype). In an equivalent manner, the right side stacked horizontal bars present similar data for type B influenza (cyan shading denoting the B/Victoria lineage, dark blue shading the B/Yamagata lineage). We see a reasonable qualitative model fit to the data, especially for the two influenza A subtypes.

Considering type A influenza, we clearly captured the alternating pattern of dominance by A(H3N2) and A(H1N1)pdm09 subtypes for the years 2012/13 to 2016/17; but more importantly the model was able to predict the unexpected result that A(H3N2) dominated in both 2016/17 and again in 2017/18. For influenza B, we obtained modest agreement for the overall magnitude of GP consultations as a result of type B, but this was not always in complete agreement with the subtype composition (Fig. 5(a)).

A more detailed comparison of predicted and observed subtypes in each season allowed us to analyse discrepancies in more detail (Fig. 6). The model predictions are shown with a violin plot to capture the variability in the distribution due to parameter uncertainty; while the observed data has confidence intervals (calculated from bootstrapping) due to the finite sample sizes. In general there was good agreement between the data and model, especially for years where particular subtypes dominated (e.g. A(H1N1)pdm09 in 2013/14 and 2015/16, A(H3N2) in 2014/15 and 2016/17, and B/Yamagata in 2012/13 and 2017/18). However, there were also discrepancies, most notably in 2015/16 where the majority of the B-influenza cases were predicted to the wrong lineage; a potential consequence of pursuing model parsimony, with preference for ensuring the generic behaviour for B/Yamagata across all other influenza seasons in the study period was correct, rather than seeking to capture every changing nuance when fitting to the data. It was also clear that the model generally performed less well when there were very low levels of a given subtype in a particular year.

**Fig. 6:**
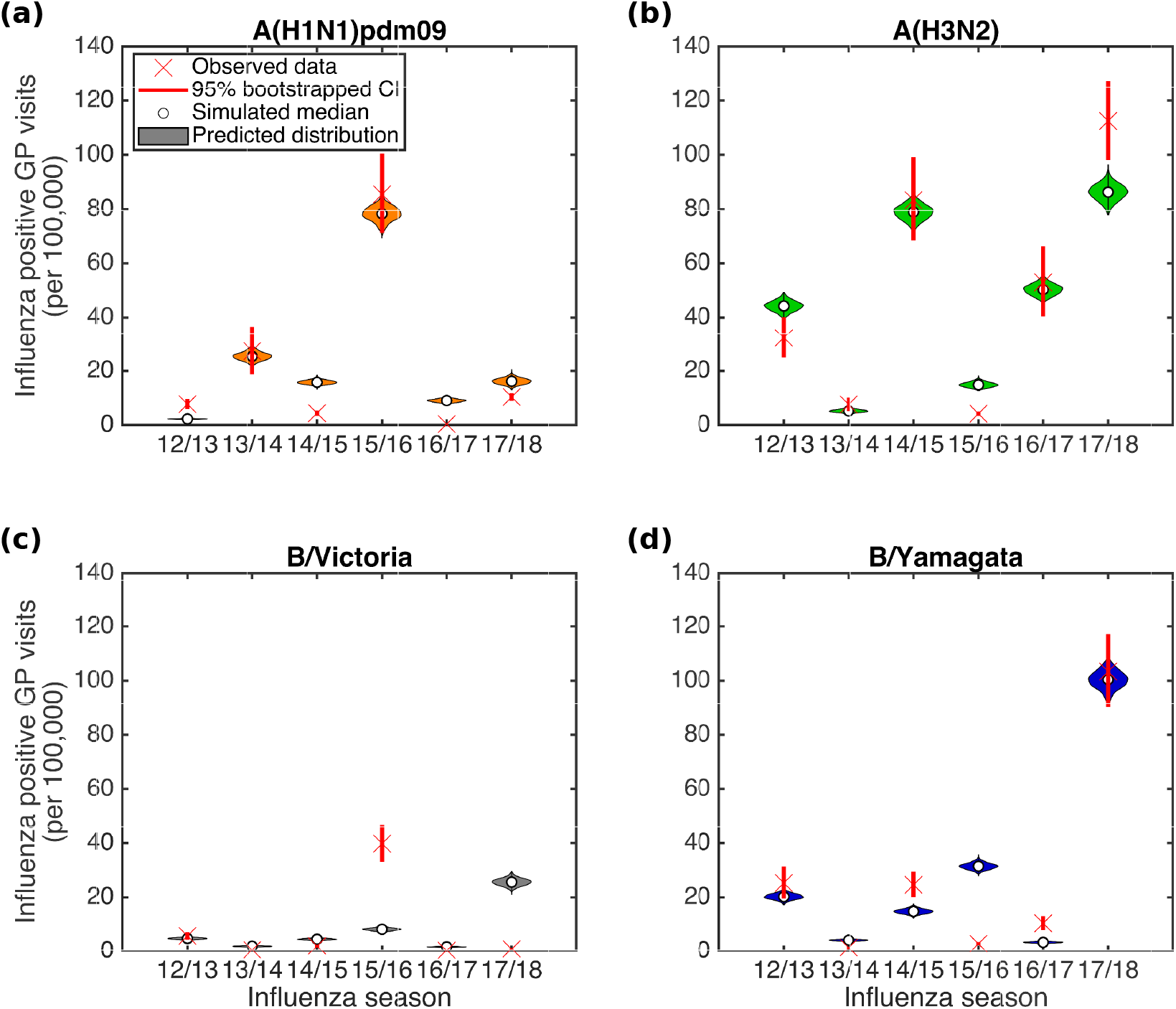
Posterior predictive influenza positive GP consultation distributions. We generated the estimated distributions from 1,000 model simulations, each using a distinct parameter set from the retained collection of particles. Crosses represent the observed data, with solid bars the range of the bootstrapped empirical data. Shaded violin plots denote outcomes from model simulations using the empirical data on vaccination, with filled circles the median value across the simulated replicates. **(a)** A(H1N1)pdm09; **(b)** A(H3N2); **(c)** B/Victoria; **(d)** B/Yamagata.

### Forward simulations

Using the parameter set from the inference scheme returning the closest correspondence with the data (lowest DEV value, Eq. (5)), we explored potential strain dynamics up to the 2029/30 influenza season under contrasting vaccine efficacy settings (Fig. 7).

**Fig. 7:**
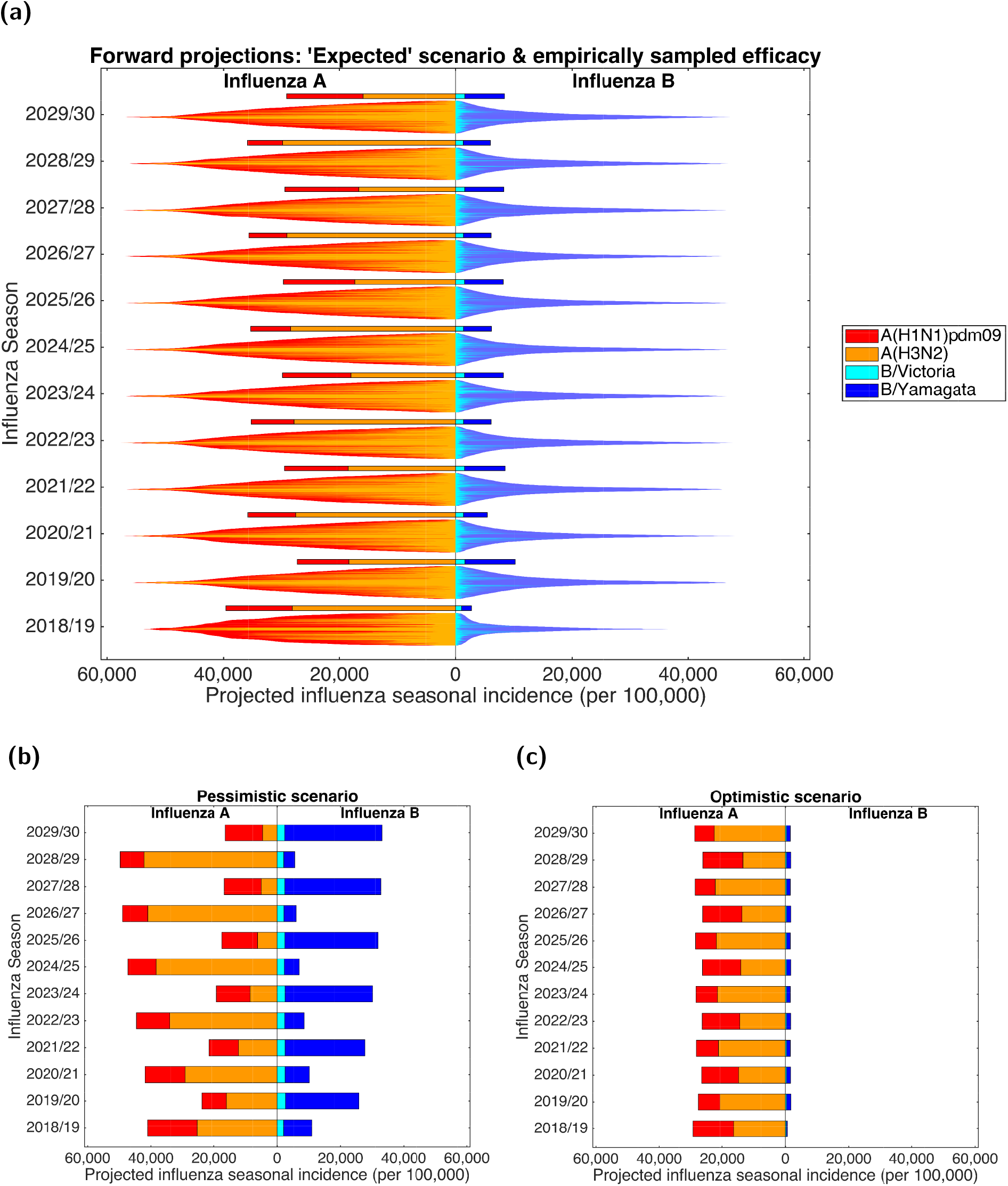
Projected influenza seasonal incidence (per 100,000 population) up to 2029/30. For all scenarios, in forward simulated seasons vaccine uptake matched that of the 2017/18 influenza season. In all panels, the left side depicts the seasonal incidence of type A influenza per 100,000 population (red shading denoting the A(H1N1)pdm09 subtype, orange shading the A(H3N2) subtype). The right side stacked horizontal bars present projected seasonal incidence of type B influenza per 100,000 population (cyan shading denoting the B/Victoria lineage, dark blue shading the B/Yamagata lineage).**(a)** Each influenza season is topped out by a thicker stacked horizontal bar plot, corresponding to the simulated estimate under the ‘expected’ vaccine efficacy scenario; efficacies by strain were as follows - A(H1N1)pdm09: 55.25%; A(H3N2): 30.45%; B/Victoria: 55.90%; B/Yamagata: 52.60%. Thin bars correspond to simulation runs where vaccine efficacy against each strain were randomly sampled from the empirical distribution (totalling 1,000 replicates). Simulations are arranged such that the largest epidemics are in the centre of the bar. **(b)** Projected incidence under a pessimistic vaccine efficacy scenario; efficacies by strain were as follows - A(H1N1)pdm09: 23.00%; A(H3N2): 0.00%; B/Victoria: 24.70%; B/Yamagata: 0.00%. **(c)** Projected incidence under an optimistic vaccine efficacy scenario; efficacies by strain were as follows - A(H1N1)pdm09: 73.00%; A(H3N2): 61.00%; B/Victoria: 92.00%; B/Yamagata: 92.00%.

When maintaining ‘expected’ vaccine efficacy (A(H1N1)pdm09: 55.25%; A(H3N2): 30.45% B/Victoria: 55.90%; B/Yamagata: 52.60%) across all future seasons (Fig. 7(a), single bars), we witness minor variability in predicted influenza incidence for both influenza types. Total influenza B incidence ranged between 2,700-10,250 cases per 100,000 each season, and seasonal influenza A incidence consistently reached 25,000-40,000 cases per 100,000. Furthermore, we predict periodic behaviour for the portion of influenza A viruses in circulation ascribed to the subtypes A(H1N1)pdm09 and A(H3N2), mirroring the observed pattern from 2012/13 to 2016/17. Influenza seasons ending in even numbered years (e.g. 2029/30) had a relatively even split between the two subtypes; whereas for influenza seasons ending in odd numbered years (e.g. 2028/29) the A(H3N2) subtype dominated the A(H1N1)pdm09, with influenza A viruses apportioned to A(H3N2) and A(H1N1)pdm09 subtypes being 70-85% and 15-30%, respectively.

An alternative approach is to use the single (best-fit) parameter set, but to sample the vaccine efficacies from historical estimates. The most striking factor was the amount of variability that comes as a direct results of uncertainty in the vaccine efficacies. As such all between-season patterns were swamped by the within-season variability, leading to distributions that were remarkably consistent between seasons. The only notable exception was 2018/19, where the definitive knowledge of the 2017/18 season had a clear impact of the predictions for 2018/19 leading to broadly higher levels of influenza A and lower levels of influenza B (Fig. 7(a)).

Comparing subtype distributions when vaccine efficacies were either sampled from historical estimates or fixed at ‘expected’ values, we found that under randomly sampled vaccine efficacies incidence of B/Yamagata tended to exceed B/Victoria in each season (holding in more than 70% of simulations, Figure S18). However, we obtained greater levels of variation for the influenza A subtypes. In an ‘expected’ vaccine efficacy setting (single bars), less than half of total influenza A cases in any season were attributed to the A(H1N1)pdm09 subtype (rather than A(H3N2)), whereas under variable vaccine efficacies (distributed bars) A(H1N1)pdm09 incidence was greater than A(H3N2) incidence in 30-50% of all simulations (Figure S18). Furthermore, in each future influenza season the majority of simulations (*∼*60-75%) performed with seasonally variable vaccine efficacies resulted in overall incidence rates exceeding the equivalent seasonal incidence estimates generated under ‘expected’ vaccine efficacy conditions.

The impact of vaccine efficay on projected incidence is further exemplified by the stark contrast in predictions when comparing the pessimistic scenario (A(H1N1)pdm09: 23.00%; A(H3N2): 0.00%; B/Victoria: 24.70%; B/Yamagata: 0.00%) to the optimistic scenario (A(H1N1)pdm09: 73.00%; A(H3N2): 61.00% B/Victoria: 92.00%; B/Yamagata: 92.00%). Under pessimistic vaccine efficacy conditions, both influenza types underwent vast fluctuations in incidence from one influenza season to the next. Type A was more severe in odd numbered years (e.g. 2028/29). Similarly, in even numbered years (e.g. 2029/30) type B incidence exceeded type A incidence. For the pessimistic vaccine efficacy scenario (compared to the other fixed vaccine efficacy scenarios) we also observe the strongest re-emergence of the alternating behaviour between dominant influenza A subtypes (Fig. 7(b)).

Maintaining a high vaccine efficacy, both influenza types had consistent incidence estimates. Whilst combined influenza B incidence was low, ranging approximately between 600-1700 cases per 100,000 each season, seasonal influenza A incidence frequently reached 25,000-30,000 cases per 100,000. Although we, once more, predict periodic behaviour for the portion of influenza A viruses in circulation ascribed to the subtypes A(H1N1)pdm09 and A(H3N2), under these optimistic vaccine efficacy conditions it was influenza seasons ending in odd numbered years having a relatively even split between the two subtypes; whereas for influenza seasons ending in even numbered years the A(H3N2) subtype dominated A(H1N1)pdm09 (Fig. 7(c)).

## Discussion

Understanding the processes that drive seasonal influenza transmission patterns is an important public-health question, given the economic cost and burden to human health afflicted by the condition [1, 2]. During recent years there has been marked progress in synthesising real-world data to inform mathematical model parameterisation. Yet, prior studies have tended to model each season and each strain circulating within that season independently [12, 14, 15]. Thus, the presence of strain and immunity interactions have been omitted. Using data subsequent to the 2009 pandemic for England, we have developed a dynamic multi-strain SEIR-type transmission model for seasonal influenza, explicitly incorporating immunity propagation mechanisms between seasons.

With a view to minimising the number of independent parameters, we fit a parsimonious mechanistic model to seasonal-level data on strain competition. In spite of the multi-strain complexity and time scale of the study period (six influenza seasons), predictions from the model attain a strong qualitative resemblance (in terms of strain composition and overall quantity of GP consultations) to the empirical subtype data from England. We attribute much of the discrepancies to the homogeneous way in which we have treated immunity propagation, as in practise this is driven by complex patterns of waning and cross immunity as well as somewhat irregular genetic drift.

Inferred transmissibility of type A influenza strains exceeding those of type B, which reflects type A being the predominant class of influenza virus in circulation over the studied time period [26]. Moreover, concentrating on the parameters relevant to immunity propagation, we uncover evidence against vaccination stimulating similar long-term immunity responses as for natural infection. These conclusions corroborate previous immunological studies showing that infection with influenza virus can induce broader and longer-lasting protection than vaccination [37, 48, 49], and authenticate prior work signalling that vaccine-mediated immunity rapidly wanes [50]. There are also indications that prior natural infection boosts vaccine responses against antigenically drifted strains, whereas prior vaccination does not [51]. The clinically observed impact of prior infection for enhancing vaccine efficacy was long-lasting, which may be used to instruct further model refinements.

In the case of finding minimal support for carry over cross-reactivity protection between influenza B lineages (parameter *b*), we have reported the best parameter fits under the modelling assumptions made. Considered collectively, the inferred parameter fits signify that, at the population level, a combination of transmissibility rates, season-specific ascertainment probabilities and modifications to susceptibility by lineage-specific immunity propagation (parameter *a*) and within-season vaccination are sufficient to obtain the closest correspondence to the data under the modelling assumptions, without requiring a prominent amount of influenza B cross-reactive immunity propagation. In other words, if there were presence of propagation of protection between influenza B lineages, it is dwarfed by other processes within the biological system. The lack of propagation of cross-reactivity protection between influenza B lineages may be unforeseen, as it has been documented that there is potential for viral interference between the influenza B virus lineages; infection with one influenza B virus lineage may be beneficial in protecting against subsequent infection with either influenza B virus lineage [39]. However, these experiments (using a ferret model of human influenza) had a short separation between primary and challenge virus infections (at most 28 days), whereas our timescale of interest is of the order of six months (spanning between peak infection in one influenza season to the initiation of the seasonal epidemic in the next).

The extent of immunity propagation stemming from earlier influenza infection and vaccination occurrences may conceivably be quantified through serological surveys, potentially the most direct and informative technique available to infer the dynamics of a population’s susceptibility and level of immunity [52]. There is scope to implement concepts monitoring changes in the immune response to infections in a rapid and cost-effective manner, by using existing primary care sentinel networks [53]. In this fashion, in England an age-stratified pilot serology study was recently initiated utilising the RCGP RSC to collect serological specimens and associated patient data to measure seropositivity and seroincidence due to seasonal influenza [54]. These progressive developments make usage of serological data to validate our study findings a realistic prospect.

Furthermore, though there have been studies carried out to quantify the strength of vaccine induced herd immunity for seasonal influenza [3, 4], the role of immunity originating from non-immunising exposure to influenza viruses in inducing herd immunity is unexplored. An upsurge in serology data would promote such assessments.

Ascertainment probabilities are key for linking population levels of influenza infection to GP-reported incidence. For greater parsimony we have assumed that ascertainment is independent of subtype, which could be an over-simplification in some years; but allowing ascertainment to depend on both season and subtype would over-fit the model. Across all considered influenza seasons, we found ascertainment probabilities were of a similar order (and concurred with the magnitudes of inferred ascertainment probabilities in [12]), with the greatest values obtained for the 2017/18 influenza season. The 2017/18 season saw moderate to high levels of influenza activity observed in the UK [24], exacerbated by a mismatch of the A(H3N2) component of the available vaccine towards the A(H3N2) strain in circulation [36]. The effects that consequential increased media coverage of an outbreak have on transmission dynamics can be complex [55], with the perception of the disease conceivably changing among the population [56]. Chiefly in this instance, for those infected and symptomatic the propensity to consult a GP may have been abnormally raised, which would result in an increased ascertainment probability relative to prior influenza seasons.

While historic immunity is predicted to play an important role in determining annual strain composition (leading to subtype patterns between seasons), our projections of the dynamics forward in time exemplify how variability in vaccine efficacy hampers our ability to make long-term predictions. Spanning our pessimistic to optimistic vaccine efficacy scenarios, we generally attained a model predicted incidence (per influenza season) of 30-50%. With volunteer challenge studies indicating that the majority of those infected by influenza are asymptomatic [57], plus only around 10% of those with ILI thought to consult a GP [58, 59], our model predicted influenza incidence pairs satisfactorily with previous influenza burden estimates covering England [60]. Additionally, influenza burden estimates in the USA (provided by the CDC) covering the 2017/18 and 2018/19 influenza seasons ascribe more than 48.8 million and 37.4-42.9 million influenza associated illnesses respectively, which is in the realms of 10-15% of the national population suffering symptomatic infection [61, 62].

Consistently high vaccine efficacies would aid forecasting efforts, providing additional benefits beyond the reduction in disease offered by new vaccines with improved clinical efficacy and effectiveness [63]. Most importantly, with annual emergence of drift variants of influenza viruses, cross-protective vaccines are needed to heighten protective effectiveness [64].

The development of the influenza transmission model presented here was built upon a collection of simplifying assumptions. A limitation of the model is fixing the influence of prior exposure history to a single season. Studies of repeated vaccination across multiple seasons suggest that vaccine effectiveness may be influenced by more than one prior season [65]. In an United Kingdom-centric analysis, Pebody *et al.* [34] scrutinised the possible effect of prior season vaccination on 2016/17 influenza season vaccine effectiveness; while there was no evidence that prior season vaccination significantly reduced the effectiveness of influenza vaccine during the current season in adults, it was deemed to increase effectiveness in children. Extending exposure history and susceptibility interaction functionality to encompass immunity propagation dependencies beyond one prior influenza season would assist efforts evaluating the potential effect of repeat influenza vaccinations in the presence of vaccine interference [66].

Second, on the basis of model parsimony, within the immunity propagation model component we limited the number of mechanisms linking prior influenza season exposure history to susceptibility in the subsequent influenza season. Though we did include a parameter corresponding to immunity proliferating across influenza B lineages between seasons, we did not include influenza A heterosubtypic immunity amongst the immunity propagation mechanisms. Recent work has found H1 and H3 influenza infection in humans induces neuraminidase-reactive antibodies displaying broad binding activity spanning the entire history of influenza A virus circulation (in humans) [37]; these developments motivate continued work and data acquisition to elicit the timespan over which cross-reactive antibodies (arising from natural influenza infection) may significantly reduce susceptibility to unrelated influenza A subtypes and influenza B lineages.

Third, we assumed the mechanism underpinning propagation of immunity from influenza vaccination in the previous season behaved linearly. We recognise a linear dependency is a strong generalisation. Given the vaccine-induced protection to an influenza virus strongly depends on the level of mismatch of the strain contained in the vaccine and the circulating strain, a vaccine that is very protective in the current influenza season might be ineffective the following year due to the influenza virus undergoing antigenic mutation. Thus, alternative non-linear formulations for the vaccine-derived immunity propagation process warrant consideration, if there is sufficiently detailed data to underpin the models.

Relatedly, to parameterise the amended strain-specific susceptibility values within exposure history groups encapsulating those both experiencing a natural infection event and undergoing vaccination in the prior influenza season, we assumed no interaction between residual vaccine and natural infection immunity (we simply enforced the dominating process). Other formulations governing the interaction between residual vaccine and natural infection immunity may be scrutinised, conditional on the emergence of relevant data to parameterise the process.

Fourth, for simplicity we assumed each season was initialised with a low level of all subtypes; in practise the subtypes seeding each epidemic will be contingent on the dynamics in the rest of the world. Finally, we assumed the value of the ascertainment probability to be constant over the course of the influenza season. The underlying assumption is that the quantities contributing towards ascertainment remain stable over time. Yet, in reality there are contributory factors that are likely to vary during the course of each seasonal outbreak; for example, the propensity to consult with a GP if inflicted with symptomatic illness.

We view the dynamic influenza transmission model presented here as a foundation stage for influenza modelling constructs incorporating strain and immunity interactions. Heterogeneity in social contact patterns and the role of age in influenza transmission potential and severity of health outcomes are important considerations, with children identified as the main spreaders of influenza infection [67, 68] and most influenza associated deaths occurring among elderly adults and those with co-morbid conditions that place them at increased risk [2, 69]. As a consequence, next steps should include augmenting population mixing patterns and age-structure into the mathematical framework. In addition, coupling the transmission model together with economic evaluation frameworks will permit cost-effectiveness appraisals of prospective vaccination programmes.

In summary, through the use of mathematical modelling and statistical inference, our analysis of seasonal influenza transmission dynamics in England quantifies the contribution of distinct immunity propagation mechanisms. We find that susceptibility in the next season to a given influenza strain type is modulated to the greatest extent through natural infection by that strain type in the current season, with residual vaccine immunity having a lesser role and inconsequential support for carry over type B cross-reactivity. As such, the approach utilised here offers a preliminary basis for long-term influenza modelling constructs incorporating strain and immunity interactions, although forecasts are strongly influenced by vaccine efficacy. We suggest that the adoption of influenza transmission modelling frameworks with immunity propagation provides a comprehensive manner to assess the impact of seasonal vaccination programmes.

## Supporting information

Manuscript tracked changes (V1 to V2)

## Acknowledgements

We thank Rachel Byford, Ana Correa, Chris McGee, Julian Sherlock and Sameera Pathiran-nehelage for their collective contribution towards the production of the RCGP RSC data extract utilised in this study. We thank Tom Irving and colleagues at Department of Health and Social Care for helpful discussions. We acknowledge Trystan Leng, Ben Atkins, Glen Guyver-Fletcher and Susie Cant for their constructive feedback on the manuscript. This report is independent research funded by the National Institute for Health Research (NIHR) (Policy Research Programme, Infectious Disease Dynamic Modelling in Health Protection, 027/0089). The views expressed are those of the authors and not necessarily those of the NIHR or the Department of Health and Social Care.

## Author contributions

**Conceptualisation:** Edward M. Hill, Matt J. Keeling, Stavros Petrou.

**Data curation:** Simon de Lusignan, Ivelina Yonova.

**Formal analysis:** Edward M. Hill.

**Funding acquisition:** Matt J. Keeling, Stavros Petrou.

**Investigation:** Edward M. Hill, Matt J. Keeling.

**Methodology:** Edward M. Hill, Matt J. Keeling.

**Software:** Edward M. Hill.

**Supervision:** Matt J. Keeling, Stavros Petrou.

**Validation:** Edward M. Hill, Matt J. Keeling.

**Visualisation:** Edward M. Hill, Matt J. Keeling.

**Writing - original draft:** Edward M. Hill.

**Writing - review & editing:** Edward M. Hill, Matt J. Keeling, Stavros Petrou, Simon de Lusignan, Ivelina Yonova.

## Data availability

The GP consultation data contain confidential information, with public data deposition non-permissible for socioeconomic reasons. The GP consultation data resides with the RCGP Research and Surveillance Centre and are available via the RCGP RSC website (www.rcgp.org.uk/rsc). All other raw data utilised in this study are publicly available; relevant references and data repositories are stated within the main manuscript and Supporting Information. Code and processed data used for the study is available at https://github.com/EdMHill/SeasonalFluImmunityPropagation.

## Competing interests

The authors declare that they have no competing interests.

## Supporting information items

### Supporting Information

**Supporting information for ‘Seasonal influenza: Modelling approaches to capture immunity propagation’.** This supplement consists of the following parts: (1) Data descriptions; (2) Complementary details of the modelling approach; (3) Parameter inference; (4) Additional results; (5) Model extension: Immunity propagation across multiple influenza seasons.

### Figure S1

Breakdown of subtype/lineage composition of sampled influenza viruses, per influenza type, from the United Kingdom in each influenza season (2009/10 onward). (a) Type A influenza, stratified by subtype; **(b)** Type B influenza, stratified by lineage.

### Figure S2

**Breakdown of subtype/lineage composition of sampled influenza viruses, as a proportion of all influenza samples, from the United Kingdom in each influenza season (2009/10 onward).** Sequenced samples confirmed as influenza positive, stratified by type and subtype/lineage. We assumed the fraction of undetermined samples ascribed to each subtype/lineage matched the observed proportions (for influenza seasons where strain-specific information were available).

### Figure S3

**All age, overall population vaccination uptake for 2009/10-2017/18 influenza seasons.** Vaccine uptake per influenza season were as follows (to 1 d.p.): 2009/10 - 21.0%; 2010/11 - 21.0%; 2011/12 - 21.7%; 2012/13 - 21.7%; 2013/14 - 23.0%; 2014/15 - 22.9%; 2015/16 - 22.9%; 2016/17 - 24.4%; 2017/18 - 26.0%.

### Figure S4

**Epidemiological model schematic.** Diagram of the compartmentalisation of the population based on an SEIR model, in conjunction with non-vaccinated (*N*) and vaccinated (*V*) statuses under a ‘leaky’ vaccination assumption. Epidemiological processes are represented by dashed lines. Vaccination processes are represented by solid lines. Demographic process (births and deaths) have been omitted. Compartments situated within the shaded region signify vaccinated groups, with vaccination occurring at rate */mu*. Transmission events lead to movement from a susceptible state (*S*) to latent (*E*,infected but not yet infectiousness). Latency is lost at rate *γ*_1*,m*_. Infected (*I*) transition to recovered (*R*) at rate *γ*_2_. Equivalent formulation for each strain *m*.

### Figure S5

**Example time series displaying proportion of population in infected states (latent or infectious).** Grey lines represent the simulated infected prevalence time series for 10 retained parameter sets from the inference procedure. With each simulation replicate starting on 1st September, red dashed vertical lines designate the beginning of March in each year. As stipulated by our parameter selection criteria, peak infection occurs prior to the 1st March.

### Figure S6

**Posterior predictive influenza positive GP visit distributions: 2012/13 influenza season.** Empirical data point estimate and bootstrap samples are depicted by the red diamond and green shaded regions respectively. Blue dots denote posterior predicted replicates. **(left)** Type A versus type B; **(top right)** influenza A subtypes, A(H1N1)pdm09 versus A(H3N2); **(bottom right)** influenza B lineages, B/Yamagata versus B/Victoria.

### Figure S7

**Posterior predictive influenza positive GP visit distributions: 2013/14 influenza season.** Empirical data point estimate and bootstrap samples are depicted by the red diamond and green shaded regions respectively. Blue dots denote posterior predicted replicates. **(left)** Type A versus type B; **(top right)** influenza A subtypes, A(H1N1)pdm09 versus A(H3N2); **(bottom right)** influenza B lineages, B/Yamagata versus B/Victoria.

### Figure S8

**Posterior predictive influenza positive GP visit distributions: 2014/15 influenza season.** Empirical data point estimate and bootstrap samples are depicted by the red diamond and green shaded regions respectively. Blue dots denote posterior predicted replicates. **(left)** Type A versus type B; **(top right)** influenza A subtypes, A(H1N1)pdm09 versus A(H3N2); **(bottom right)** influenza B lineages, B/Yamagata versus B/Victoria.

### Figure S9

**Posterior predictive influenza positive GP visit distributions: 2015/16 influenza season.** Empirical data point estimate and bootstrap samples are depicted by the red diamond and green shaded regions respectively. Blue dots denote posterior predicted replicates. **(left)** Type A versus type B; **(top right)** influenza A subtypes, A(H1N1)pdm09 versus A(H3N2); **(bottom right)** influenza B lineages, B/Yamagata versus B/Victoria.

### Figure S10

**Posterior predictive influenza positive GP visit distributions: 2016/17 influenza season.** Empirical data point estimate and bootstrap samples are depicted by the red diamond and green shaded regions respectively. Blue dots denote posterior predicted replicates. **(left)** Type A versus type B; **(top right)** influenza A subtypes, A(H1N1)pdm09 versus A(H3N2); **(bottom right)** influenza B lineages, B/Yamagata versus B/Victoria.

### Figure S11

**Posterior predictive influenza positive GP visit distributions: 2017/18 influenza season.** Empirical data point estimate and bootstrap samples are depicted by the red diamond and green shaded regions respectively. Blue dots denote posterior predicted replicates. **(left)** Type A versus type B; **(top right)** influenza A subtypes, A(H1N1)pdm09 versus A(H3N2); **(bottom right)** influenza B lineages, B/Yamagata versus B/Victoria.

### Figure S12

**Results of the ABC scheme, fitting to empirical data covering 2012/13-2015/16 influenza seasons (inclusive). (a)** Summary metric threshold value upon completion of each generation of the inference scheme. The inset panel displays the latter quarter of generations. **(b)** Inferred parameter distributions estimated from 10,000 retained samples following completion of 1,100 generations of the inference scheme. Vertical red lines indicate the (non-weighted) median values for the model constants estimated from the inference procedure.

### Figure S13

**Results of the ABC scheme, fitting to empirical data covering 2012/13-2016/17 influenza seasons (inclusive). (a)** Summary metric threshold value upon completion of each generation of the inference scheme. The inset panel displays the latter quarter of generations. **(b)** Inferred parameter distributions estimated from 10,000 retained samples following completion of 1,000 generations of the inference scheme. Vertical red lines indicate the (non-weighted) median values for the model constants estimated from the inference procedure.

### Figure S14

**Posterior predictive influenza positive GP consultation distributions, post-fitting to the empirical data covering the 2012/13-2015/16 influenza seasons (inclusive).** We generated the estimated distributions from 1,000 model simulations, each using a distinct parameter set from the retained collection of particles. **(a)** Back-to-back stacked bars per simulation replicate. Each influenza season is topped out by a thicker stacked horizontal bar plot, corresponding to the strain-stratified point estimates for the empirical data. **(b)** Comparison of model simulated outcomes (shaded violin plots, with filled circles corresponding to the median value across the simulated replicates) versus the observed data (crosses denote the point estimate, with solid bars the range of the bootstrapped empirical data).

### Figure S15

**Posterior predictive influenza positive GP consultation distributions, post-fitting to the empirical data covering the 2012/13-2016/17 influenza seasons (inclusive).** We generated the estimated distributions from 1,000 model simulations, each using a distinct parameter set from the retained collection of particles. **(a)** Back-to-back stacked bars per simulation replicate. Each influenza season is topped out by a thicker stacked horizontal bar plot, corresponding to the strain-stratified point estimates for the empirical data. **(b)** Comparison of model simulated outcomes (shaded violin plots, with filled circles corresponding to the median value across the simulated replicates)versus the observed data (crosses denote the point estimate, with solid bars the range of the bootstrapped empirical data).

### Figure S16

**Results of the ABC scheme, fitting to synthetic data. (a)** Summary metric threshold value upon completion of each generation of the inference scheme. The inset panel displays the latter quarter of generations. **(b)** Inferred parameter distributions estimated from 10,000 retained samples following completion of 1,000 generations of the inference scheme. Vertical red lines indicate the (non-weighted) median values for the model constants estimated from the inference procedure, and vertical black lines correspond to the true values of the parameters from which the data were generated. We ably recovered the parameter values from which the synthetic data had been generated.

### Figure S17

**Posterior predictive distributions for influenza positive GP consultations per 100,000 population, using the parameter sets generated when fitting to the synthetic data.** Stratified by influenza season, we present back-to-back stacked bars for 1,000 simulation replicates, each using a distinct parameter set representing a sample from the posterior distribution when fitting our mathematical model to the synthetic data. Each influenza season is topped by a thicker stacked horizontal bar plot, corresponding to the synthetic data values. The left side depicts the cumulative total of ILI GP consultations attributable to type A influenza per 100,000 population (red shading denoting the A(H1N1)pdm09 subtype, orange shading the A(H3N2) subtype). In an equivalent manner, the right side stacked horizontal bars present similar data for type B influenza (cyan shading denoting the B/Victoria lineage, dark blue shading the B/Yamagata lineage). Overall, simulated outcomes from each parameter set had strong agreement with the synthetic data.

### Figure S18

**Quantification of the projected dominating influenza A subtype and influenza B lineage by influenza season, up to 2029/30.** The left half depicts the proportion of simulations in which the major subtype contributor (*>*50% of cases) to the overall type A influenza incidence was A(H1N1)pdm09 (red bars) or A(H3N2) (orange bars). In an equivalent manner, the right half depicts the proportion of simulations in which the major lineage contributor (*>*50% of cases) to the overall type B influenza incidence was B/Victoria (cyan bars) or B/Yamagata (dark blue bars). Constructed from simulation runs where vaccine efficacy against each strain were randomly sampled from the empirical distribution (totalling 1,000 replicates). For all forward simulated seasons, vaccine uptake matched that of the 2017/18 influenza season.

### Figure S19

**Schematic showing the links between the vaccination, immunity propagation, epidemiological and observation model components for the extended model.** We adopt the visualisation conventions of [12], with ellipses indicating variables, and rectangles indicating data. Dotted arrows indicate relationships between prior season epidemiological outcomes and immunity propagation factors. Solid arrows indicate within-season processes. Circled capitalised letters indicate the relationships connecting the variables or data involved. These relationships are: process A, propagation of immunity as a result of exposure to influenza virus in the previous influenza season (through natural infection or vaccination); process B, modulation of current influenza season virus susceptibility; process C, estimation of influenza case load via the SEIR model of transmission; process D, ascertainment of cases through ILI recording at GP.

### Figure S20

**Mechanism for conferring immunity to influenza across multiple influenza seasons.** At the beginning of a generic influenza season *y*, those susceptible may be divided into two categories based on exposure history: (i) susceptibles in either the naive exposure history group, with unmodified susceptibility to infection, or in the vaccinated only exposure history group (labelled *S*); (ii) all those that are susceptible and in an exposure history group involving natural infection (labelled *Ŝ*). During the influenza season, those that were initially susceptible may be infected (state *I*). Of those remaining in state *Ŝ* at the end of season *y*, a proportion *δ* are retained in state *Ŝ* at the beginning of season *y* + 1. The remainder (1 - *δ*) transition to state *S*. Note, *δ* = 0 corresponds to no residual immunity being retained due to natural infection beyond one subsequent influenza season, while *δ* = 1 coincides with residual immunity being kept in all future seasons until the occurrence of a new infection event.

### Figure S21

**Results of the ABC scheme, fitting the extended model to the empirical data. (a)** Summary metric threshold value upon completion of each generation of the inference scheme. The inset panel displays the latter quarter of generations. **(b)** Inferred parameter distributions estimated from 10,000 retained samples following completion of 750 generations of the inference scheme. Vertical red lines indicate the (non-weighted) median values for the model constants estimated from the inference procedure. Particularly noteworthy outcomes include: transmissibility of type A viruses exceeding type B viruses; vaccine carry over had little impact on present season susceptibility; support for immunity stemming from natural infection being retained long term; the highest ascertainment probability occurred in the 2017/18 influenza season.

### Figure S22

**Posterior predictive influenza positive GP consultation distributions, post-fitting the extended model to the empirical data covering the 2012/13-2017/18 influenza seasons (inclusive).** We generated the estimated distributions from 1,000 model simulations, each using a distinct parameter set from the retained collection of particles. **(a)** Back-to-back stacked bars per simulation replicate. Each influenza season is topped out by a thicker stacked horizontal bar plot, corresponding to the strain-stratified point estimates for the empirical data. (**b**) Comparison of model simulated outcomes (shaded violin plots, with filled circles corresponding to the median value across the simulated replicates) versus the observed data (crosses denote the point estimate, with solid bars the range of the bootstrapped empirical data).

### Figure S23

**Results of the ABC scheme, fitting the extended model to synthetic data. (a)** Summary metric threshold value upon completion of each generation of the inference scheme. The inset panel displays the latter tenth of generations. **(b)** Inferred parameter distributions estimated from 10,000 retained samples following completion of 1,650 generations of the inference scheme. Vertical red lines indicate the (non-weighted) median values for the model constants estimated from the inference procedure, and vertical black lines correspond to the true values of the parameters from which the data were generated. We ably recovered the parameter values from which the synthetic data had been generated.

### Figure S24

**Posterior predictive distributions for influenza positive GP consultations per 100,000 population, using the parameter sets generated fitting the extended model to the synthetic data.** Stratified by influenza season, we present back-to-back stacked bars for 1,000 simulation replicates, each using a distinct parameter set representing a sample from the posterior distribution when fitting our mathematical model to the synthetic data. Each influenza season is topped by a thicker stacked horizontal bar plot, corresponding to the synthetic data values. The left side depicts the cumulative total of ILI GP consultations attributable to type A influenza per 100,000 population (red shading denoting the A(H1N1)pdm09 subtype, orange shading the A(H3N2) subtype). In an equivalent manner, the right side stacked horizontal bars present similar data for type B influenza (cyan shading denoting the B/Victoria lineage, dark blue shading the B/Yamagata lineage). Overall, simulated outcomes from each parameter set had strong agreement with the synthetic data.

### Table S1

**Adjusted vaccine effectiveness estimates for influenza by season and strain type.** 95% confidence intervals are stated within parentheses.

### Table S2

**Values (posterior weighted quantiles) for the transmission, exposure history and ascertainment probability parameters inferred fitting to 2012/13-2015/16, 2012/13-2016/17 and 2012/13-2017/18 influenza season data.** Numbers inside brackets indicate 95% credible intervals. All values are given to 4 d.p.

### Table S3

**Values (posterior weighted quantiles) for the transmission, exposure history and ascertainment probability parameters inferred fitting to synthetic data**. Numbers inside brackets indicate 95% credible intervals. All values are given to 4 d.p.

### Table S4

**Overview of parameters in the extended model.**

### Table S5

**Values (posterior weighted quantiles) for the transmission, exposure history and ascertainment probability parameters inferred fitting the extended model to the empirical data.** Numbers inside brackets indicate 95% credible intervals. All values are given to 4 d.p.

### Table S6

**Values (posterior weighted quantiles) for the transmission, exposure history and ascertainment probability parameters inferred fitting the extended model to synthetic data.** Numbers inside brackets indicate 95% credible intervals. All values are given to 4 d.p.

## Notes

#### Summary of Updates

Figure 6 revised (added labels to panels)

