## Supporting Information for "Seasonal influenza: Modelling approaches to capture immunity propagation"

### Supporting Information: Seasonal influenza: Modelling approaches to capture immunity propagation

### 1 Data descriptions

#### 1.1 Consultations in General Practices

The text contained in this subsection was provided by the Royal College of General Practitioners (RCGP). Thus, throughout this subsection "we" refers to contributors from RCGP who extracted the data.

**Method:**
Data were extracted from four RCGP Research and Surveillance Centre (RSC) databases [1]. UK general practice is a registration based system where all citizens can register with a single GP of their choice. Practices are computerised, and data entered into computerised medical record systems either as coded data [2], or free text. We extracted the coded data, and our results are based on this element of the record [3]. We extract all coded data, pseudonymising as close to sources as possible. Where patients have a range of codes inserted suggesting they opt out of record sharing we do not analyse their data [4].

The data sources was the Real World Evidence (RWE) database, established at University of Surrey in March 2015. This database contains all continuous historical data contained in the GP systems. Bulk data are extracted four times per year taking historic coded data. Where extracts fail for major reports we attempt to extract data using Morbidity Information and Export Syntax (MIQUEST). This is a Department of Health sponsored data extract tool.

RCGP provided weekly influenza-like-illness (ILI) rates for the RSC population from week 1 2000 to week 52 2018, disaggregated as follows:

1. **ISOYear**
2. **ISOWeek**
3. **Age**
4. **Chronic Disease**
5. **Population**
6. **Number of people with ILI**
7. **Rates per 100,000**

The data compared the ILI rates of people with chronic diseases (Chronic Disease = 1) against people without chronic diseases (Chronic Disease = 0).

**Contributors:**

1. **Simon de Lusignan** Director, guarantor for these data, assisted with clinical knowledge
2. **Rachel Byford** Put together structures for the data extraction.
3. **Ana Correa** Carried out the quality assessment of the output.
4. **Chris McGee** Carried out data extraction of ILI rates (covering week 1 2000 to week 52 2016).
5. **Julian Sherlock** Carried out data extraction of ILI rates and data quality checks (covering week 1 2017 to week 35 2018).

6. **Sameera Pathirannehelage** Carried out data extraction of ILI rates (covering week 1 2017 to week 52 2018).

7. **Ivelina Yonova** liaison with practices, and review of the original data request.

**Acknowledgements:**
Practices who have agreed to be part of the RCGP RSC and allow us to extract and used health data for surveillance and research. Filipa Ferreira (programme manager), and other members of the Clinical Informatics and Health Outcomes Research Group at University of Surrey. Apollo Medical Systems for data extraction. Collaboration with EMIS, TPP, In-Practice and Micro-test CMR supplier for facilitating data extraction. Colleagues at Public Health England.

#### 1.2 Respiratory Virus RCGP Surveillance

Data pertaining to the the weekly percentage of sentinel virology samples that were influenza positive were sourced from a subset of general practices in the RCGP Weekly Returns Service that submitted respiratory samples for virological testing from patients presenting in primary care with an ILI.

These data were displayed in figures within PHE annual influenza reports (for the 2009/10 to 2017/18 influenza seasons inclusive) [5, 6], and PHE Weekly National Influenza reports (for the 2013/14 influenza season onwards) [7].

PHE annual influenza reports were our baseline source for virological sample positivity data . Typically, curves displaying the data covered week 40 up to a variable week number dependent upon the season (week 13: 2016/2017; week 13: 2010/2011; week 15: 2011/2012,2014/2015, 2017/2018; week 20: 2013/2014). As an exception, the data curve for the entire 2009/2010 influenza season was available within the 2010/2011 PHE annual influenza report [5].

In all seasons, we assumed weeks 36-39 and weeks 21-39 had an influenza positivity of 0%.

To inform positivity values for weeks preceding week 20 for the 2013/14 influenza season onwards that were not illustrated within the PHE annual influenza report, we used PHE Weekly National Influenza reports [7], which provided UK GP sentinel swabbing scheme sample positivity on a weekly basis. Note that we gave precedence to figures explicitly stated within annual reports over positivity quantities within the weekly PHE influenza summaries.

#### 1.3 Circulating strain composition

Here, we expand on our use of FluNet [8] to inform the circulating influenza strain distribution per influenza season.

FluNet is a global web-based tool for influenza virological surveillance first launched in 1997. The virological data entered into FluNet, e.g. number of influenza viruses detected by subtype, are critical for tracking the movement of viruses globally and interpreting the epidemiological data.

FluNet reports weekly influenza surveillance data that are provided from over 140 National Influenza Centres of the Global Influenza Surveillance and Response System, national influenza reference laboratories, and WHO regional databases [9]. Per the WHO Global Epidemiological Surveillance Standards for Influenza guidance, influenza testing is conducted on specimens collected from persons presenting for medical care at participating surveillance sites who meet a clinical definition for influenza-like illness (defined as an acute respiratory infection with measured fever of $\geq 38°C$, and cough, with onset within the last 10 days), or severe acute respiratory infection (defined as an acute respiratory infection with history of fever or measured fever of $\geq 38°C$, and cough, with onset within the last 10 days, and requiring hospitalization) [9]. Influenza is confirmed by accepted laboratory diagnostic methods and actively reported by the reference laboratory to FluNet [10].

We computed the empirical circulating strain distribution in each influenza season (from the 2009/10 season onward) using data from FluNet for the United Kingdom. A salient feature of the influenza type data is the presence of samples whose exact subtype/lineage were not determined (Fig. S1). We assumed the fraction of undetermined samples ascribed to each subtype/lineage matched the proportions observed for the set of samples where strain-specific information were available (Fig. S2)

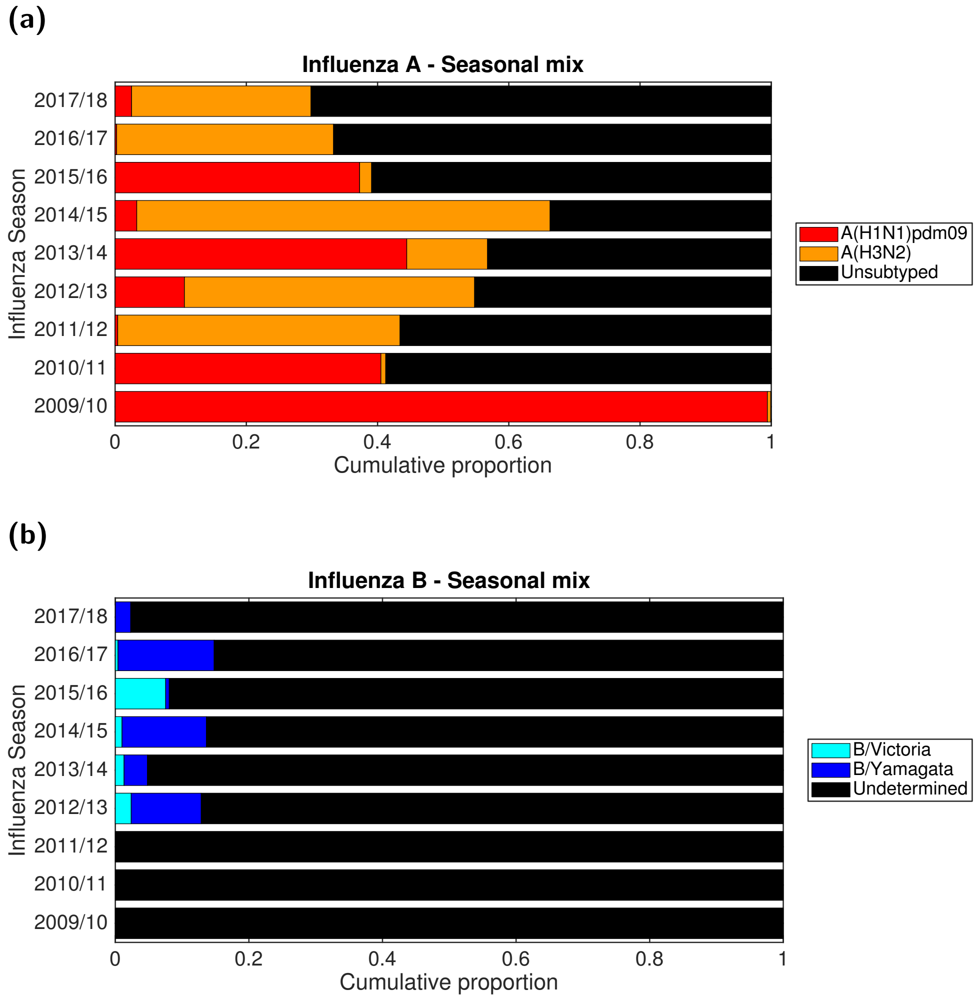

**Fig. S1: Breakdown of subtype/lineage composition of sampled influenza viruses, per influenza type, from the United Kingdom in each influenza season (2009/10 onward).** **(a)** Type A influenza, stratified by subtype; **(b)** Type B influenza, stratified by lineage.

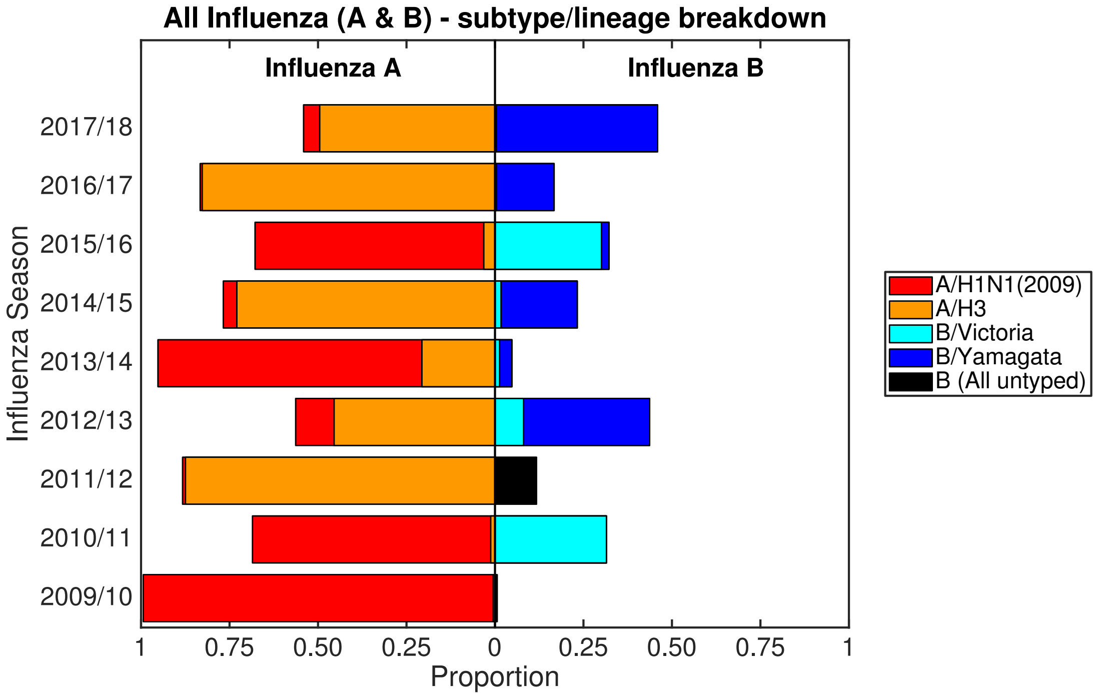

**Fig. S2: Breakdown of subtype/lineage composition of sampled influenza viruses, as a proportion of all influenza samples, from the United Kingdom in each influenza season (2009/10 onward).** Sequenced samples confirmed as influenza positive, stratified by type and subtype/lineage. We assumed the fraction of undetermined samples ascribed to each subtype/lineage matched the observed proportions (for influenza seasons where strain-specific information were available).

#### 1.4 Vaccine uptake

For the 2009/10 influenza season, we acquired vaccine uptake profiles from survey results on H1N1 vaccine uptake amongst patient groups in primary care [11].

For the 2010/11 influenza season onward, we sourced seasonal influenza vaccine uptake figures from Public Health England official statistics [6, 7]. These data comprised uptake profiles per target vaccination age group (e.g. at-risk under 65 years, 65 year and over, 2 year-olds, 3 year-olds) for each influenza season from PHE Weekly National Influenza Reports. Population-level measures were acquired through weighting each uptake curve by the the respective population proportion. Age-partitioned population distribution estimates for England were obtained from the Office of National Statistics (ONS) [12, 13].

All age, overall population influenza vaccine coverage has been on the rise, increasing from 21.0% to 26.0% between the 2009/10 and 2017/18 influenza seasons (Fig. S3).

We constructed daily uptake rates from either weekly or monthly uptake values via linear interpolation.

The temporal resolution of the empirical vaccine uptake data was dependent upon the age category, stratified by: (i) Non-school aged children (weekly coverage figures); (ii) monthly, school-aged children (monthly coverage figures).

**(i) Non-school aged children** Delivery of vaccination to those aged under 65 years in a clinical risk group or 65+ years old is through their GP. Additionally, all 2 and 3 year-olds eligible for influenza vaccination are administered the vaccine through their GPs. Uptake profiles are

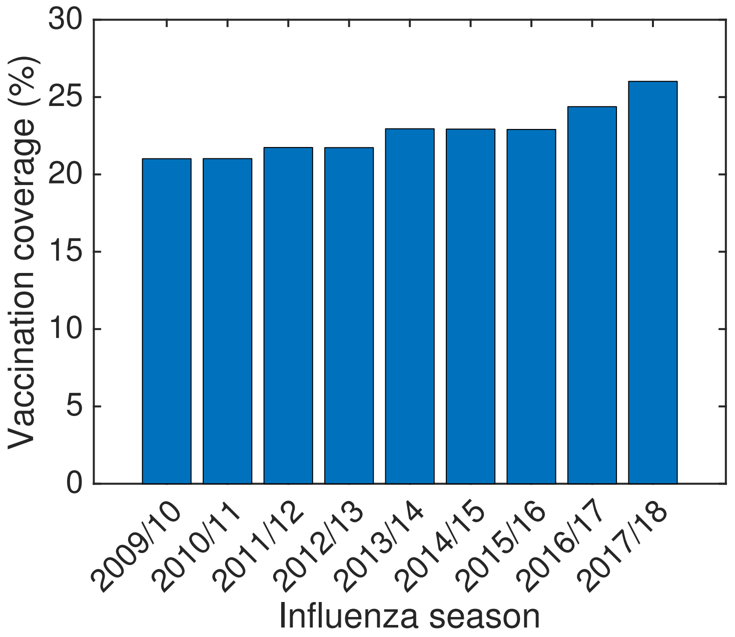

**Fig. S3: All age, overall population vaccination uptake for 2009/10-2017/18 influenza seasons.** Vaccine uptake per influenza season were as follows (to 1 d.p.): 2009/10 - 21.0%; 2010/11 - 21.0%; 2011/12 - 21.7%; 2012/13 - 21.7%; 2013/14 - 23.0%; 2014/15 - 22.9%; 2015/16 - 22.9%; 2016/17 - 24.4%; 2017/18 - 26.0%.

constructed based upon GP practices reporting weekly to Immform (the influenza vaccine uptake monitoring programme from Public Health England).

For the groupings of at-risk under 65 years and 65 years and above, vaccine uptake profiles were available from 2012/2013 influenza season onwards. For the 2010/2011 and 2011/2012 influenza seasons we invoked the uptake profile from the 2012/2013 influenza season.

Collection of cumulative weekly vaccine uptake for 2 year-olds and 3 year-olds started in 2013/2014.

**(ii) School-aged children** For school-aged children, monthly vaccination uptake estimates were available. These uptake estimates specified the proportion of children in England who received the influenza vaccine via school, pharmacy or GP practice.

#### 1.5 Vaccine efficacy

We informed population level vaccine efficacy estimates for each historical influenza season from publications detailing end-of-season age adjusted seasonal influenza vaccine effectiveness for adults and children in preventing laboratory-confirmed influenza in primary care in the United Kingdom [14–20]. In seasons where an equivalent publication were not available, we used mid-season or provisional end-of-season age adjusted seasonal vaccine efficacy estimates from Public Health England reports [21, 22].

The 2009/10 season was an exception. Due to the adjusted seasonal influenza vaccine efficacy being -30% (-89%, 11%) [14], we instead used the pandemic vaccine to inform efficacy against A(H1N1)pdm09, with the effectiveness against all other types set to zero.

The all age, strain-specific adjusted vaccine efficacy estimates gathered from the literature are displayed in Table S1. In the UK, since the incremental introduction of the universal childhood influenza vaccine programme began in the 2013/14 influenza season, an intra-nasally administered live attenuated influenza vaccine (LAIV) has been administered to children. Adult age classes have generally been offered an inactivated influenza vaccine (IIV). Therefore, due to using a population-averaged efficacy measure, in influenza seasons where IIV and LAIV formulations were administered to different age classes and/or risk groups the overall efficacy estimate is a combination of IIV and LAIV effectiveness.

With the empirical data not providing individual estimates for each influenza A subtype and influenza B lineage, we invoked a series of assumptions to produce the strain-specific vaccine efficacy quantities used within our study (Table 1).

We outline herein the set of enacted assumptions.

#### Approach to address absent influenza A efficacy data

##### Case one: Data absent for a single subtype only

Applicable to the following seasons: 2014/2015, 2015/2016, 2016/2017.

When there were estimates present for overall influenza A efficacy, but estimates for only one of the H1N1, H3N2 subtypes, for the subtype with no efficacy data we assumed its efficacy matched the overall influenza A efficacy estimate.

##### Case two: Data absent for all influenza A and one subtype

Applicable to the following seasons: 2010/2011,2011/2012.

In those seasons where there is an efficacy estimate for one of the two influenza A subtypes, but efficacy estimates for the other influenza A subtype and against influenza A overall are absent, we assumed the efficacy for the subtype with no relevant empirical data available was equal to that of the other influenza A subtype (for which there is an available estimate).

#### Approach to address absent influenza B efficacy data

##### Case one: Data absent for a single lineage only

Applicable to the following seasons: 2015/2016, 2016/2017.

When there were estimates present for overall influenza B efficacy, but estimates for only one of the B/Yam and B/Vic lineages, for the subtype with no efficacy data we assumed its efficacy matched the overall influenza B efficacy estimate.

##### Case two: Data absent for both lineages, but present for all influenza B

Applicable to the following seasons: 2011/2012, 2012/2013, 2013,2014, 2014/2015, 2017/2018.

In those seasons where there was an overall influenza B efficacy estimate, but efficacy estimates for both influenza B lineages were absent, we assumed the vaccine efficacy for both lineages matched the overall influenza B efficacy estimate.

**Table S1: Adjusted vaccine effectiveness estimates for influenza by season and strain type.** 95% confidence intervals are stated within parentheses.

| Season | Vaccine efficacy (95% CI) | | | | | | Source |
|---|---|---|---|---|---|---|---|
| | A(H1N1)pdm09 | A(H3N2) | All A | B/Yamagata | B/Victoria | All B | |
| 2009/10 | 72 (21, 90) | 0 | — | 0 | 0 | 0 | [14]* |
| 2010/11 | 56 (42, 66) | — | — | -34 (-448, 68)) | 78 (51, 91) | 57 (42,68) | [15] |
| 2011/12 | — | 23 (-10, 47) | — | — | — | 92 (38, 99) | [16] |
| 2012/13 | 73 (37, 89) | 26 (-4, 48) | 35 (11, 53) | — | — | 51 (34, 63) | [17] |
| 2013/14 | — | — | 61 (-8, 86) | — | — | 61 (-8, 86) | [21]† |
| 2014/15 | — | 29.3 (8.6, 45.3) | 29.9 (10.5, 45.1) | — | — | 46.3 (13.9, 66.5) | [18] |
| 2015/16 | 54.5 (41.6, 64.5) | — | 54.5 (41.6, 64.5) | — | 57.3 (28.4, 74.6) | 54.2 (33.1, 68.6) | [19] |
| 2016/17 | — | 31.6 (10.3, 47.8) | 36.7 (18.3, 51.0) | 58.5 (3.1, 82.2) | — | 54.5 (10.8, 76.8) | [20] |
| 2017/18 | 66.3 (33.4, 82.9) | -16.4 (-59.3, 14.9) | — | — | — | 24.7 (1.1, 42.7) | [22]‡ |

*: The adjusted seasonal influenza vaccine efficacy in the 2009/10 season was -30% (-89%, 11%) [14]. We therefore used the pandemic vaccine to inform efficacy against A(H1N1)pdm09, with the effectiveness against all other types set to zero.

†: Mid-season estimate of seasonal influenza vaccine effectiveness from [21]. Low incidence throughout the 2013/2014 influenza season meant reliable end-of-season estimates for the vaccine efficacy could not be attained (Personal communication, Public Health England).

‡: Provisional end-of-season influenza vaccine effectiveness results.

,

### 2 Complementary details of the modelling approach

#### 2.1 Epidemiological model specifics

Figure S4 depicts the epidemiological model, capturing within season vaccination and transmission dynamics.

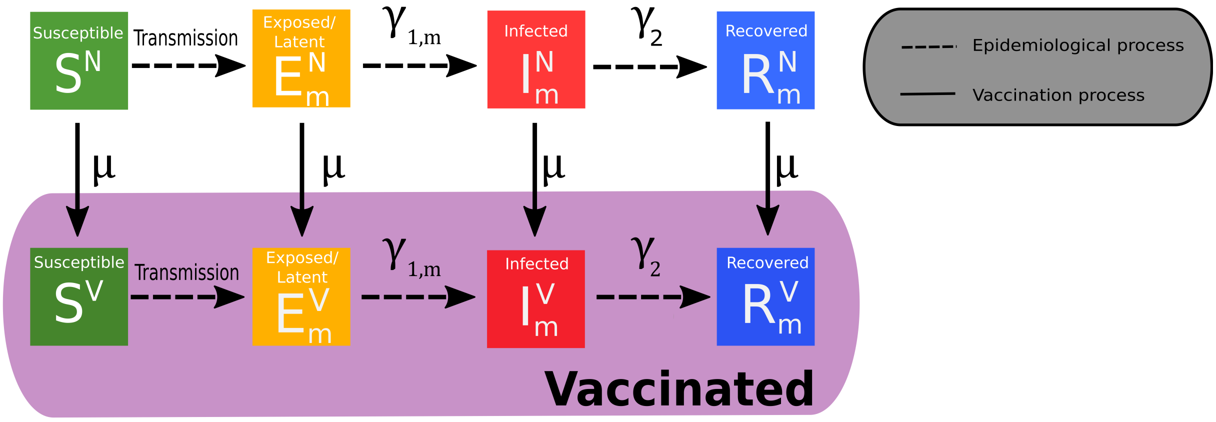

**Fig. S4: Epidemiological model schematic.** Diagram of the compartmentalisation of the population based on an SEIR model, in conjunction with non-vaccinated ($N$) and vaccinated ($V$) statuses under a "leaky" vaccination assumption. Epidemiological processes are represented by dashed lines. Vaccination processes are represented by solid lines. Demographic process (births and deaths) have been omitted. Compartments situated within the shaded region signify vaccinated groups, with vaccination occurring at rate $\mu$. Transmission events lead to movement from a susceptible state ($S$) to latent ($E$,infected but not yet infectiousness). Latency is lost at rate $\gamma_{1,m}$. Infected ($I$) transition to recovered ($R$) at rate $\gamma_2$. Equivalent formulation for each strain $m$.

The epidemiological model is a deterministic, non-age, multi-strain structured compartmental based model capturing demography, influenza infection status (with susceptible-latent-infected-recovered, SEIR, dynamics) and vaccine uptake. In addition, we assumed disease transmission to be frequency-dependent.

We let $E_m^X$, $I_m^X$ and $R_m^X$ denote the proportion of the population that have vaccination status $X \in \{N, V\}$ (indexed by $N$ for non-vaccinated, $V$ for vaccinated), and that are latent, infectious and recovered (as a result of natural infection) with respect to strain $m$.

Exposure to influenza virus in the previous influenza season, through natural infection or vaccination, modulated current influenza season susceptibility. Tracking immunity derived from natural infection and vaccination separately required ten distinct exposure history groups, with the susceptibility to strain $m$ for a given exposure history group $h$ encoded into the susceptibility array $f(h, m)$ (Fig. 3).

To incorporate exposure history from the previous influenza season, we let $S^{X,h}$ denote the proportion of the population that have (current influenza season) vaccination status $X \in \{N, V\}$ and exposure history $h$ that are susceptible to all strains. With ten exposure history groups $h$ in place, a total of 20 susceptibility states were used: ten $S^{N,h}$ states, accounting for susceptibles not vaccinated in the current influenza season stratified by exposure history grouping; ten $S^{V,h}$

states, tracking susceptibles who have been administered the vaccine in the current influenza season whilst retaining exposure history group information.

The epidemiological model has the following formulation, with time dependencies dropped:

$$\begin{cases} \frac{dS^N}{dt} &= B - \left(\sum_m \left(\sum_h f(h,m) S^{N,h}\right) \lambda_m\right) - \mu S^N - DS^N \\ \frac{dE^N_m}{dt} &= \left(\sum_h f(h,m) S^{N,h}\right) \lambda_m - \gamma_{1,m} E^N_m - \mu E^N_m - DE^N_m \\ \frac{dI^N_m}{dt} &= \gamma_{1,m} E^N_m - \gamma_2 I^N_m - \mu I^N_m - DI^N_m \\ \frac{dR^N_m}{dt} &= \gamma_2 I^N_m - \mu R^N_m - DR^N_m \\ \frac{dS^V}{dt} &= -\left(\sum_m \left(\sum_h f(h,m) S^{N,h}\right) (1-\alpha_m)\lambda_m\right) + \mu S^N - DS^V \\ \frac{dE^V_m}{dt} &= \left(\sum_h f(h,m) S^{V,h}\right) (1-\alpha_m)\lambda_m - \gamma_{1,m} E^V_m + \mu E^N_m - DE^V_m \\ \frac{dI^V_m}{dt} &= \gamma_{1,m} E^V_m - \gamma_2 I^V_m + \mu I^N_m - DI^V_m \\ \frac{dR^V_m}{dt} &= \gamma_2 I^V_m + \mu R^N_m - DR^V_m \end{cases} \tag{1}$$

where $\gamma_{1,m}$ is the strain-dependent rate of loss of latency, $\gamma_2$ the rate of loss of infectiousness, $\alpha_m$ the strain-specific vaccine efficacy, and $\mu = \frac{\nu}{S^N + \sum\limits_m (E^N_m + I^N_m + R^N_m)}$ with $\nu$ the rate of vaccination in the population.

To maintain a constant population size, birth and death rates were equal ($B = D$). We set the mortality rate based upon a gender-averaged life expectancy for newborns in England, computed from male and female specific statistics from ONS data [23]. We computed the gender-averaged life expectancy for newborns in England to be approximately 81 years, with the corresponding daily mortality rate being $D = \frac{1}{81 \times 365}$.

The strain specific force of infection, $\lambda_m$, satisfies $\lambda_m = \beta_m \sum\limits_X I^X_m$, where $\beta_m$ is the transmission rate for strain $m$, implicitly comprising the contact rate and the transmissibility of the virus (the probability that a contact between an infectious person and a susceptible person leads to transmission).

#### 2.2 Between season exposure history group mappings

At the beginning of each influenza season (1st September), we enacted the following exposure history group assignments:

- $S^N \to S^{N,h=\bar{N}}$
- $S^V \to S^{N,h=\bar{V}}$
- $\{E^{N,m}, I^{N,m}, R^{N,m}\} \to S^{N,h=m}$
- $\{E^{V,m}, I^{V,m}, R^{V,m}\} \to S^{N,h=m\&\bar{V}}$

where $X$ represents vaccination status, with $N$ corresponding to non-vaccinated and $V$ vaccinated (bars on vaccination state symbols relate to vaccination status in the previous influenza season).

### 3 Parameter inference

The aim of our parameter estimation was to select parameter sets generating influenza-attributed ILI GP consultation model predictions that best resemble the empirical data.

In full, we sought to estimate: transmissibility of each influenza virus strain ($q_m$), modified susceptibility to strain $m$ given infection by a strain $m$ type virus the previous season ($a$), carry over cross-reactivity protection between influenza B lineages ($b$), proportion of prior season vaccine efficacy propagated to the current season ($\xi$), and an ascertainment probability per influenza season ($\epsilon_t$).

We selected an Approximate Bayesian Computation (ABC) approach, where results are first simulated with a parameter set, after which it is rejected or accepted based on a divergence measure or statistic.

We sought parameter sets that: (i) adhered to our infection prevalence temporal profile check; the peak in influenza infection could not occur after February (i.e. during March through August inclusive) in any season; (ii) minimised our deviance statistic, measuring the correspondence of the model predicted influenza-attributed ILI GP consultations versus the observed data (Equation 1).

To update the parameter sets we employed an adaptive population Monte Carlo ABC algorithm [24], an ABC scheme combined with particle Monte Carlo methods to minimise the number of simulations.

Outlining the inference scheme in brief, we initially drew 10,000 particles (i.e. 10,000 parameter sets) from the prior distributions, simulated and calculated their statistics. From these first-generation particles, each with equal weight, we generated 10,000 additional particles. Next, we sampled new particles according to the weight of the first generation particles and perturbed according to a multivariate normal distribution with twice the weighted variance-covariance matrix (computed using analytic weights) of the first generation particles. After simulation of the new particles, we computed their statistics and new weights. Of the now 20,000 particles, we retained the 10,000 particles with the lowest statistics.

The procedure was repeated iteratively, except all subsequent perturbations were carried out with a local perturbation kernel. Local perturbation kernels assign a different covariance matrix for each particle of the previous population. The class of perturbation kernel we selected was the multivariate normal kernel with optimal local covariance matrix (OLCM) [25]. We ran the process for at least 600 generations.

The final generation of 10,000 particles represents a sample from the posterior distribution. To compute posterior parameter distribution summary statistics, parameter values were sorted in ascending order, $x_1, \ldots, x_{10,000}$, with associated normalised weights $\bar{w}_1, \ldots, \bar{w}_{10,000}$ (sum of all weights equal to one). The weighted median was defined as the element $x_k$ such that:

$$\sum_{i=1}^{k-1} \bar{w}_i \leq 0.5 \qquad \text{and} \qquad \sum_{i=k+1}^{n} \bar{w}_i \leq 0.5.$$

The 95% credible intervals were calculated as

$$Q_{2.5}(x) = \inf \left\{ x_k : \sum_{i=0}^{k} \bar{w}_i \leq 0.025 \right\}$$
$$Q_{97.5}(x) = \sup \left\{ x_k : \sum_{i=0}^{k} \bar{w}_i \geq 0.975 \right\}.$$

We additionally utilised the retained particles for counterfactual and forward projection.

### 4 Additional results

#### 4.1 Fitting to the empirical data (2012/13-2017/18 influenza seasons)

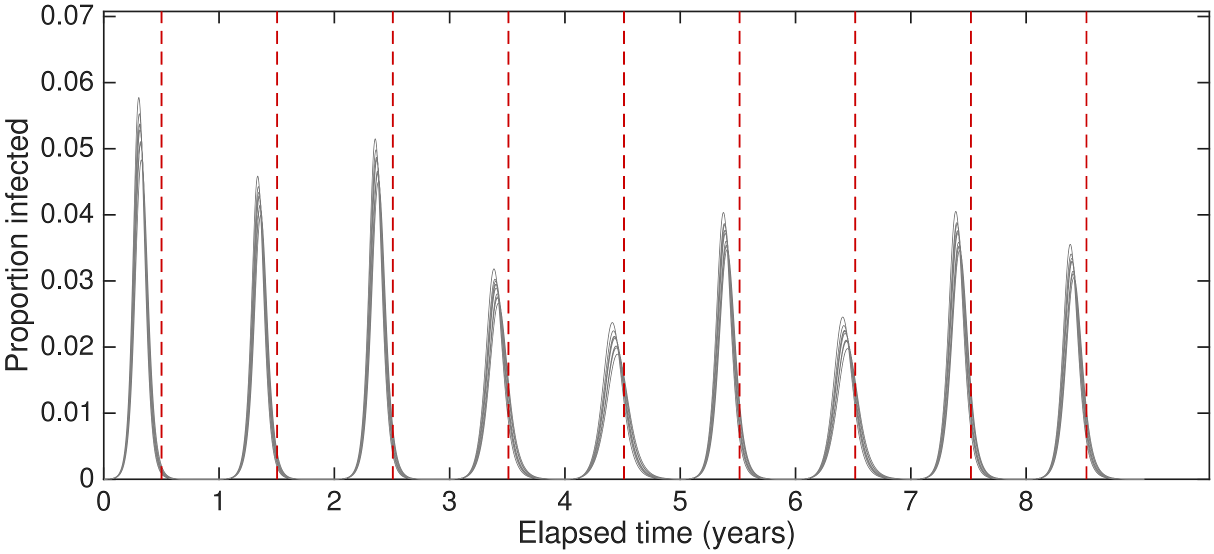

**Fig. S5: Example time series displaying proportion of population in infected states (latent or infectious).** Grey lines represent the simulated infected prevalence time series for 10 retained parameter sets from the inference procedure. With each simulation replicate starting on 1st September, red dashed vertical lines designate the beginning of March in each year. As stipulated by our parameter selection criteria, peak infection occurs prior to the 1st March.

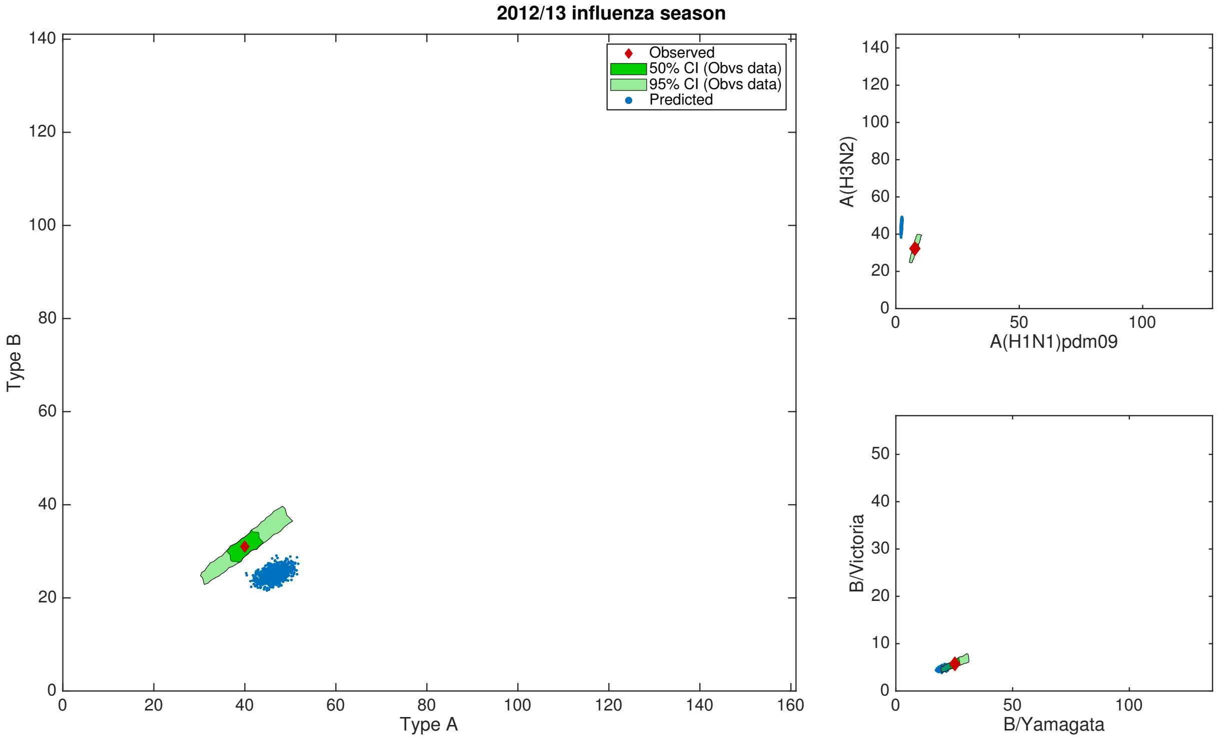

**Fig. S6: Posterior predictive influenza positive GP visit distributions: 2012/13 influenza season.** Empirical data point estimate and bootstrap samples are depicted by the red diamond and green shaded regions respectively. Blue dots denote posterior predicted replicates. **(left)** Type A versus type B; **(top right)** influenza A subtypes, A(H1N1)pdm09 versus A(H3N2); **(bottom right)** influenza B lineages, B/Yamagata versus B/Victoria.

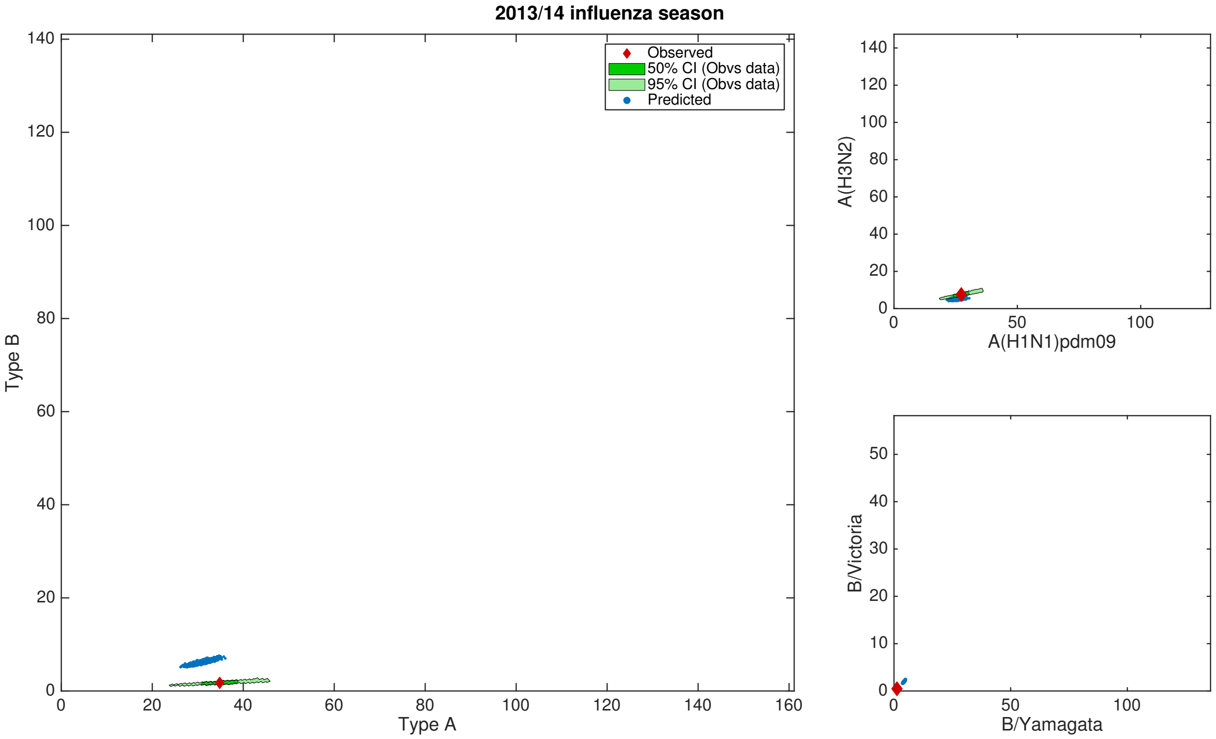

**Fig. S7: Posterior predictive influenza positive GP visit distributions: 2013/14 influenza season.** Empirical data point estimate and bootstrap samples are depicted by the red diamond and green shaded regions respectively. Blue dots denote posterior predicted replicates. **(left)** Type A versus type B; **(top right)** influenza A subtypes, A(H1N1)pdm09 versus A(H3N2); **(bottom right)** influenza B lineages, B/Yamagata versus B/Victoria.

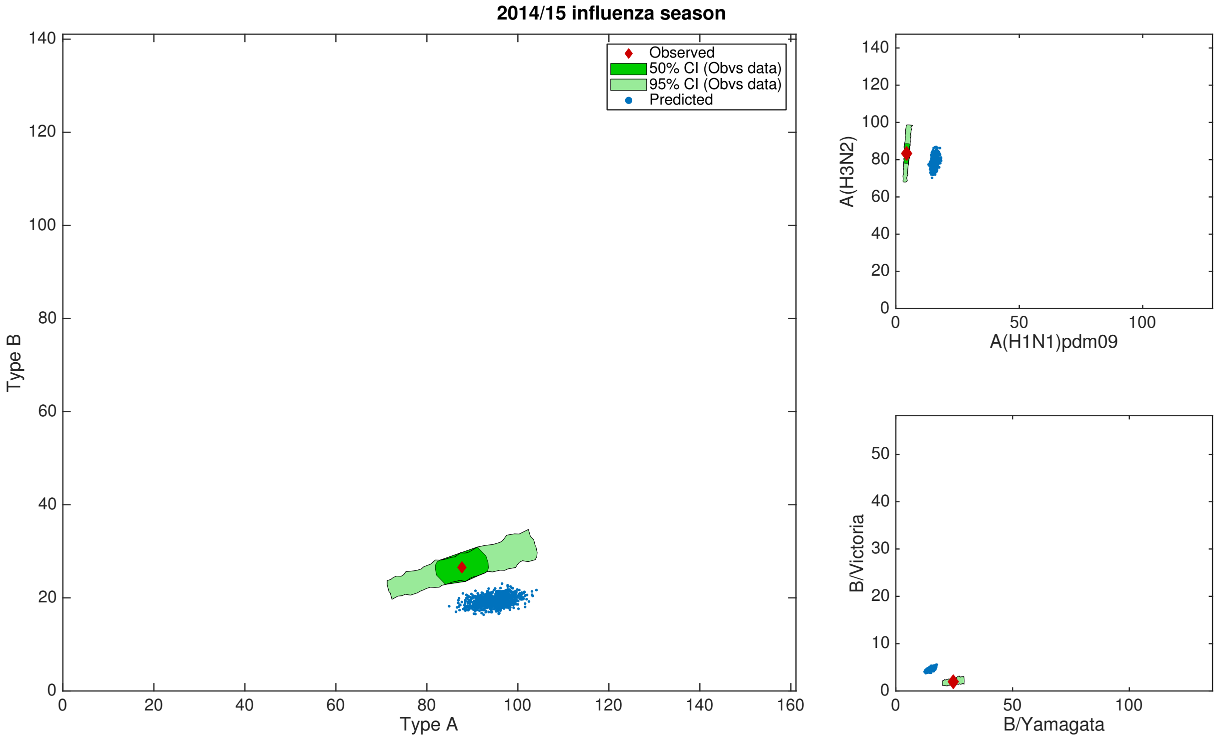

**Fig. S8: Posterior predictive influenza positive GP visit distributions: 2014/15 influenza season.** Empirical data point estimate and bootstrap samples are depicted by the red diamond and green shaded regions respectively. Blue dots denote posterior predicted replicates. **(left)** Type A versus type B; **(top right)** influenza A subtypes, A(H1N1)pdm09 versus A(H3N2); **(bottom right)** influenza B lineages, B/Yamagata versus B/Victoria.

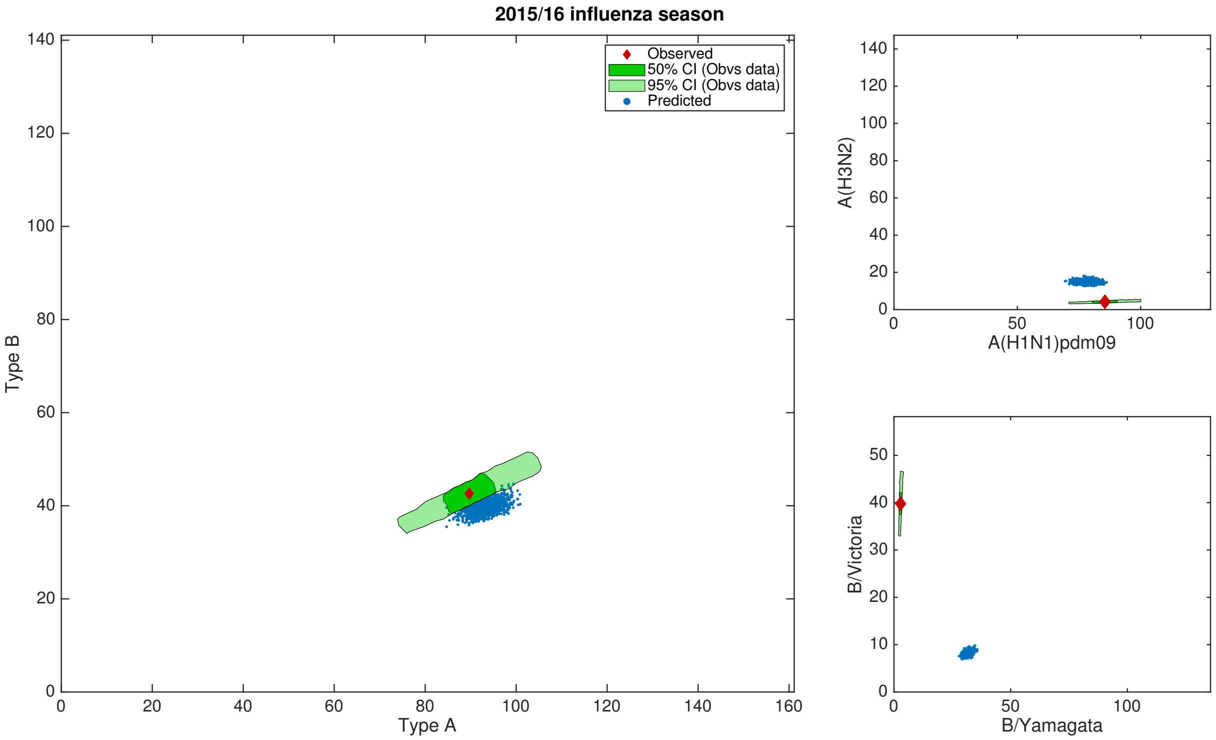

**Fig. S9: Posterior predictive influenza positive GP visit distributions: 2015/16 influenza season.** Empirical data point estimate and bootstrap samples are depicted by the red diamond and green shaded regions respectively. Blue dots denote posterior predicted replicates. **(left)** Type A versus type B; **(top right)** influenza A subtypes, A(H1N1)pdm09 versus A(H3N2); **(bottom right)** influenza B lineages, B/Yamagata versus B/Victoria.

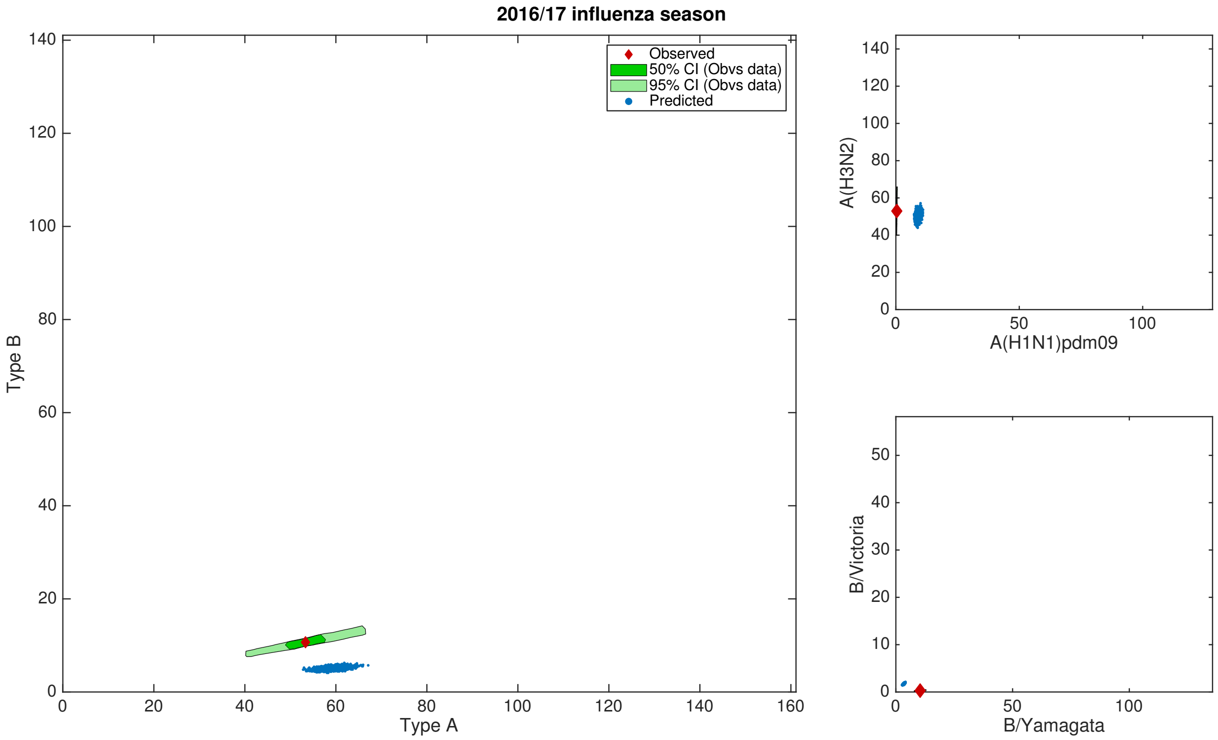

**Fig. S10: Posterior predictive influenza positive GP visit distributions: 2016/17 influenza season.** Empirical data point estimate and bootstrap samples are depicted by the red diamond and green shaded regions respectively. Blue dots denote posterior predicted replicates. **(left)** Type A versus type B; **(top right)** influenza A subtypes, A(H1N1)pdm09 versus A(H3N2); **(bottom right)** influenza B lineages, B/Yamagata versus B/Victoria.

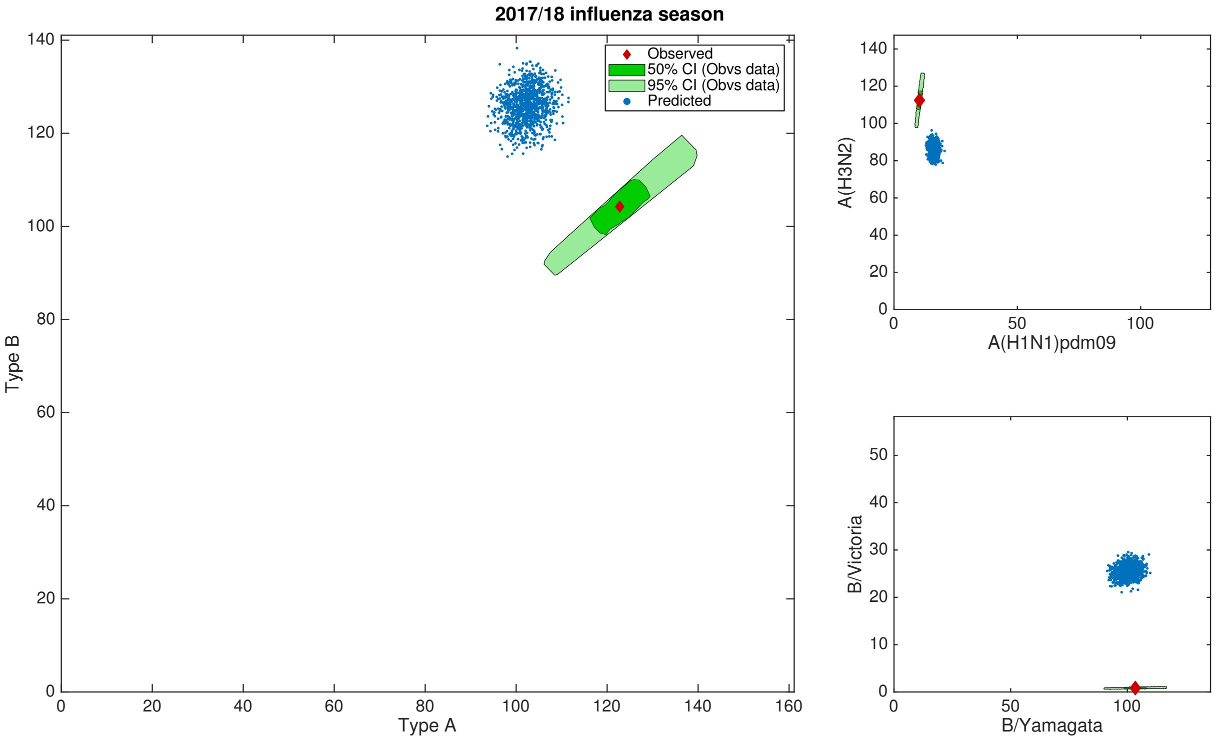

**Fig. S11: Posterior predictive influenza positive GP visit distributions: 2017/18 influenza season.** Empirical data point estimate and bootstrap samples are depicted by the red diamond and green shaded regions respectively. Blue dots denote posterior predicted replicates. **(left)** Type A versus type B; **(top right)** influenza A subtypes, A(H1N1)pdm09 versus A(H3N2); **(bottom right)** influenza B lineages, B/Yamagata versus B/Victoria.

#### 4.2 Fitting to subsets of the empirical data

To delve into model robustness, we sought parameter fits using subsets of the historical influenza season data. Categorically, we fit to two datasets. The first covered four influenza seasons spanning 2012/13-2015/16. The second also included the 2016/17 influenza season, bringing the total number of influenza seasons to five.

Invoking the adaptive population Monte Carlo ABC scheme upon each dataset, we obtained 10,000 parameter sets following the completion of 1,100 and 1,000 generations respectively (Figs. S12 and S13). To view detailed summary statistic per parameter, we refer the reader to Table S2.

We witness agreement with the statistical fit to the full dataset on the following aspects : (i) influenza A transmissibility exceeding type B virus transmissibility; (ii) amongst the three mechanisms for immunity propagation, retained natural infection in the previous influenza season having the greatest influence; (iii) little support for immunity propagation originating from influenza B cross-reactivity ($b$ attaining values near 1); (iv) quantitatively similar ascertainment probabilities.

Whilst outcomes fitting to a range of influenza seasons generally exhibited qualitative consistency, there were notable quantitative differences. Transmissibility magnitudes were raised, with the transmissibility of the A(H3N2) subtype exceeding that of the A(H1N1)pdm09 subtype (whereas in the complete fit we inferred similar transmissibility values for the type A viruses). The elevated transmissibility levels were counteracted by the enlarged effect of prior infection and vaccination reducing susceptibility. Explicitly, while the parameter distributions inferred fitting to the period 2012/13-2017/18 indicated little residual vaccine immunity and an estimated 20% drop in strain-specific susceptibility as a result of natural infection in the previous season, when fitting to the shorter time window susceptibility at the outset of an influenza season against a given strain type saw a near 55% reduction if naturally infected in the prior season by a virus of that class, and roughly 40% of prior season vaccine efficacy was retained (Table S2).

In a similar fashion to the dataset encompassing influenza seasons ranging from 2012/13 up to and including 2017/18, we found strong, qualitative correspondence between simulations of the paramterised model and the data (Figs. S14 and S15).

**(a)**

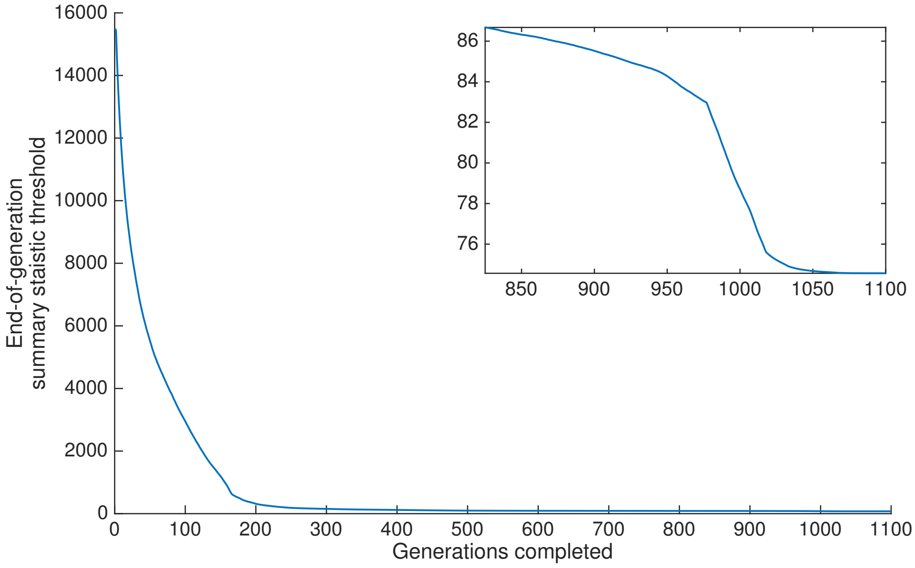

**(b)**

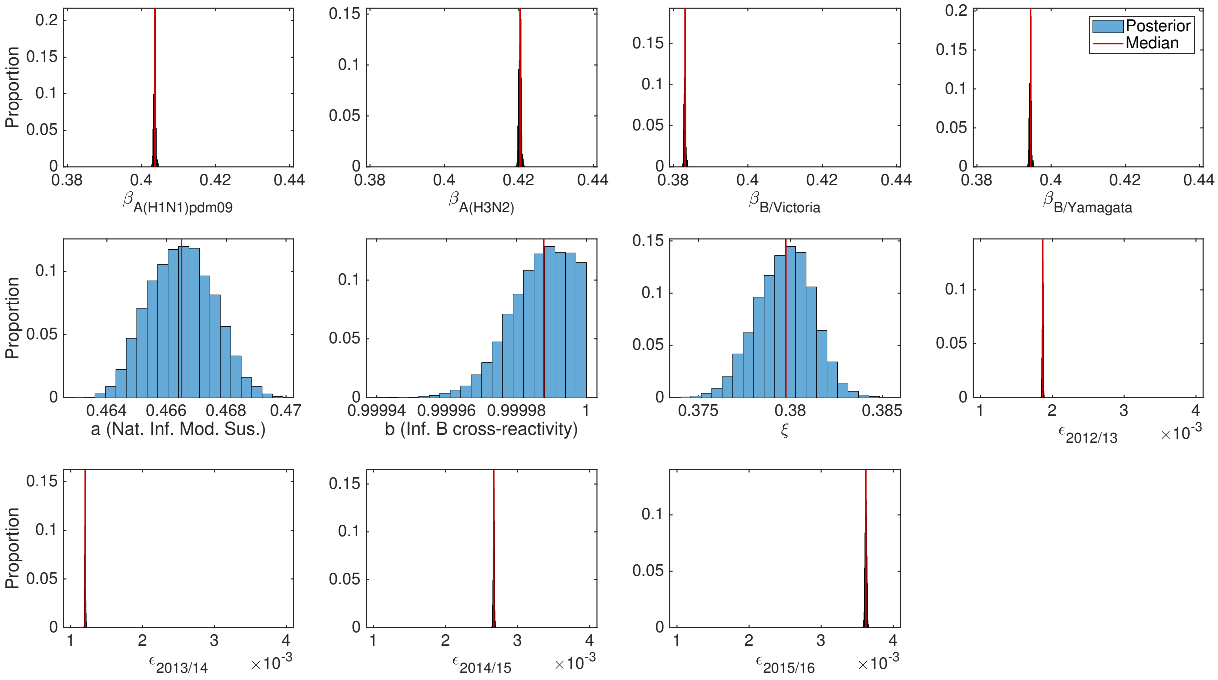

**Fig. S12: Results of the ABC scheme, fitting to empirical data covering 2012/13-2015/16 influenza seasons (inclusive). (a)** Summary metric threshold value upon completion of each generation of the inference scheme. The inset panel displays the latter quarter of generations. **(b)** Inferred parameter distributions estimated from 10,000 retained samples following completion of 1,100 generations of the inference scheme. Vertical red lines indicate the (non-weighted) median values for the model constants estimated from the inference procedure.

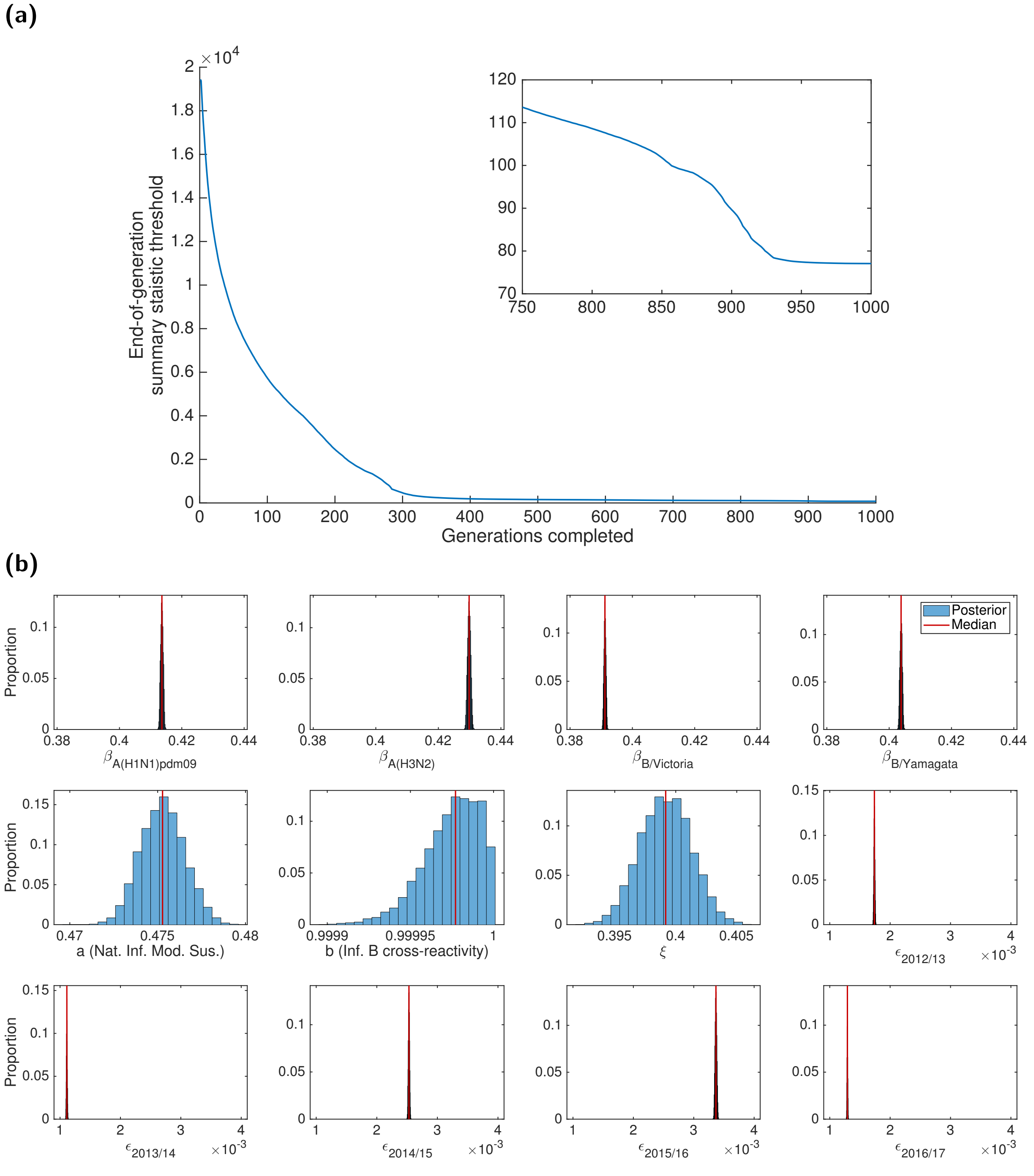

**Fig. S13: Results of the ABC scheme, fitting to empirical data covering 2012/13-2016/17 influenza seasons (inclusive). (a)** Summary metric threshold value upon completion of each generation of the inference scheme. The inset panel displays the latter quarter of generations. **(b)** Inferred parameter distributions estimated from 10,000 retained samples following completion of 1,000 generations of the inference scheme. Vertical red lines indicate the (non-weighted) median values for the model constants estimated from the inference procedure.

**(a)**

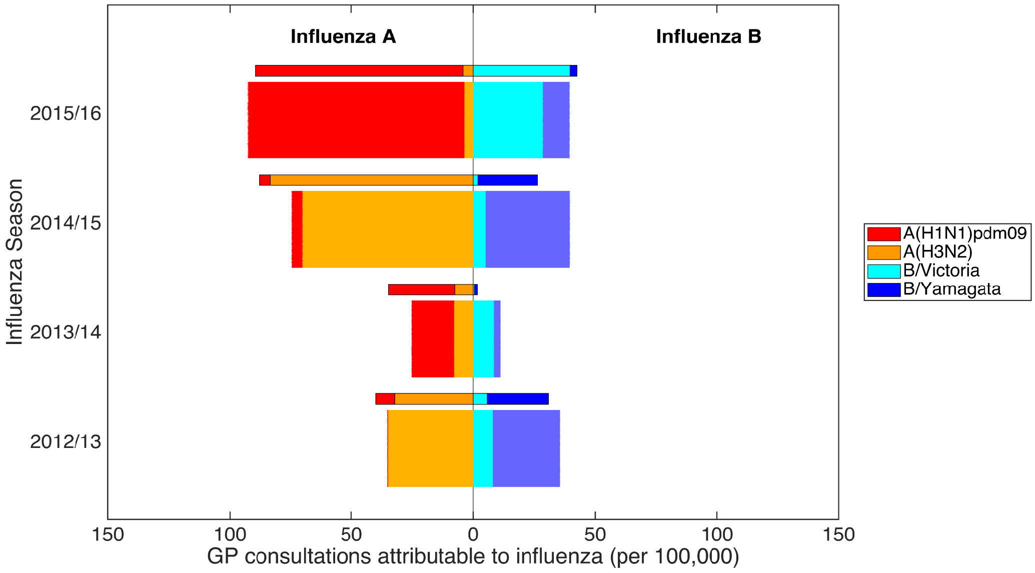

**(b)**

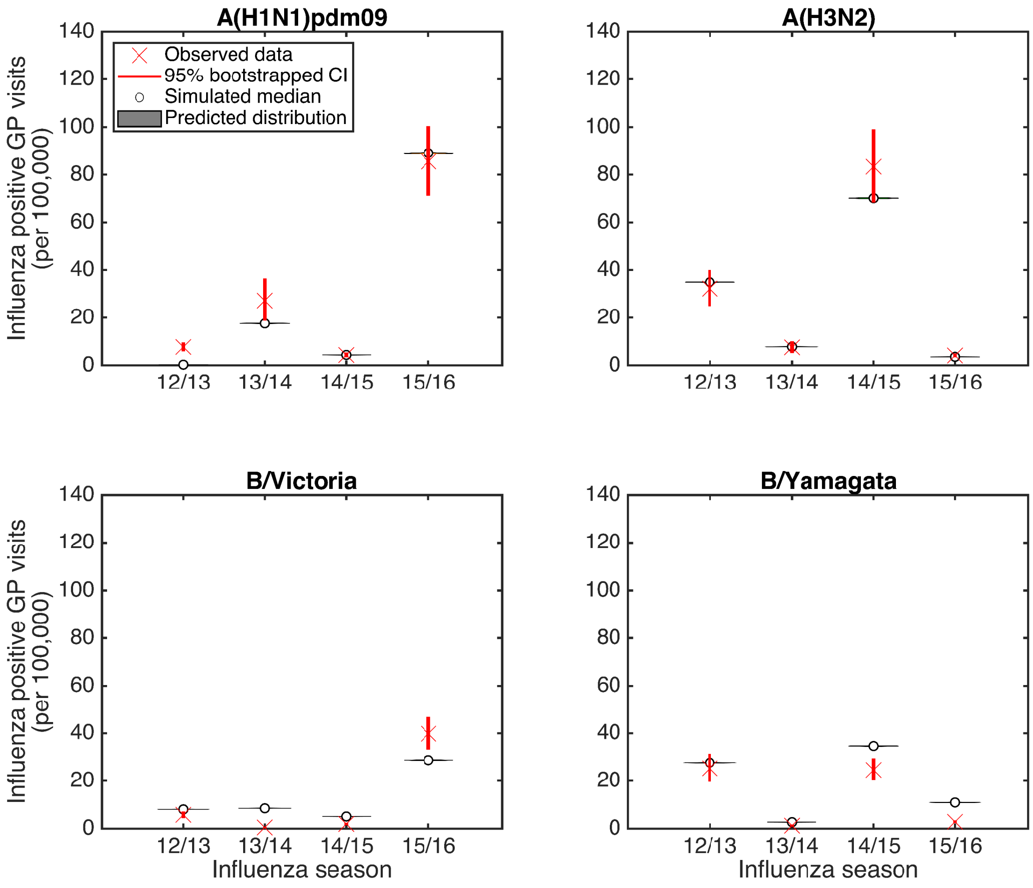

**Fig. S14: Posterior predictive influenza positive GP consultation distributions, post-fitting to the empirical data covering the 2012/13-2015/16 influenza seasons (inclusive).** We generated the estimated distributions from 1,000 model simulations, each using a distinct parameter set from the retained collection of particles. **(a)** Back-to-back stacked bars per simulation replicate. Each influenza season is topped out by a thicker stacked horizontal bar plot, corresponding to the strain-stratified point estimates for the empirical data. **(b)** Comparison of model simulated outcomes (shaded violin plots, with filled circles corresponding to the median value across the simulated replicates) versus the observed data (crosses denote the point estimate, with solid bars the range of the bootstrapped empirical data).

**(a)**

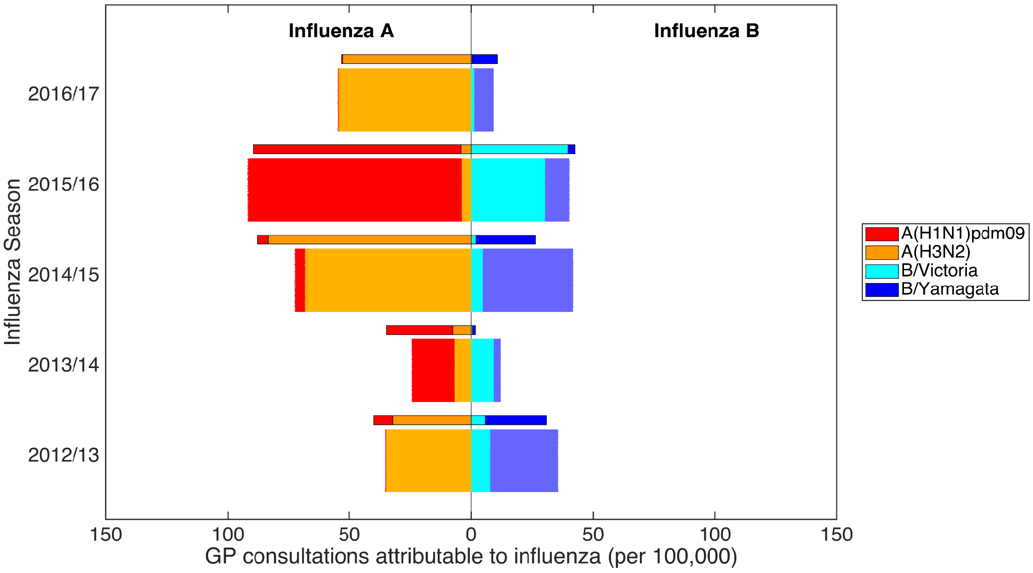

**(b)**

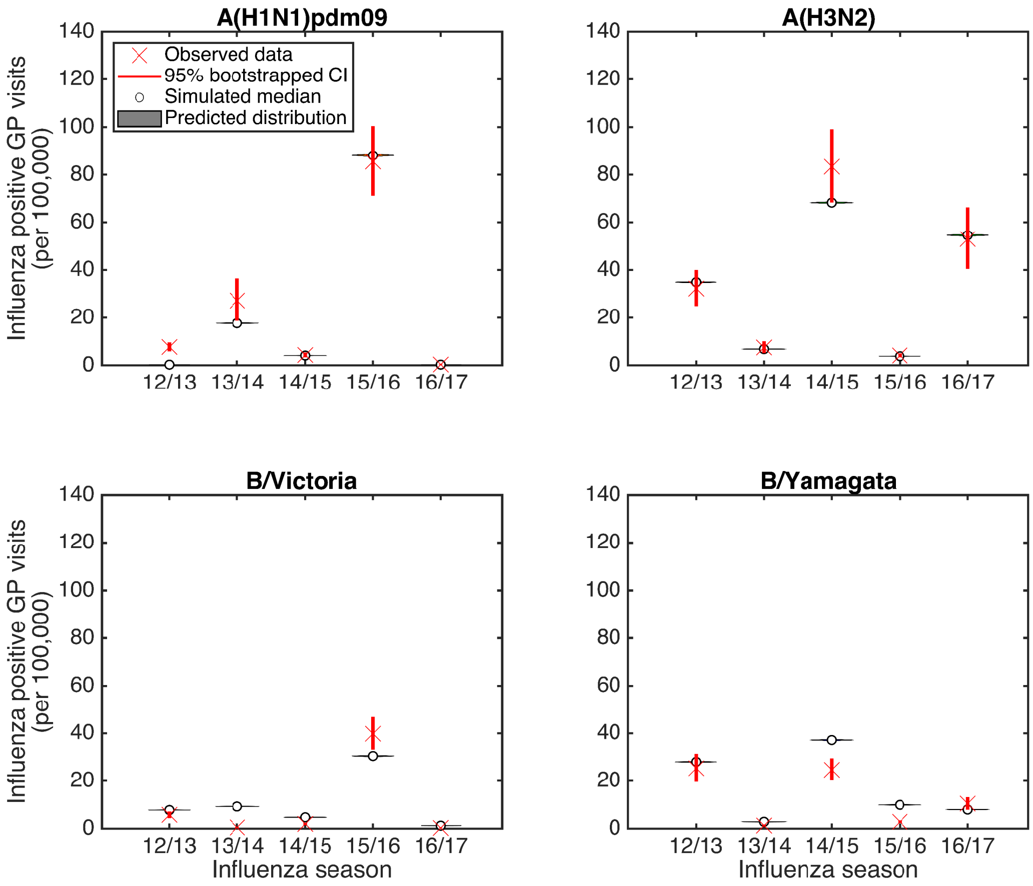

**Fig. S15: Posterior predictive influenza positive GP consultation distributions, post-fitting to the empirical data covering the 2012/13-2016/17 influenza seasons (inclusive).** We generated the estimated distributions from 1,000 model simulations, each using a distinct parameter set from the retained collection of particles. **(a)** Back-to-back stacked bars per simulation replicate. Each influenza season is topped out by a thicker stacked horizontal bar plot, corresponding to the strain-stratified point estimates for the empirical data. **(b)** Comparison of model simulated outcomes (shaded violin plots, with filled circles corresponding to the median value across the simulated replicates) versus the observed data (crosses denote the point estimate, with solid bars the range of the bootstrapped empirical data).

**Table S2: Values (posterior weighted quantiles) for the transmission, exposure history and ascertainment probability parameters inferred fitting to 2012/13-2015/16, 2012/13-2016/17 and 2012/13-2017/18 influenza season data.** Numbers inside brackets indicate 95% credible intervals. All values are given to 4 d.p.

| Description | Notation | Median [95% credible interval] | | |
|---|---|---|---|---|
| | | **2012/13-2015/16** | **2012/13-2016/17** | **2012/13-2017/18** |
| **Transmission parameters** | | | | |
| A(H1N1)pdm09 | $\beta_{\text{A(H1N1)pdm09}}$ | 0.4037 [0.4030, 0.4044] | 0.4136 [0.4128, 0.4144] | 0.3913 [0.3900, 0.4041] |
| A(H3N2) | $\beta_{\text{A(H3N2)}}$ | 0.4204 [0.4196, 0.4212] | 0.4298 [0.4288, 0.4310] | 0.3917 [0.3917, 0.4074] |
| B/Victoria | $\beta_{\text{B/Victoria}}$ | 0.3832 [0.3826, 0.3837] | 0.3912 [0.3905, 0.3918] | 0.3510 [0.3510, 0.3606] |
| B/Yamagata | $\beta_{\text{B/Yamagata}}$ | 0.3945 [0.3938, 0.3951] | 0.4038 [0.4030, 0.4046] | 0.3653 [0.3653, 0.3761] |
| **Exposure history parameters** | | | | |
| Natural infection in prior season | $a$ | 0.4667 [0.4642, 0.4689] | 0.4752 [0.4724, 0.4777] | 0.7883 [0.7847, 0.7964] |
| Type B influenza cross-reactivity | $b$ | 1.0000 [1.0000, 1.000] | 1.0000 [0.9999, 1.0000] | 0.9703 [0.9558, 0.9988] |
| Prior season vaccine efficacy carry over | $\xi$ | 0.3800 [0.3764, 0.3831] | 0.3996 [0.3944, 0.4038] | 0.0051 [0.0004, 0.0143] |
| **Ascertainment probabilities** | | | | |
| 2012/13 | $\epsilon_{2012/13}$ | 0.0019 [0.0018, 0.0019] | 0.0017 [0.0017, 0.0018] | 0.0017 [0.0015, 0.0018] |
| 2013/14 | $\epsilon_{2013/14}$ | 0.0012 [0.0012, 0.0012] | 0.0011 [0.0011, 0.0011] | 0.0009 [0.0009, 0.0012] |
| 2014/15 | $\epsilon_{2014/15}$ | 0.0027 [0.0026, 0.0027] | 0.0025 [0.0025, 0.0026] | 0.0024 [0.0022, 0.0026] |
| 2015/16 | $\epsilon_{2015/16}$ | 0.0036 [0.0036, 0.0036] | 0.0034 [0.0033, 0.0034] | 0.0037 [0.0032, 0.0039] |
| 2016/17 | $\epsilon_{2016/17}$ | — | 0.0013 [0.0013, 0.0013] | 0.0014 [0.0012, 0.0015] |
| 2017/18 | $\epsilon_{2017/18}$ | — | — | 0.0055 [0.0047, 0.0055] |

#### 4.3 Parameter identifiability

We generated and used a synthetic dataset to verify the parameter identifiability capabilities of our ABC inference scheme. To mimic the format of our empirical data, the synthetic data corresponded to strain-specific influenza positive ILI GP consultations rates per 100,000 population from the 2012/13 to 2017/18 influenza season (inclusive). The parameter values we used for producing the synthetic are listed in Table S3.

When fitting the model to the synthetic data, we employed our inference scheme in an equivalent manner as for fitting to the empirical data. We ably recovered the parameter values from which the synthetic data had been generated (Fig. S16 and Table S3). Furthermore, simulated outcomes from each parameter set had strong agreement with the synthetic data (Fig. S17).

**Table S3: Values (posterior weighted quantiles) for the transmission, exposure history and ascertainment probability parameters inferred fitting to synthetic data.** Numbers inside brackets indicate 95% credible intervals. All values are given to 4 d.p.

| Description | Notation | 'True' Value | Median [95% CI] |
|---|---|---|---|
| **Transmission parameters** | | | |
| A(H1N1)pdm09 | $\beta_{\text{A(H1N1)pdm09}}$ | 0.4200 | 0.4202 [0.4170, 0.4231] |
| A(H3N2) | $\beta_{\text{A(H3N2)}}$ | 0.4300 | 0.4302 [0.4268, 0.4333] |
| B/Victoria | $\beta_{B/\text{Victoria}}$ | 0.4150 | 0.4152 [0.4122, 0.4179] |
| B/Yamagata | $\beta_{B/\text{Yamagata}}$ | 0.4050 | 0.4052 [0.4024, 0.4077] |
| **Exposure history parameters** | | | |
| Modified susceptibility given natural infection in prior season | $a$ | 0.5837 | 0.5838 [0.5810, 0.5863] |
| Modified susceptibility due to type B influenza cross-reactivity | $b$ | 0.4351 | 0.4357 [0.4128, 0.4566] |
| Prior season vaccine efficacy carry over | $\xi$ | 0.7026 | 0.7037 [0.6930, 0.7124] |
| **Ascertainment probabilities** | | | |
| 2012/13 | $\epsilon_{2012/13}$ | 0.0015 | 0.0015 [0.0015, 0.0015] |
| 2013/14 | $\epsilon_{2013/14}$ | 0.0007 | 0.0007 [0.0007, 0.0007] |
| 2014/15 | $\epsilon_{2014/15}$ | 0.0023 | 0.0023 [0.0023, 0.0023] |
| 2015/16 | $\epsilon_{2015/16}$ | 0.0021 | 0.0021 [0.0020, 0.0022] |
| 2016/17 | $\epsilon_{2016/17}$ | 0.0004 | 0.0004 [0.0004, 0.0004] |
| 2017/18 | $\epsilon_{2017/18}$ | 0.0030 | 0.0030 [0.0030, 0.0031] |

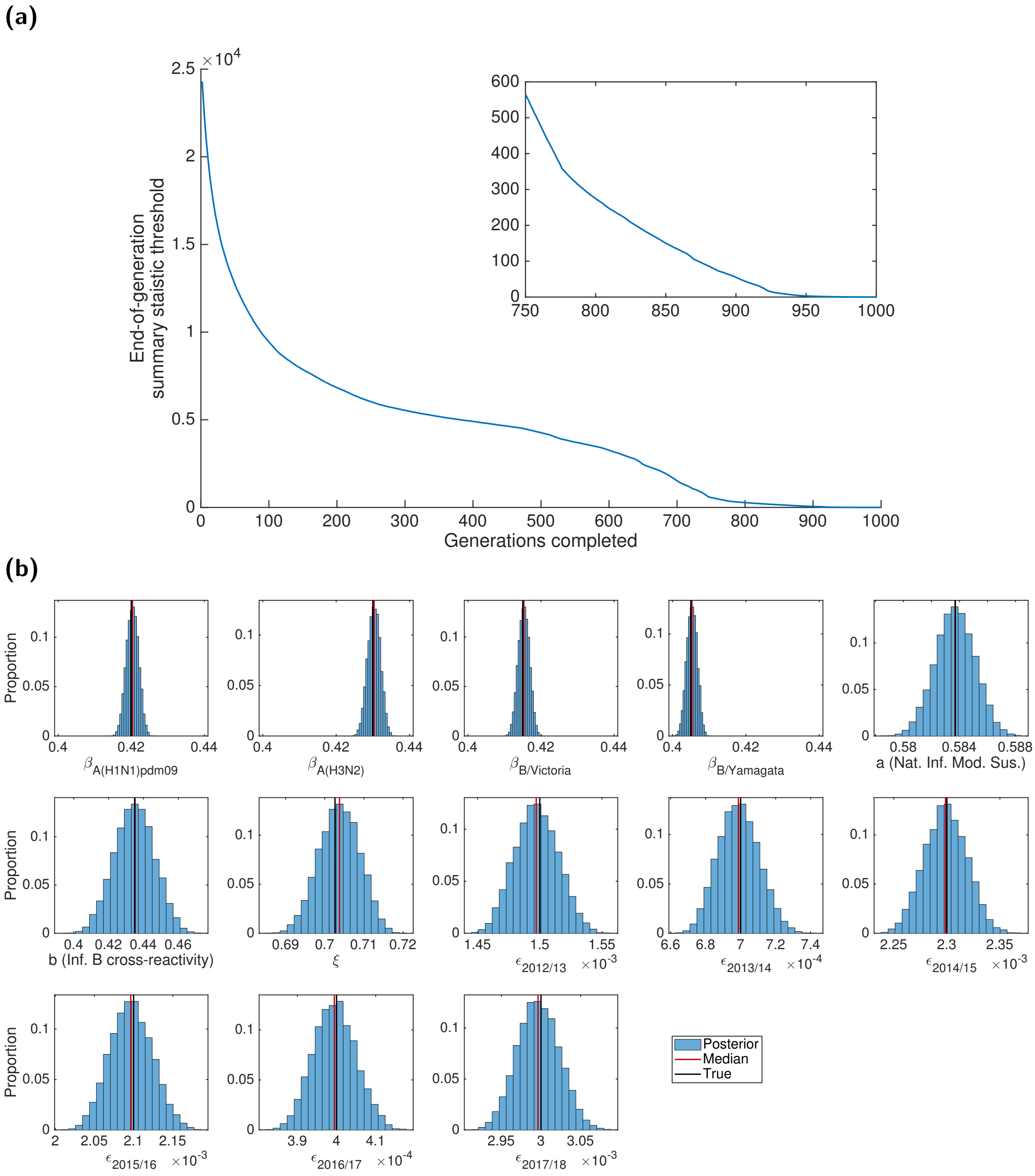

**Fig. S16: Results of the ABC scheme, fitting to synthetic data.** **(a)** Summary metric threshold value upon completion of each generation of the inference scheme. The inset panel displays the latter quarter of generations. **(b)** Inferred parameter distributions estimated from 10,000 retained samples following completion of 1,000 generations of the inference scheme. Vertical red lines indicate the (non-weighted) median values for the model constants estimated from the inference procedure, and vertical black lines correspond to the true values of the parameters from which the data were generated. We ably recovered the parameter values from which the synthetic data had been generated.

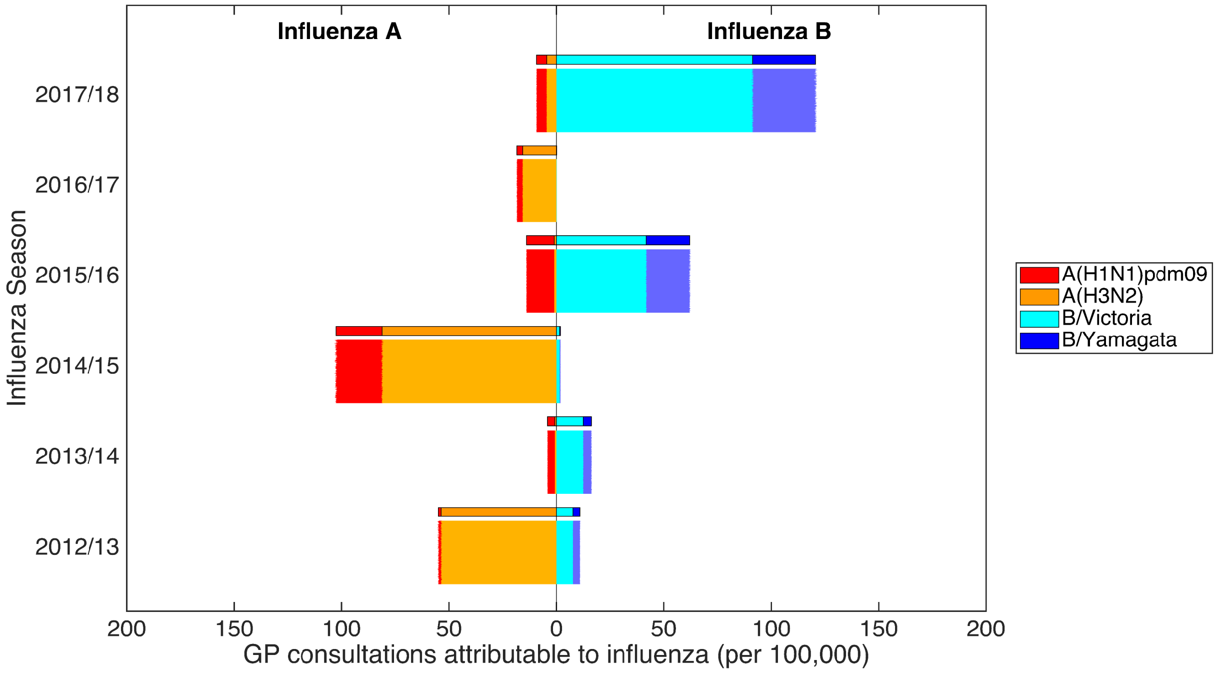

**Fig. S17: Posterior predictive distributions for influenza positive GP consultations per 100,000 population, using the parameter sets generated when fitting to the synthetic data.** Stratified by influenza season, we present back-to-back stacked bars for 1,000 simulation replicates, each using a distinct parameter set representing a sample from the posterior distribution when fitting our mathematical model to the synthetic data. Each influenza season is topped by a thicker stacked horizontal bar plot, corresponding to the synthetic data values. The left side depicts the cumulative total of ILI GP consultations attributable to type A influenza per 100,000 population (red shading denoting the A(H1N1)pdm09 subtype, orange shading the A(H3N2) subtype). In an equivalent manner, the right side stacked horizontal bars present similar data for type B influenza (cyan shading denoting the B/Victoria lineage, dark blue shading the B/Yamagata lineage). Overall, simulated outcomes from each parameter set had strong agreement with the synthetic data.

##### 4.4 Forward simulations

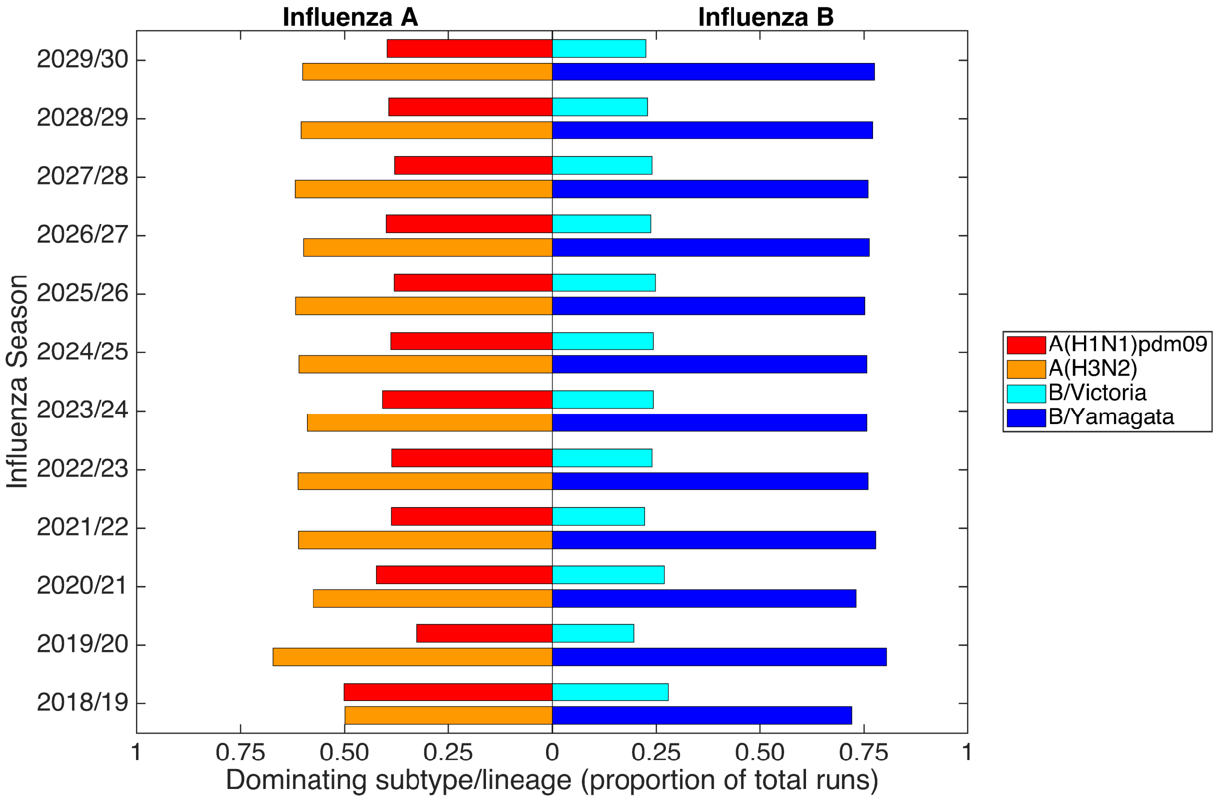

**Fig. S18: Quantification of the projected dominating influenza A subtype and influenza B lineage by influenza season, up to 2029/30.** The left half depicts the proportion of simulations in which the major subtype contributor (>50% of cases) to the overall type A influenza incidence was A(H1N1)pdm09 (red bars) or A(H3N2) (orange bars). In an equivalent manner, the right half depicts the proportion of simulations in which the major lineage contributor (>50% of cases) to the overall type B influenza incidence was B/Victoria (cyan bars) or B/Yamagata (dark blue bars). Constructed from simulation runs where vaccine efficacy against each strain were randomly sampled from the empirical distribution (totalling 1,000 replicates). For all forward simulated seasons, vaccine uptake matched that of the 2017/18 influenza season.

### 5 Model extension: Immunity propagation across multiple influenza seasons

In our extended model, we altered the immunity propagation model component. The vaccination, epidemiological and observation model components were all unchanged.

We display a modified directed acyclic graph, which includes the revised immunity propagation component structure and incorporation of the data streams (Fig. S19).

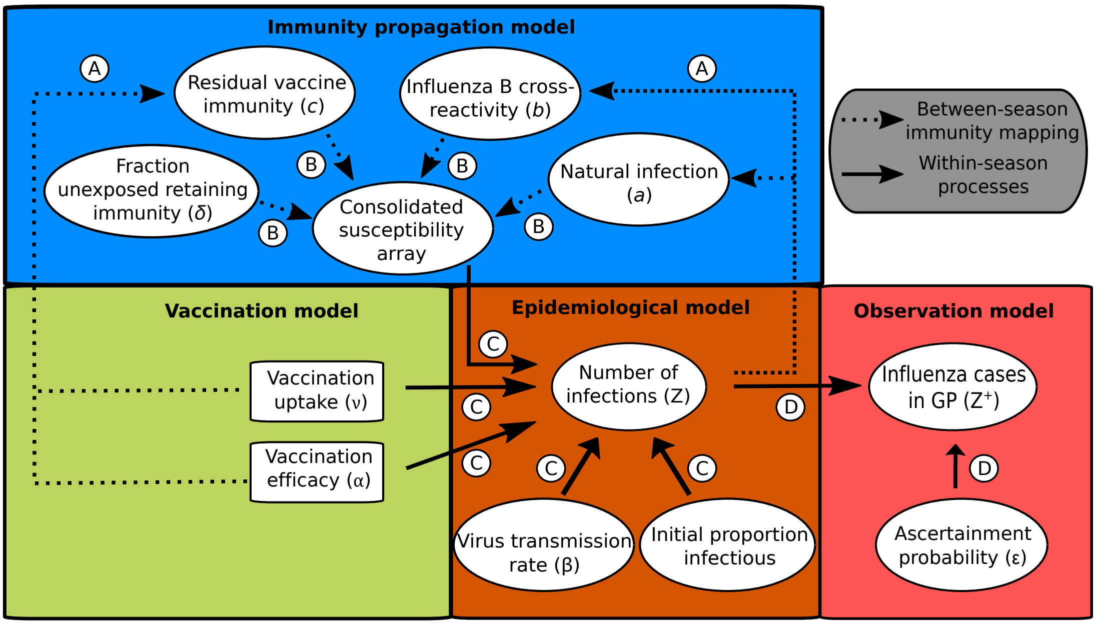

**Fig. S19: Schematic showing the links between the vaccination, immunity propagation, epidemiological and observation model components for the extended model.** We adopt the visualisation conventions of [26], with ellipses indicating variables, and rectangles indicating data. Dotted arrows indicate relationships between prior season epidemiological outcomes and immunity propagation factors. Solid arrows indicate within season processes. Circled capitalised letters indicate the relationships connecting the variables or data involved. These relationships are: process A, propagation of immunity as a result of exposure to influenza virus in the previous influenza season (through natural infection or vaccination); process B, modulation of current influenza season virus susceptibility; process C, estimation of influenza case load via the SEIR model of transmission; process D, ascertainment of cases through ILI recording at GP.

#### 5.1 Details of the expanded immunity propagation mechanism

As before, we tracked immunity derived from natural infection and vaccination separately. We kept our three, original susceptibility modifying factors: (i) modified susceptibility to strain $m$ given infection by a strain $m$ type virus the previous season, denoted $a$; (ii) carry over cross-reactivity protection between influenza B lineages, denoted $b$ (to account for infection with one influenza B virus lineage being potentially beneficial in protecting against subsequent infection with either influenza B virus lineage [27]); (iii) residual strain-specific protection carried over from the prior season influenza vaccine, denoted $c_m$. We again mandated that $0 < a,b,c_m < 1$.

To enhance the flexibility of the immunity propagation component, we introduced and fit an additional parameter, $\delta$, representing the proportion of those who both began the influenza season in an exposure history group linked to natural infection (Fig. 3: rows 2-5, 7-10) and were unexposed to influenza virus during the current season (who at the end of of the current influenza season remained susceptible), that retained their pre-existing natural infection acquired immunity. Those keeping immunity arising from natural infection were mapped to the relevant prior infection exposure history group dependent upon current season vaccination status. The remaining proportion $(1-\delta)$ transitioned in the same manner as in the original model, reverting to either the naive exposure history group or vaccinated only exposure history group (Fig. S20).

For clarity, there was no mechanism in the model framework to confer vaccine-induced immunity beyond one influenza season (i.e. vaccine-induced immunity could be retained for, at most, a single additional influenza season). As before, the collection of exposure history groupings and associated strain-specific susceptibilities were consolidated into a single susceptibility array (Fig. S19, process B; Fig. 3).

##### 5.2 Between season exposure history group assignments

At the beginning of each influenza season (1st September), we enacted the following exposure history group assignments:

- $\left\{S^{N,h=\bar{N}}, S^{N,h=\bar{V}}, (1-\delta)S^{N,h=m}, (1-\delta)S^{N,h=m\&\bar{V}}\right\} \rightarrow S^{N,h=\bar{N}}$
- $\left\{S^{V,h=\bar{N}}, S^{V,h=\bar{V}}, (1-\delta)S^{V,h=m}, (1-\delta)S^{V,h=m\&\bar{V}}\right\} \rightarrow S^{N,h=\bar{V}}$
- $\left\{\delta S^{N,h=m}, \delta S^{N,h=m\&\bar{V}}, E^{N,m}, I^{N,m}, R^{N,m}\right\} \rightarrow S^{N,h=m}$
- $\left\{\delta S^{V,h=m}, \delta S^{V,h=m\&\bar{V}}, E^{V,m}, I^{V,m}, R^{V,m}\right\} \rightarrow S^{N,h=m\&\bar{V}}$

where $X$ represents vaccination status, with $N$ corresponding to non-vaccinated and $V$ vaccinated in the current influenza season (bars on vaccination state symbols relate to vaccination status in the previous influenza season).

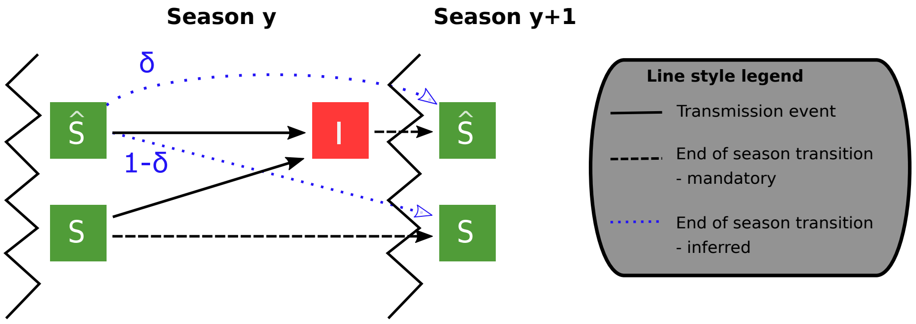

**Fig. S20: Mechanism for conferring immunity to influenza across multiple influenza seasons.** At the beginning of a generic influenza season $y$, those susceptible may be divided into two categories based on exposure history: (i) susceptibles in either the naive exposure history group , with unmodified susceptibility to infection, or in the vaccinated only exposure history group (labelled $S$); (ii) all those that are susceptible and in an exposure history group involving natural infection (labelled $\hat{S}$). During the influenza season, those that were initially susceptible may be infected (state $I$). Of those remaining in state $\hat{S}$ at the end of season $y$, a proportion $\delta$ are retained in state $\hat{S}$ at the beginning of season $y+1$. The remainder (1 - $\delta$) transition to state $S$. Note, $\delta = 0$ corresponds to no residual immunity being retained due to natural infection beyond one subsequent influenza season, while $\delta = 1$ coincides with residual immunity being kept in all future seasons until the occurrence of a new infection event.

**Table S4:** Overview of parameters in the extended model.

| Description | Notation | Value | Sources |
|---|---|---|---|
| **Fixed parameters** | | | |
| Mortality rate (day$^{-1}$) | B,D | $\frac{1}{81\times365}$ | [23] |
| Rate of latency loss, influenza A subtypes (day$^{-1}$) | $\gamma_{1,A}$ | $\frac{1}{1.4}$ | [28] |
| Rate of latency loss, influenza B lineages (day$^{-1}$) | $\gamma_{1,B}$ | $\frac{1}{0.6}$ | [28] |
| Recovery rate (day$^{-1}$) | $\gamma_2$ | $\frac{1}{3.8}$ | [29] |
| **Time-varying parameters** | | | |
| Vaccination rate at time t | $\nu(t)$ | — | [6, 7, 11] |
| Vaccine efficacy, season $y$ strain $m$ | $\alpha_m^y$ | — | [14–22] |

| Inferred parameter description | Notation | Prior |
|---|---|---|
| Influenza virus transmissibility, strain $m$ | $\beta_m$ | $\mathcal{U}(0.2632,0.7896)$ |
| Modified susceptibility given natural infection in prior season | $a$ | $\mathcal{U}(0,1)$ |
| Modified susceptibility due to type B influenza cross-reactivity | $b$ | $\mathcal{U}(0,1)$ |
| Proportion of prior season vaccine efficacy carried over | $\xi$ | $\mathcal{U}(0,1)$ |
| Fraction unexposed retaining immunity | $\delta$ | $\mathcal{U}(0,1)$ |
| Ascertainment probability in season $y$ | $\epsilon_y$ | $\mathcal{U}(0,0.05)$ |

#### 5.3 Fitting the extended model to the data

We performed parameter inference using an equivalent ABC scheme to that used to fit the original model. For the extended model, we fit the following collection of parameters (Table S4): transmissibility of each influenza virus strain ($\beta_m$), modified susceptibility to strain $m$ given infection by a strain $m$ type virus the previous season ($a$), carry over cross-reactivity protection between influenza B lineages ($b$), residual protection carried over from the prior season influenza vaccine ($\xi$), the fraction unexposed in the prior influenza season that retain immunity ($\delta$), and an ascertainment probability per influenza season ($\epsilon_y$).

We amassed 10,000 parameter sets (representing a sample from the posterior distribution) to determine credible values of the model parameters (Table S5). Samples were obtained after completion of 750 generations of the adaptive-population Monte Carlo ABC scheme, at which point the generation-by-generation tolerance level updates were minor (Fig. S21(a)). Note that the error values attained for the parameter sets inferred for the extended model closely matched the error values for the parameter sets inferred for the original (less complex) model. The inferred model parameters for the extended model were well defined with Gaussian-shaped histograms (Fig. S21(b)).

Comparing the parameter fits to those for the original model, we maintained the main outcome of propagation of immunity being weaker if vaccine derived, compared to natural immunity from infection. As before, we inferred little carry over of prior season vaccine efficacy, with the majority of the posterior distribution for $\xi$ massed near 0.

We found several other similarities amongst the transmission parameters and ascertainment probabilities when comparing the two model fits. Virus transmission rates were larger for the two influenza A subtypes than than the corresponding estimates for the two type B lineages, and B/Yamagata transmissibility estimates exceeded those for B/Victoria. Furthermore, for each influenza season the ascertainment probability distributions closely resembled each other. An exception was, for all strains, inferred virus transmission rate distributions for the extended model being shifted towards larger values.

Discrepancies did arise amongst parameters tied to immunity processes. First, a decrease in value for parameter $a$, corresponding to the modified susceptibility to a given influenza type that results from natural infection by a prior influenza virus of that type (median values, extended model vs original model: 0.7168 vs 0.7883). These predictions correspond to an almost 30% reduction in susceptibility in the extended model compared to little more than 20% in the original model. Second, posterior distributions for $b$ imply a similar amount of modulation to susceptibility via influenza B cross-reactivity. This is in stark contrast to the less complex model, where there was little support for carry over of type B influenza cross-reactivity from one influenza season to the next (median values: 0.7214 vs 0.9703).

Inspecting the posterior distribution for $\delta$, the majority of samples attained values greater than 0.9. The inferred magnitude implies long term retention of immunity acquired from natural infection (assuming no further vaccination or infection event to alter the immunity landscape), in agreement with prior studies supporting infection acquired immunity waning over a time scale exceeding a year [30–32]).

Performing 1,000 independent simulations of the extended model with parameter sets drawn from the ABC inference procedure, we found a remarkably similar fit to the data with the extended model (Fig. S22) compared to the less complex, original model (Figs. 5-6). Thus, despite the added complexity, the extended model did not give noticeable improvements in correspondence of model outputs with the data (relative to fits with a model using the simpler immunity propagation setup).

**Table S5: Values (posterior weighted quantiles) for the transmission, exposure history and ascertainment probability parameters inferred fitting the extended model to the empirical data.** Numbers inside brackets indicate 95% credible intervals. All values are given to 4 d.p.

| **Description** | **Notation** | **Median [95% credible interval]** |
|---|---|---|
| **Transmission parameters** | | |
| A(H1N1)pdm09 | $\beta_{\text{A(H1N1)pdm09}}$ | 0.4112 [0.4094, 0.4139] |
| A(H3N2) | $\beta_{\text{A(H3N2)}}$ | 0.4302 [0.4281, 0.4336] |
| B/Victoria | $\beta_{\text{B/Victoria}}$ | 0.3620 [0.3600, 0.3634] |
| B/Yamagata | $\beta_{\text{B/Yamagata}}$ | 0.3735 [0.3722, 0.3748] |
| **Exposure history parameters** | | |
| Natural infection in prior season | $a$ | 0.7168 [0.7146, 0.7181] |
| Type B influenza cross-reactivity | $b$ | 0.7214 [0.6978, 0.7530] |
| Prior season vaccine efficacy propagation | $\xi$ | 0.0004 [0.0000, 0.0012] |
| Fraction unexposed retaining immunity | $\delta$ | 0.9553 [0.9310, 0.9907] |
| **Ascertainment probabilities** | | |
| 2012/13 | $\epsilon_{2012/13}$ | 0.0018 [0.0018, 0.0019] |
| 2013/14 | $\epsilon_{2013/14}$ | 0.0011 [0.0010, 0.0011] |
| 2014/15 | $\epsilon_{2014/15}$ | 0.0025 [0.0024, 0.0026] |
| 2015/16 | $\epsilon_{2015/16}$ | 0.0038 [0.0038, 0.0039] |
| 2016/17 | $\epsilon_{2016/17}$ | 0.0014 [0.0014, 0.0015] |
| 2017/18 | $\epsilon_{2017/18}$ | 0.0052 [0.0052, 0.0053] |

**(a)**

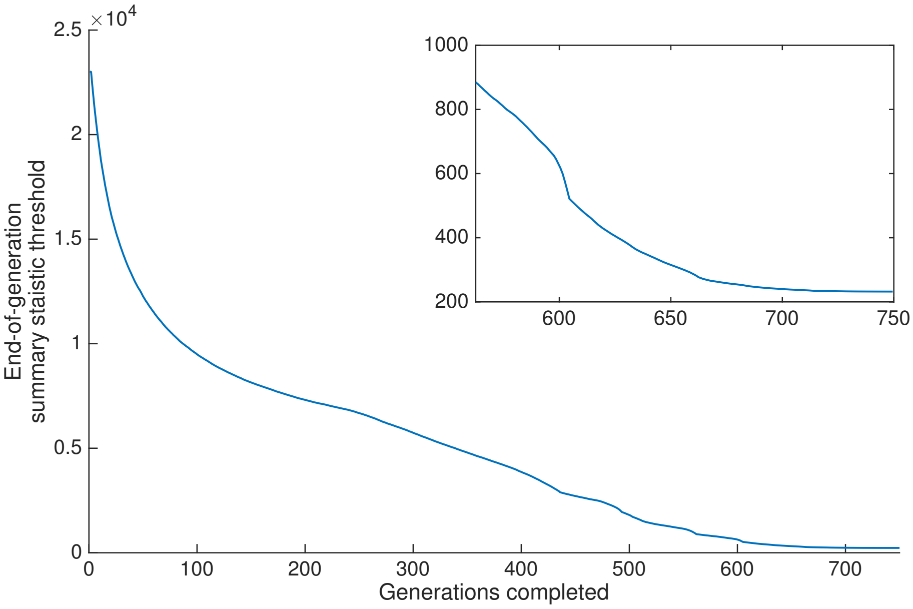

**(b)**

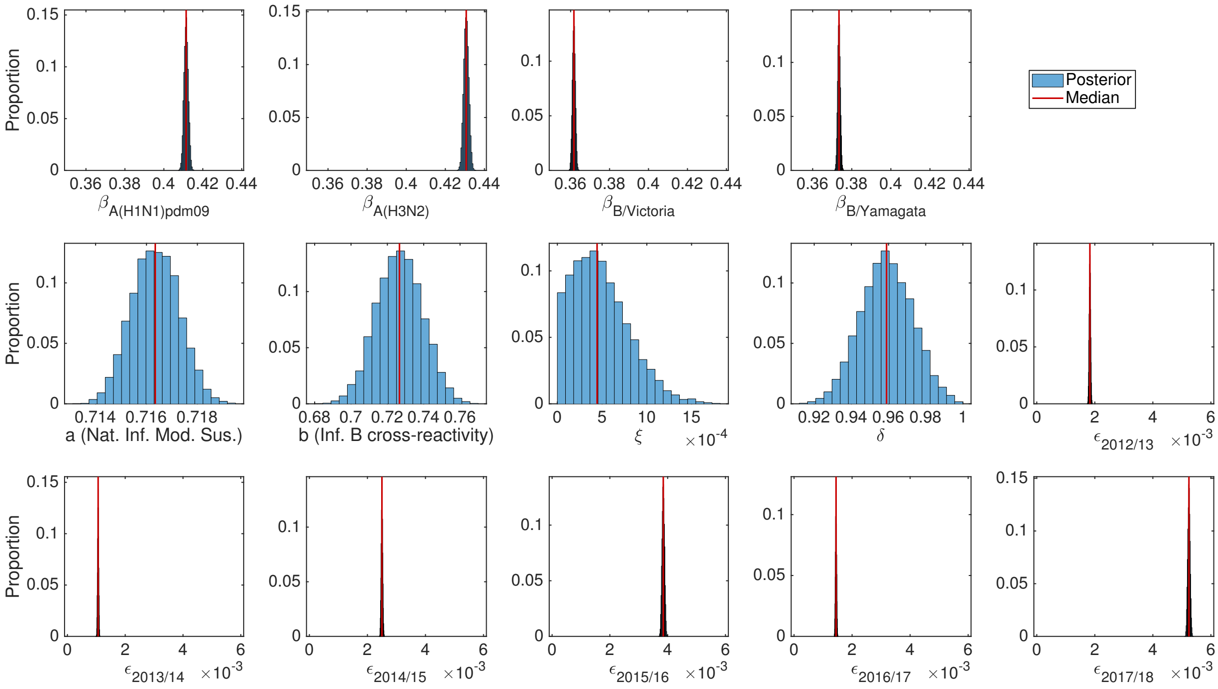

**Fig. S21: Results of the ABC scheme, fitting the extended model to the empirical data.** **(a)** Summary metric threshold value upon completion of each generation of the inference scheme. The inset panel displays the latter quarter of generations. **(b)** Inferred parameter distributions estimated from 10,000 retained samples following completion of 750 generations of the inference scheme. Vertical red lines indicate the (non-weighted) median values for the model constants estimated from the inference procedure. Particularly noteworthy outcomes include: transmissibility of type A viruses exceeding type B viruses; vaccine carry over had little impact on present season susceptibility; support for immunity stemming from natural infection being retained long term; the highest ascertainment probability occurred in the 2017/18 influenza season.

**(a)**

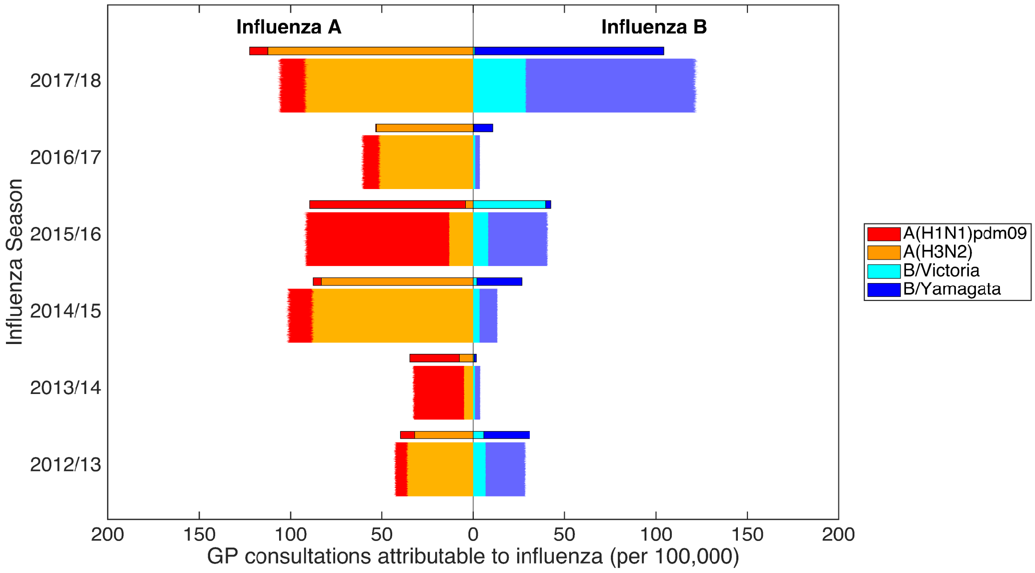

**(b)**

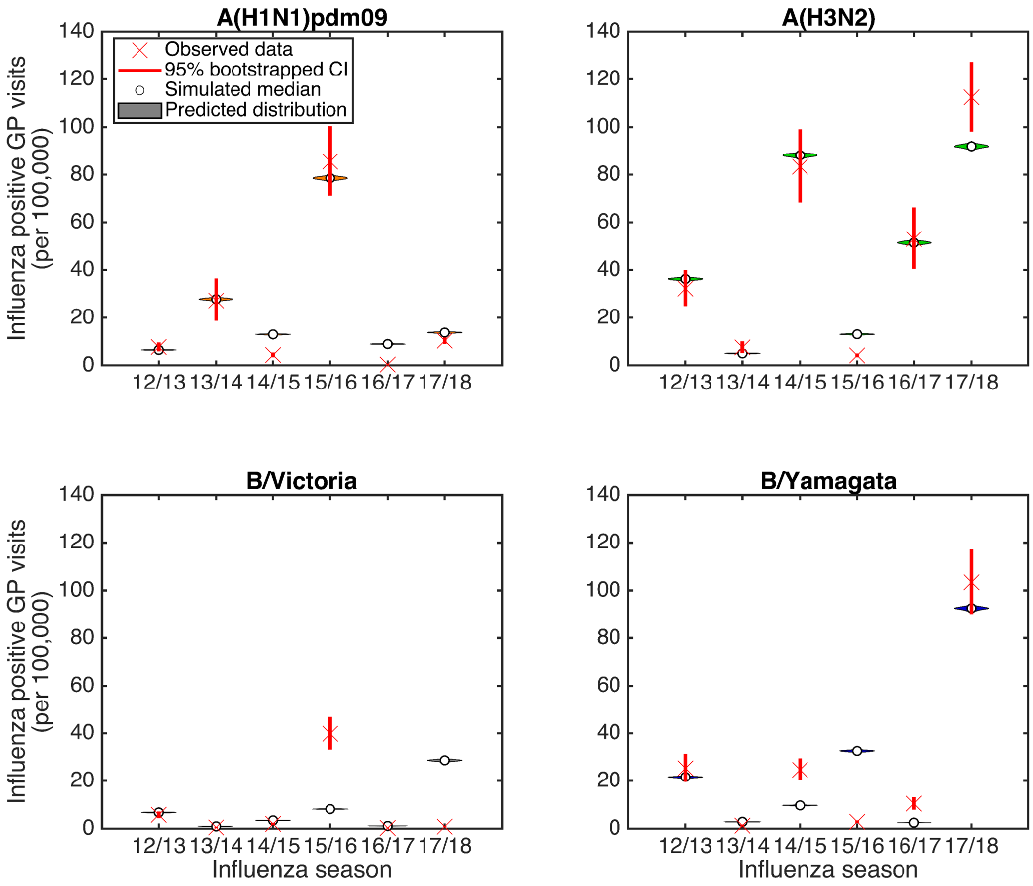

**Fig. S22: Posterior predictive influenza positive GP consultation distributions, post-fitting the extended model to the empirical data covering the 2012/13-2017/18 influenza seasons (inclusive).** We generated the estimated distributions from 1,000 model simulations, each using a distinct parameter set from the retained collection of particles. **(a)** Back-to-back stacked bars per simulation replicate. Each influenza season is topped out by a thicker stacked horizontal bar plot, corresponding to the strain-stratified point estimates for the empirical data. **(b)** Comparison of model simulated outcomes (shaded violin plots, with filled circles corresponding to the median value across the simulated replicates) versus the observed data (crosses denote the point estimate, with solid bars the range of the bootstrapped empirical data).

#### 5.4 Parameter identifiability

We again generated and used a synthetic dataset to verify the parameter identifiability capabilities of our ABC inference scheme under the revised model framework. The synthetic data corresponded to strain-specific influenza positive ILI GP consultations rates per 100,000 population from the 2012/13 to 2017/18 influenza season (inclusive), to mimic the format of our empirical data. The parameter values we used for producing the synthetic are listed in Table S6.

Once more, we ably recovered the parameter values from which the synthetic data had been generated (Fig. S23 and Table S6). Furthermore, simulated outcomes from each parameter set had strong agreement with the synthetic data (Fig. S24).

**Table S6: Values (posterior weighted quantiles) for the transmission, exposure history and ascertainment probability parameters inferred fitting the extended model to synthetic data.** Numbers inside brackets indicate 95% credible intervals. All values are given to 4 d.p.

| Description | Notation | 'True' Value | Median [95% CI] |
|---|---|---|---|
| **Transmission parameters** | | | |
| A(H1N1)pdm09 | $\beta_{\mathrm{A(H1N1)pdm09}}$ | 0.4200 | 0.4206 [0.4166, 0.4236] |
| A(H3N2) | $\beta_{\mathrm{A(H3N2)}}$ | 0.4300 | 0.4307 [0.4265, 0.4337] |
| B/Victoria | $\beta_{B/\mathrm{Victoria}}$ | 0.4150 | 0.4157 [0.4117, 0.4190] |
| B/Yamagata | $\beta_{B/\mathrm{Yamagata}}$ | 0.4050 | 0.4056 [0.4019, 0.4086] |
| **Exposure history parameters** | | | |
| Modified susceptibility given natural infection in prior season | $a$ | 0.5837 | 0.5835 [0.5776, 0.5902] |
| Modified susceptibility due to type B influenza cross-reactivity | $b$ | 0.4351 | 0.4365 [0.4102, 0.4581] |
| Prior season vaccine efficacy carry over | $\xi$ | 0.7026 | 0.7046 [0.6889, 0.7194] |
| Fraction unexposed retaining immunity | $\delta$ | 0.1000 | 0.1007 [0.0832, 0.1139] |
| **Ascertainment probabilities** | | | |
| 2012/13 | $\epsilon_{2012/13}$ | 0.0015 | 0.0015 [0.0015, 0.0015] |
| 2013/14 | $\epsilon_{2013/14}$ | 0.0007 | 0.0007 [0.0007, 0.0007] |
| 2014/15 | $\epsilon_{2014/15}$ | 0.0023 | 0.0023 [0.0023, 0.0023] |
| 2015/16 | $\epsilon_{2015/16}$ | 0.0021 | 0.0021 [0.0020, 0.0022] |
| 2016/17 | $\epsilon_{2016/17}$ | 0.0004 | 0.0004 [0.0004, 0.0004] |
| 2017/18 | $\epsilon_{2017/18}$ | 0.0030 | 0.0030 [0.0029, 0.0031] |

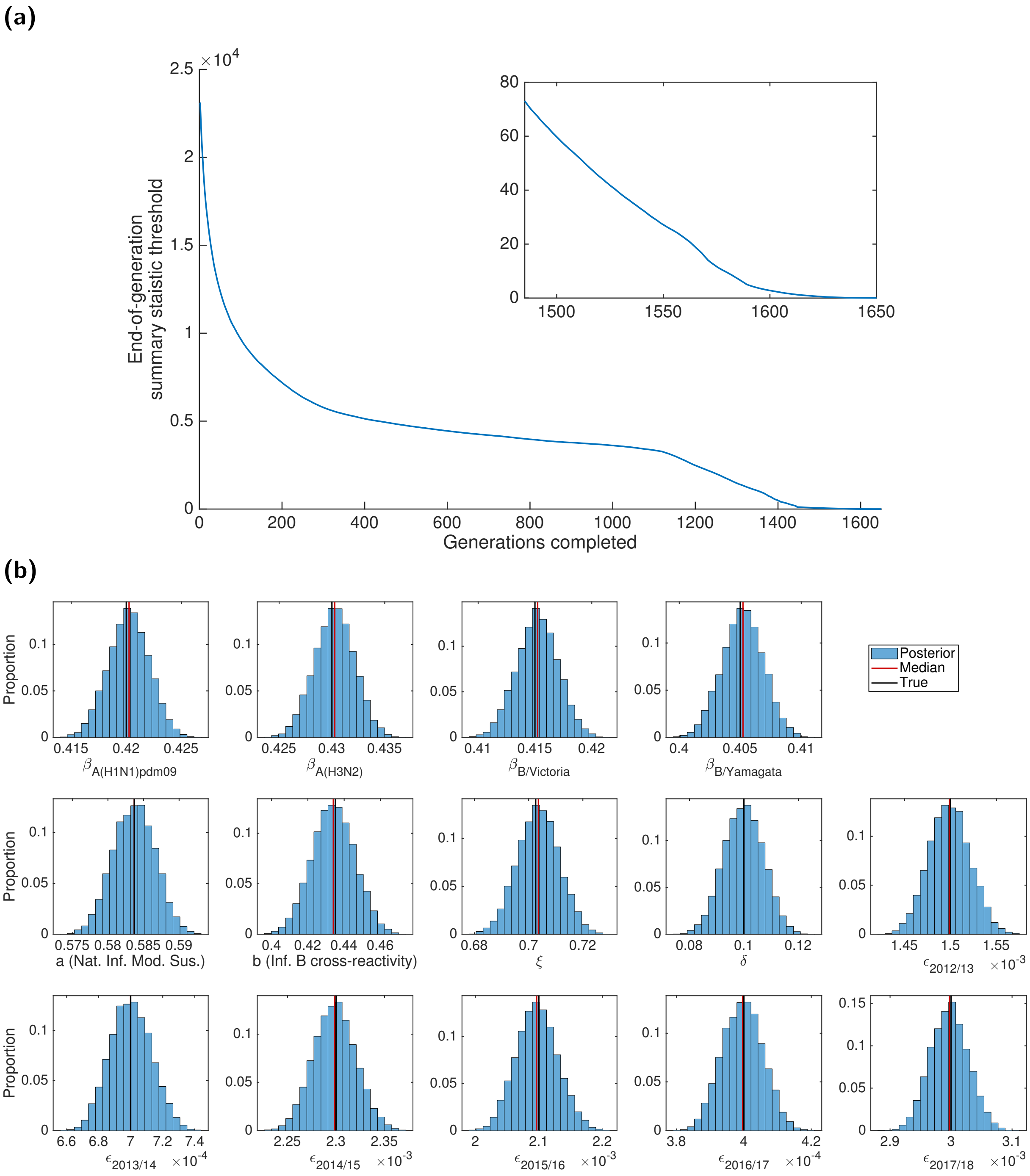

**Fig. S23: Results of the ABC scheme, fitting the extended model to synthetic data. (a)** Summary metric threshold value upon completion of each generation of the inference scheme. The inset panel displays the latter tenth of generations. **(b)** Inferred parameter distributions estimated from 10,000 retained samples following completion of 1,650 generations of the inference scheme. Vertical red lines indicate the (non-weighted) median values for the model constants estimated from the inference procedure, and vertical black lines correspond to the true values of the parameters from which the data were generated. We ably recovered the parameter values from which the synthetic data had been generated.

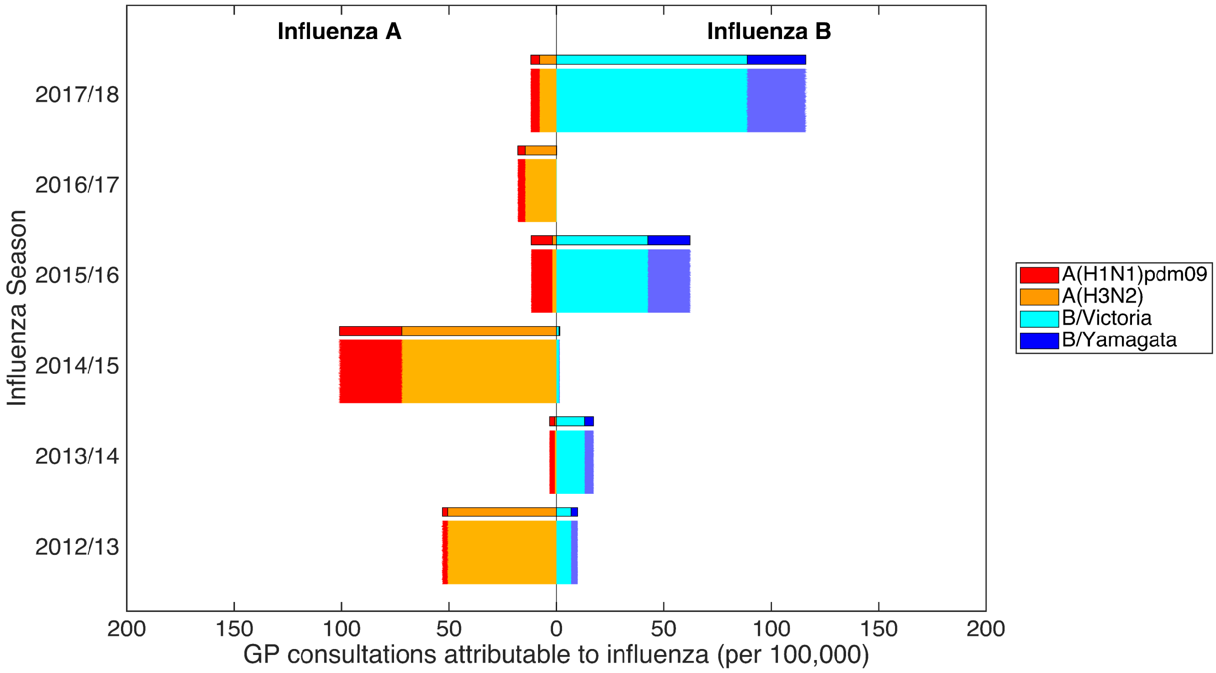

**Fig. S24: Posterior predictive distributions for influenza positive GP consultations per 100,000 population, using the parameter sets generated fitting the extended model to the synthetic data.** Stratified by influenza season, we present back-to-back stacked bars for 1,000 simulation replicates, each using a distinct parameter set representing a sample from the posterior distribution when fitting our mathematical model to the synthetic data. Each influenza season is topped by a thicker stacked horizontal bar plot, corresponding to the synthetic data values. The left side depicts the cumulative total of ILI GP consultations attributable to type A influenza per 100,000 population (red shading denoting the A(H1N1)pdm09 subtype, orange shading the A(H3N2) subtype). In an equivalent manner, the right side stacked horizontal bars present similar data for type B influenza (cyan shading denoting the B/Victoria lineage, dark blue shading the B/Yamagata lineage). Overall, simulated outcomes from each parameter set had strong agreement with the synthetic data.
