## Supplementary material for "Seasonal influenza: Modelling approaches to capture immunity propagation": Manuscript tracked changes (V1 to V2)

$$\begin{aligned} \text{GP consultation rate for strain } m \text{ in season } y \;=\; C_{m,y} \;=\; & \text{GP ILI consultation rate} \times \ldots \\ & \text{Proportion of ILI samples influenza positive} \times \ldots \\ & \text{Proportion of influenza viruses in circulation of strain type}\, m \end{aligned} \tag{1}$$

These computations resulted in seasonal rates of GP consultations attributable for the two influenza A subtypes and two influenza B lineages per 100,000 population (Fig. 1(a)). Uncertainty distributions for these values were obtained via bootstrapping given the sample sizes for the three components.

##### Vaccine uptake and efficacy

For the 2009/10 influenza season, we acquired vaccine uptake profiles from survey results on H1N1 vaccine uptake amongst patient groups in primary care [27]. For the 2010/11 influenza season onward we collated vaccine uptake information from PHE official statistics [24, 25].

We took age adjusted vaccine efficacy estimates for each historical influenza season (2009/2010-2017/2018 inclusive) from publications detailing end-of-season seasonal influenza vaccine effectiveness in the United Kingdom [28–34]. In seasons where equivalent publications were not available, we used mid-season or provisional end-of-season age adjusted vaccine efficacy estimates from PHE reports [35, 36].

There has been substantial variation in vaccine efficacy between seasons and also within seasons across strains, with estimates spanning 23-92% (Table 1). Further, there have been two instances of a mismatch between the vaccine and circulating viruses causing the vaccine to be ineffective (i.e. 0% efficacy) against a particular strain group (B/Yamagata in the 2010/11 influenza season, A(H3N2) in the 2017/18 influenza season). As a sole exception, for the 2009/10 influenza season we used the pandemic vaccine (rather than the seasonal influenza vaccine efficacy) to inform efficacy against A(H1N1)pdm09, with the effectiveness against all other strain types set to zero [28].

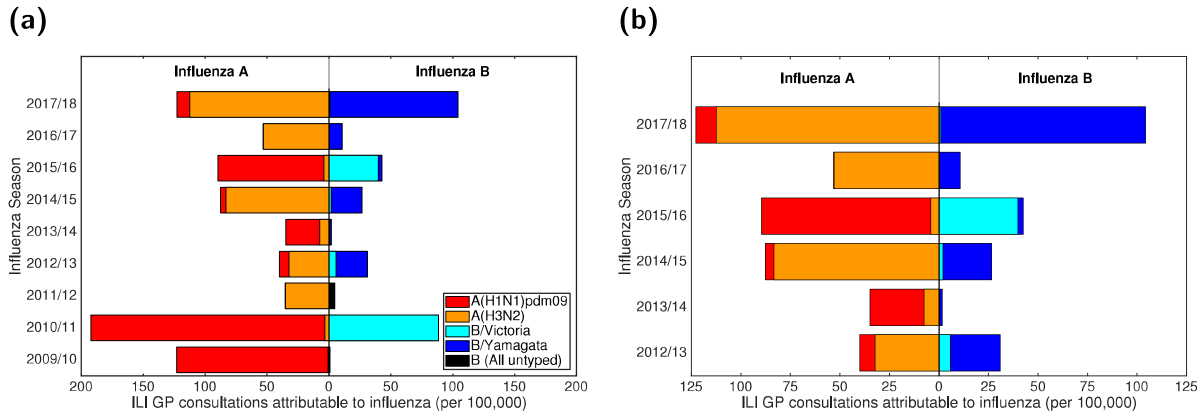


**Fig. 1: Empirical, strain-stratified data for ILI GP consultations attributable to influenza per 100,000 population.** In both panels, stacked horizontal bars in the left-half depict the cumulative total of ILI GP consultations attributable to type A influenza per 100,000 population (red shading denoting the A(H1N1)pdm09 subtype, orange shading the A(H3N2) subtype). In an equivalent manner, on the right-half of each panel stacked horizontal bars present similar data for type B influenza (cyan shading denoting the B/Victoria lineage, dark blue shading the B/Yamagata lineage). **(a)** Influenza seasons 2009/10 to 2017/18 (inclusive). In the 2009/10 and 2011/12 influenza seasons, all samples identified as influenza B were untyped (represented by black filled bars). **(b)** Influenza seasons 2012/13 to 2017/18 (inclusive), the time span for which we performed parameter inference.

The empirical data did not provide individual vaccine efficacy estimates for each influenza A subtype and influenza B lineage (Table S1). We therefore used a set of simplifying assumptions to produce the strain-specific, vaccine efficacy point estimates used within our study (Table 1). The time-varying vaccine uptake data, together with the season-specific information on vaccine efficacy, act as forcing for the within-season epidemiological model (details are described in Sections 1.4-1.5 of the Supporting Information).

#### Modelling overview

Our motivation was to study the role of immunity propagation in seasonal influenza transmission dynamics by developing and applying a modelling framework incorporating multiple mechanisms linking prior season infection (or vaccination) to immunity in subsequent seasons.

Influenza transmission dynamics are highly complex, with many temporal and structural heterogeneities that will influence the precise pattern of recorded infection. Consequently, our aim was to capture the general trends in the data, such as the pattern of high and low infection levels, rather than the precise values for each influenza season. We, therefore, deliberately chose a parsimonious mechanistic modelling framework, without age-structure, to highlight the impact of immunity propagation.

Following the visualisation convention used in [10], we display a directed acyclic graph representing the model structure and incorporation of the data streams (Fig. 2). The model takes the form of a deterministic continuous-time set of ordinary differential equations (ODEs), which determines the within-season epidemiological dynamics, and a discrete-time map, informing the propagation of immunity from one season to the next.

Our model captures the four strains targeted by the quadrivalent seasonal influenza vaccine, namely, two influenza A subtypes, A(H1N1)pdm09 and A(H3N2), and two influenza B lineages,

**Table 1:** ~~Influenza vaccine efficacy point estimates, by season and strain type.~~ Age adjusted, influenza vaccine efficacy point estimates (by season and strain type) All estimates are presented as percentages. The empirical adjusted influenza vaccine efficacy estimates by season and strain type (presented in Table S1) did not provide individual vaccine efficacy estimates for each influenza A subtype and influenza B lineage. We therefore implemented a series of assumptions to produce the strain-specific, vaccine efficacy point estimates used within our study (described in Section 1.5 of the Supporting Information)).

| Season | Vaccine efficacy | | | | Source |
|---|---|---|---|---|---|
| | A(H1N1)pdm09 | A(H3N2) | B/Victoria | B/Yamagata | |
| 2009/10 | 72.0 | 0.0 | 0.0 | 0.0 | [28]* |
| 2010/11 | 56.0 | 56.0 | 78.0 | 0.0 | [29] |
| 2011/12 | 23.0 | 23.0 | 92.0 | 92.0 | [30] |
| 2012/13 | 73.0 | 26.0 | 51.0 | 51.0 | [31] |
| 2013/14 | 61.0 | 61.0 | 61.0 | 61.0 | [35]† |
| 2014/15 | 29.9 | 29.3 | 46.3 | 46.3 | [32] |
| 2015/16 | 54.5 | 54.5 | 57.3 | 54.2 | [33] |
| 2016/17 | 36.7 | 31.6 | 54.5 | 58.5 | [34] |
| 2017/18 | 66.3 | 0.0 | 24.7 | 24.7 | [36]‡ |

*: The adjusted seasonal influenza vaccine efficacy in the 2009/10 season was -30% (-89%, 11%) [28]. We therefore used the pandemic vaccine to inform efficacy against A(H1N1)pdm09, with the effectiveness against all other types set to zero.

†: Mid-season estimate of seasonal influenza vaccine effectiveness from [35]. Low incidence throughout the 2013/2014 influenza season meant reliable end-of-season estimates for the vaccine efficacy could not be attained (Personal communication, PHE).

‡: Provisional end-of-season influenza vaccine effectiveness results.

B/Victoria and B/Yamagata. The multi-strain model construction permits exploration of cross-reactive immunity mechanisms and their impact on the disease dynamics. Multiple infections per influenza season and/or co-infection events were not permitted. In ~~others~~ other words, it was presumed that individuals may only be infected by one strain of influenza virus per season, analogous to natural infection eliciting short-term cross immunity to all other strain types [37] .

Next, we summarise, in turn, the vaccination, immunity propagation, epidemiological and observation model components.

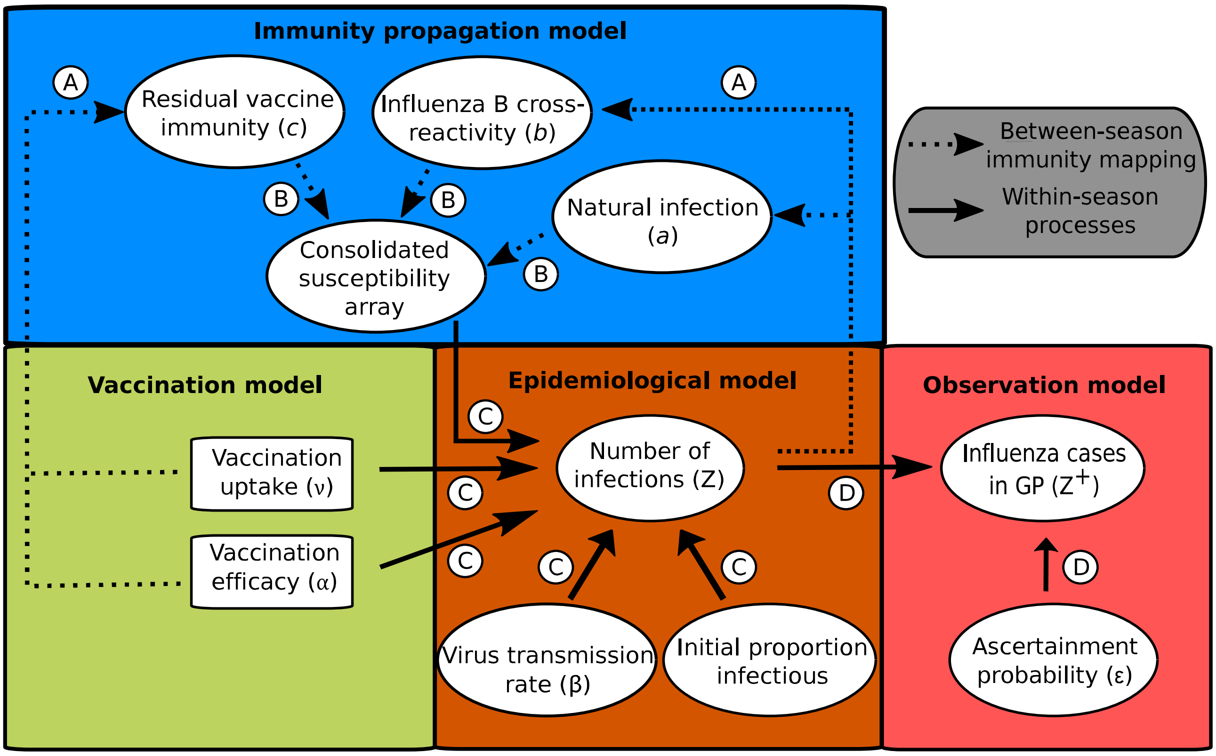


**Fig. 2: Schematic showing the links between the vaccination, immunity propagation, epidemiological and observation model components.** We adopt the visualisation conventions of [10], with ellipses indicating variables, and rectangles indicating data. Dotted arrows indicate relationships between prior season epidemiological outcomes and immunity propagation factors. Solid arrows indicate within-season processes. Circled capitalised letters indicate the relationships connecting the variables or data involved. These relationships are: process A, propagation of immunity as a result of exposure to influenza virus in the previous influenza season (through natural infection or vaccination); process B, modulation of current influenza season virus susceptibility; process C, estimation of influenza case load via the SEIR model of transmission; process D, ascertainment of cases through ILI recording at GP.

We assumed the immunity propagation due to vaccination in the previous season ($c_m$) to be ~~related~~ associated to the strain-specific vaccine efficacy in the previous season ($\alpha_m^{y-1}$). In particular, we introduce a linear scaling factor $\xi \in (0,1)$, which reduces the level of vaccine derived immunity between seasons: $c_m^y = 1 - \xi\alpha_m^{y-1}$.

| | | Strain susceptibility | | | |
|---|---|---|---|---|---|
| | | A(H1N1)pdm09 | A(H3N2) | B/Victoria | B/Yamagata |
| Exposure history (h) | Naïve | 1 | 1 | 1 | 1 |
| | A(H1N1)pdm09 | $a$ | 1 | 1 | 1 |
| | A(H3N2) | 1 | $a$ | 1 | 1 |
| | B/Yamagata | 1 | 1 | $a$ | $b$ |
| | B/Victoria | 1 | 1 | $b$ | $a$ |
| | Vacc. (V) | $c_{A(H1N1)}$ | $c_{A(H3N2)}$ | $c_{B/Victoria}$ | $c_{B/Yamagata}$ |
| | A(H1N1)pdm09 & V | min($a$, $c_{A(H1N1)}$) | $c_{A(H3N2)}$ | $c_{B/Victoria}$ | $c_{B/Yamagata}$ |
| | A(H3N2) & V | $c_{A(H1N1)}$ | min($a$, $c_{A(H3N2)}$) | $c_{B/Victoria}$ | $c_{B/Yamagata}$ |
| | B/Victoria & V | $c_{A(H1N1)}$ | $c_{A(H3N2)}$ | min($a$, $c_{B/Victoria}$) | min($b$, $c_{B/Yamagata}$) |
| | B/Yamagata & V | $c_{A(H1N1)}$ | $c_{A(H3N2)}$ | min($b$, $c_{B/Victoria}$) | min($a$, $c_{B/Yamagata}$) |

**Fig. 3: Infographic presenting the interaction between exposure history and susceptibility.** The interaction between exposure history $h$ and susceptibility to strain $m$ in the current season, $f(h, m)$, was classified into ten distinct groups: One group for the naive (uninfected and not vaccinated, row 1); one group per strain, infected but not vaccinated (rows 2-5); one group for those vaccinated and experiencing no natural infection (row 6); one group per strain for being infected and vaccinated (rows 7-10). We let $a$ denote modified susceptibility to strain $m$ given infection by a strain $m$ type virus the previous season (dark green shading), $b$ modified susceptibility due to cross-reactivity between type B influenza lineages (dark blue shading), and $c_m$ the change in susceptibility to strain $m$ given vaccination in the previous season (gold shading). Unmodified susceptibilities retained a value of 1 (red shading). We enforced $0 < a,b,c_m < 1$.

For use in our observation model, we tracked the incidence $Z_m(y)$ of new strain $m$ influenza infections in season $y$ as a rate per 100,000 population:

$$Z_m(y) = \left( \int_{y-1}^{y} \gamma_{1,m}(E_m^N + E_m^V)\,\mathrm{dt} \right) \times 100,000. \tag{3}$$

##### Observation model

As discussed above, individuals consulting a GP for ILI do not necessarily directly correspond with individuals infected with one of the circulating influenza strains. To address this discrepancy, the final segment of the mathematical model linked the GP consultation data with the number of infections due to circulating influenza viruses in the population (Fig. 2, process D).

Our interest resided in the number of ascertainable cases, where we assumed that each individual infected by the strain of influenza under consideration had a probability $\epsilon$ of being ascertainable, i.e. going to the GP, being recorded as having ILI, and having a detectable influenza viral load.

**Table 2:** Overview of parameters in the model.

| Description | Notation | Value | Sources |
|---|---|---|---|
| **Fixed parameters** | | | |
| Mortality rate ($\text{day}^{-1}$) | B,D | $\frac{1}{81\times365}$ | [45] |
| Rate of latency loss, influenza A subtypes ($\text{day}^{-1}$) | $\gamma_{1,A}$ | $\frac{1}{1.4}$ | [43] |
| Rate of latency loss, influenza B lineages ($\text{day}^{-1}$) | $\gamma_{1,B}$ | $\frac{1}{0.6}$ | [43] |
| Recovery rate ($\text{day}^{-1}$) | $\gamma_2$ | $\frac{1}{3.8}$ | [44] |
| **Time-varying parameters** | | | |
| Vaccination rate at time t | $\nu(t)$ | — | [24, 25, 27] |
| Vaccine efficacy, season $y$ strain $m$ | $\alpha_m^y$ | — | [28–36] |

$$\text{DEV} = 2\sum_{y}\sum_{m}\left(C_{m,y}\ln\left(\frac{C_{m,y}}{M_{m,y}}\right) - (C_{m,y} - M_{m,y})\right), \tag{5}$$

with $C_{m,y}$ the observed value for strain $m$ in season $y$, and $M_{m,y}$ the model estimate for strain $m$ in season $y$.

We amassed 10,000 particles representing a sample from the posterior distribution (see Section 3 of the Supporting Information for expanded details on the parameter estimation methodology).

#### Forward simulations

We used the parameter set returning the lowest error (in the inference procedure) to explore potential strain dynamics up to the 2029/~~2030~~ 30 influenza season. Vaccination attributes (uptake and efficacy) for the 2009/~~2010~~ 10 to 2017/~~2018~~ 18 influenza seasons were taken from the observed data. For the forecasted seasons (2018/19 influenza season to the end of each simulation), we studied ~~two~~ four scenarios arising from different hypotheses concerning future vaccine efficacy: (i) ~~high efficacy ,~~ randomly sampled from all previous vaccine efficacy values (1,000 simulation replicates); (ii) 'expected' scenario (single simulation), strain-specific vaccine efficacies set at the ~~maximum attained efficacy (~~median attained efficacy across 2010/11-2017/18 seasonal influenza vaccines (note, efficacy estimates from 2009/10 influenza season were not included as for 2009/10 we used pandemic influenza vaccine efficacy estimates); (iii) pessimistic scenario (single simulation), strain-specific vaccine efficacies set at the minimum attained efficacy across 2010/11-2017/18 seasonal influenza vaccines; and (~~ii) randomly sampled from all previous vaccine efficacy values (1~~iv) optimistic scenario (single simulation), ~~000 simulationreplicates)~~strain-specific vaccine efficacies set at the maximum attained efficacy across 2010/11-2017/18 seasonal influenza vaccines. In addition, vaccine uptake in all projected influenza seasons matched that of the 2017/18 influenza season.

**Table 3:** ~~**Values (posterior quantiles) for the transmission, exposure history and ascertainment probability parameters inferred fitting to the empirical data.**~~ **Values (posterior weighted quantiles) for the transmission, exposure history and ascertainment probability parameters inferred fitting to the empirical data.** Numbers inside brackets indicate 95% credible intervals. All values are given to 4 d.p.

| Description | Notation | Median [95% credible interval] |
|---|---|---|
| **Transmission parameters** | | |
| A(H1N1)pdm09 | $\beta_{\text{A(H1N1)pdm09}}$ | ~~0.3955~~ 0.3913 [~~0.3880, 0.4039~~0.3900, 0.4041] |
| A(H3N2) | $\beta_{\text{A(H3N2)}}$ | ~~0.3984~~ 0.3917 [~~0.3904, 0.4076~~0.3917, 0.4074] |
| B/Victoria | $\beta_{\text{B/Victoria}}$ | ~~0.3548~~ 0.3510 [~~0.3498, 0.3603~~0.3510, 0.3606] |
| B/Yamagata | $\beta_{\text{B/Yamagata}}$ | ~~0.3694~~ 0.3653 [~~0.3635~~0.3653, 0.3759] |
| **Exposure history parameters** | | |
| Natural infection in prior season | $a$ | ~~0.7898~~ 0.7883 [~~0.7834, 0.7966~~0.7847, 0.7964] |
| Type B influenza cross-reactivity | $b$ | ~~0.9797~~ 0.9703 [~~0.9511, 0.9983~~0.9558, 0.9988] |
| Prior season vaccine efficacy propagation | $\xi$ | ~~0.0061~~ 0.0051 [0.0004, ~~0.0163~~0.0143] |
| **Ascertainment probabilities** | | |
| 2012/13 | $\epsilon_{2012/13}$ | 0.0017 [0.0015, 0.0018] |
| 2013/14 | $\epsilon_{2013/14}$ | ~~0.0010~~ 0.0009 [0.0009, 0.0012] |
| 2014/15 | $\epsilon_{2014/15}$ | 0.0024 [0.0022, 0.0026] |
| 2015/16 | $\epsilon_{2015/16}$ | ~~0.0036~~ 0.0037 [~~0.0033, 0.0040~~0.0032, 0.0039] |
| 2016/17 | $\epsilon_{2016/17}$ | 0.0014 [0.0012, 0.0015] |
| 2017/18 | $\epsilon_{2017/18}$ | ~~0.0051~~ 0.0055 [0.0047, 0.0055] |

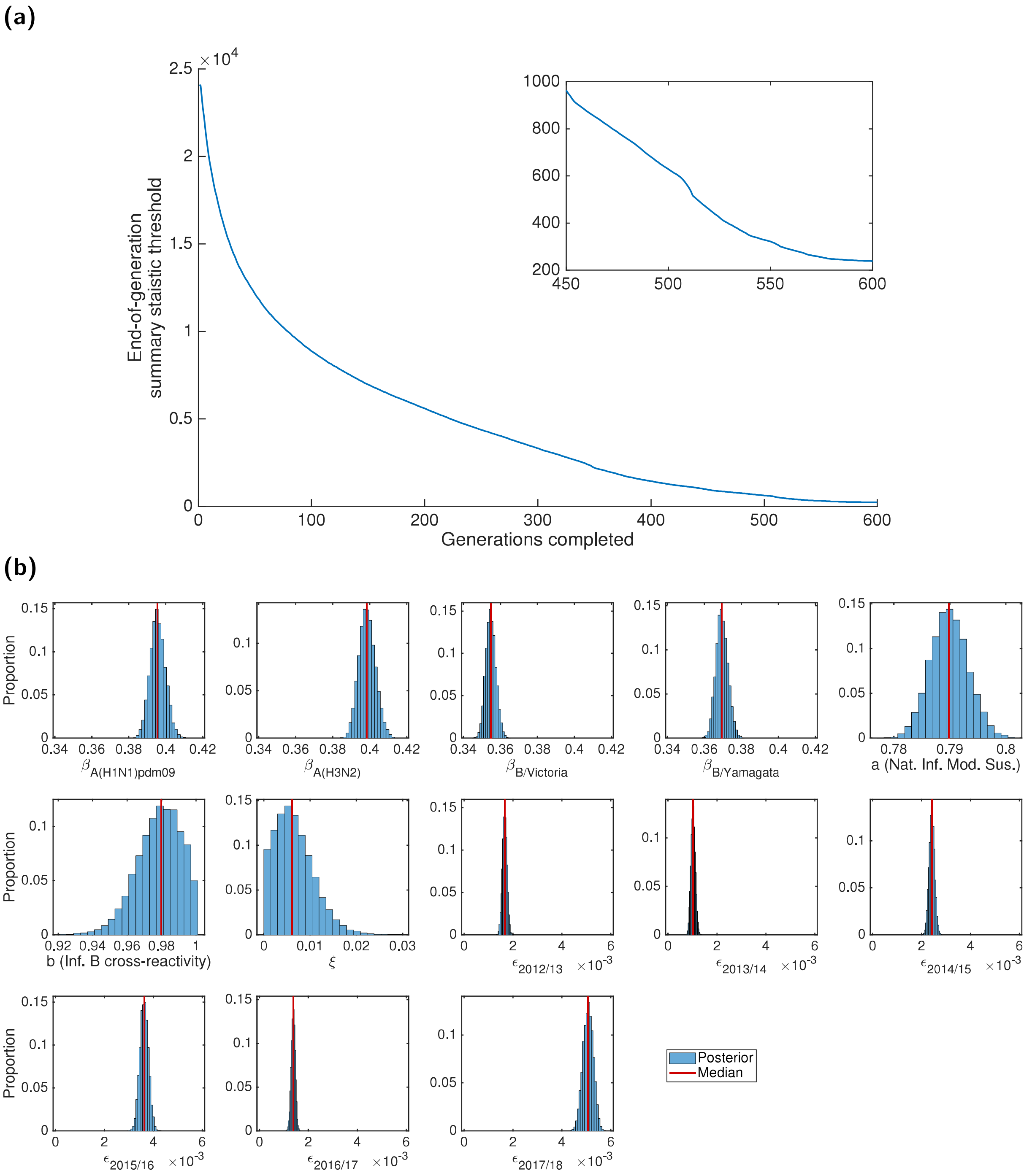


**Fig. 4: Results of the ABC scheme, fitting to the empirical data.** **(a)** Summary metric threshold value upon completion of each generation of the inference scheme. The inset panel displays the latter quarter of generations. **(b)** Inferred parameter distributions estimated from 10,000 retained samples following completion of 600 generations of the inference scheme. Vertical red lines indicate the (non-weighted) median values for the model constants estimated from the inference procedure. Particularly noteworthy outcomes include: transmissibility of type A viruses exceeding type B viruses; prior season influenza B cross-reactivity and vaccine carry over had little impact on present season susceptibility; the highest ascertainment probability occurred in the 2017/18 influenza season.

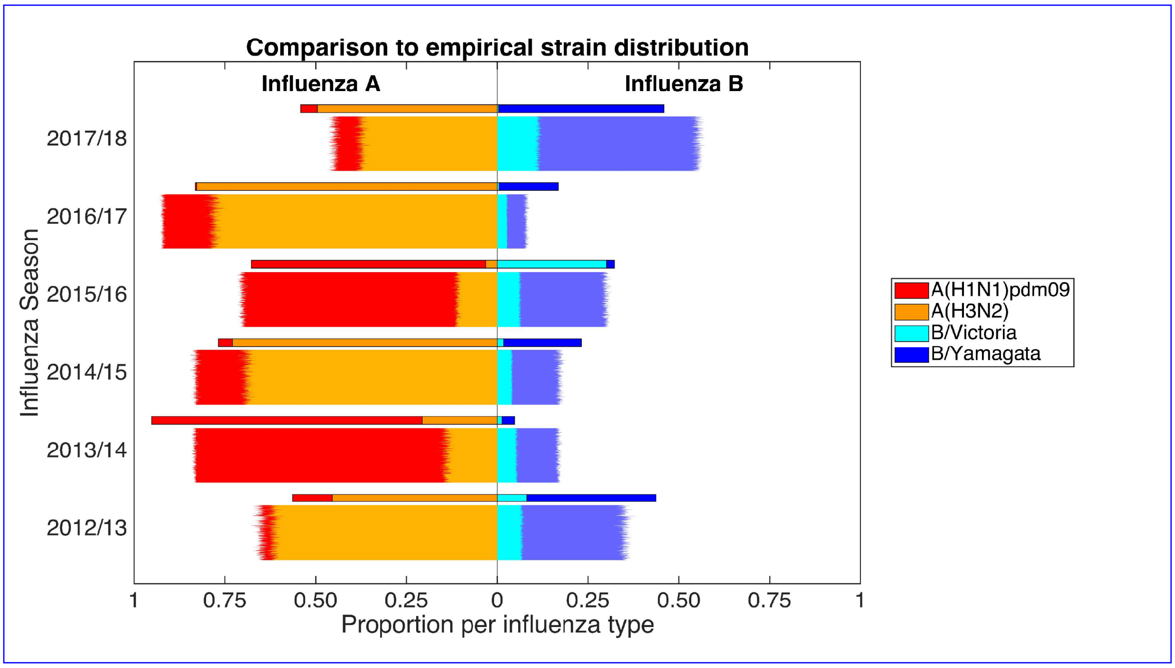


**(b)**

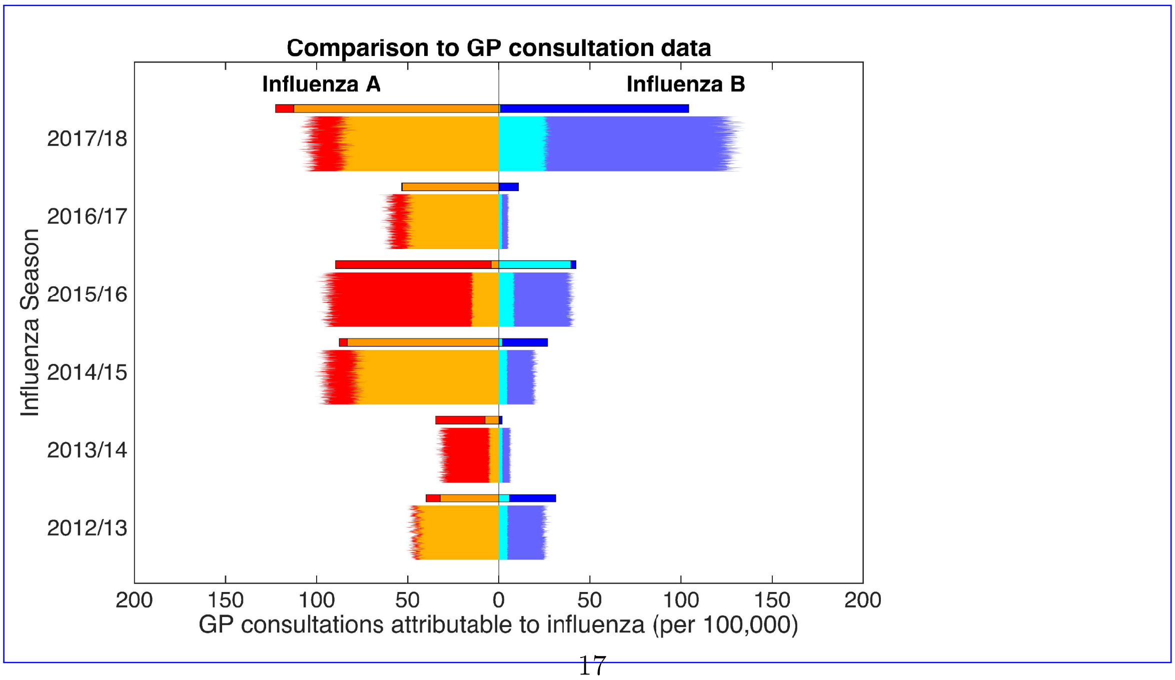




**Fig. 5:** **Posterior predictive distributions for strain composition and influenza positive GP consultations per 100,000 population.** Stratified by influenza season, we present back-to-back stacked bars per simulation replicate, with 1,000 replicates performed, each using a distinct parameter set representing a sample from the posterior distribution. Each influenza season is topped out by a thicker stacked horizontal bar plot, corresponding to the strain-stratified point estimates for the empirical data. (a) Posterior predictive distributions for circulating influenza virus subtype/lineage composition. (b)

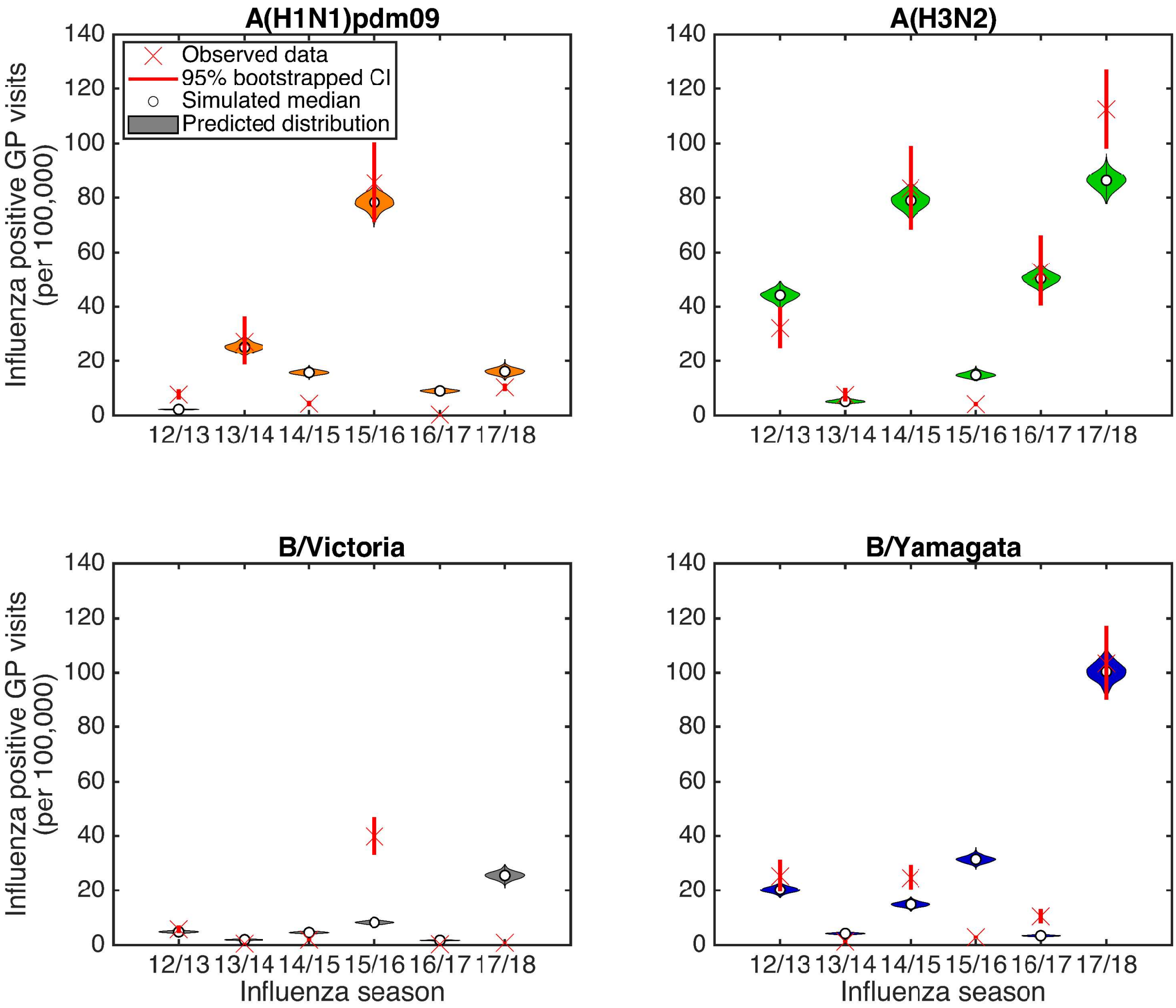


**Fig. 6: Posterior predictive influenza positive GP consultation distributions.** We generated the estimated distributions from 1,000 model simulations, each using a distinct parameter set from the retained collection of particles. Crosses represent the observed data, with solid bars the range of the bootstrapped empirical data. Shaded violin plots denote outcomes from model simulations using the empirical data on vaccination, with filled circles the median value across the simulated replicates. **(a)** A(H1N1)pdm09; **(b)** A(H3N2); **(c)** B/Victoria; **(d)** B/Yamagata.

#### Forward simulations

Using the parameter set from the inference scheme returning the closest correspondence with the data (lowest DEV value, Eq. (5)), we explored potential strain dynamics up to the 2029/30 influenza season under contrasting vaccine efficacy settings (Fig. 7).

When maintaining ~~the same high~~ 'expected' vaccine efficacy (A(H1N1)pdm09: ~~73~~55.25%; A(H3N2): ~~61~~30.45% B/Victoria: ~~92~~55.90%; B/Yamagata: ~~92~~52.60%) across all future seasons (Fig. 7(a), single bars), ~~consistent levels of influenza incidence were predicted~~ we witness minor variability in predicted influenza incidence for both influenza types. ~~Whilst combined~~ Total influenza B incidence ~~was low, ranging approximately between 600-1700~~ ranged between 2,700-10,250 cases per 100,000 each season, and seasonal influenza A incidence consistently reached 25,~~000-30~~000-40,000 cases per 100,000. Furthermore, we ~~again~~ predict periodic behaviour for the portion of influenza A viruses in circulation ascribed to the subtypes A(H1N1)pdm09 and A(H3N2), mirroring the observed pattern from 2012/13 to 2016/17. Influenza seasons ending in ~~odd~~ even numbered years (e.g. ~~2028~~2029/~~29~~30) had a relatively even split between the two subtypes; whereas for influenza seasons ending in ~~even~~ odd numbered years (e.g. ~~2029~~2028/~~30~~29) the A(H3N2) subtype dominated the A(H1N1)pdm09, with influenza A viruses apportioned to A(H3N2) and A(H1N1)pdm09 subtypes being ~~75-80% and 20-25~~70-85% and 15-30%, respectively.

An alternative approach is to use the single (best-fit) parameter set, but to sample the vaccine efficacies from historical estimates. The most striking factor was the amount of variability that comes as a direct results of uncertainty in the vaccine efficacies. As such all between-season patterns were swamped by the within-season variability, leading to distributions that were remarkably consistent between seasons. The only notable exception was 2018/19, where the definitive knowledge of the 2017/18 season had a clear impact of the predictions for 2018/19 leading to broadly higher levels of influenza A and lower levels of influenza B ~~. —~~ (Fig. 7(a)).

Comparing subtype distributions when vaccine efficacies were either sampled from historical estimates or fixed at 'expected' values, we found that under randomly sampled vaccine efficacies incidence of B/Yamagata tended to exceed B/Victoria in each season (holding in more than 70% of simulations, Figure S18). However, we obtained greater levels of variation for the influenza A subtypes. ~~For high vaccine efficacy conditions~~ In an 'expected' vaccine efficacy setting (single bars), less than half of total influenza A cases in any season were attributed to the A(H1N1)pdm09 subtype (rather than A(H3N2)), whereas under variable vaccine efficacies (distributed bars) A(H1N1)pdm09 incidence was greater than A(H3N2) incidence in 30-50% of all simulations ~~. Unsurprisingly,~~ (Figure S18). Furthermore, in each future influenza season the majority of simulations (∼~~85~~60-75%) performed with seasonally variable vaccine efficacies ~~returned predicted incidence rates (in all influenza seasons)that exceeded the equivalent seasonal incidence estimatesgenerated under high vaccine efficacy conditions; this was most striking for influenza B.~~ resulted in overall incidence rates exceeding the equivalent seasonal incidence estimates generated under 'expected' vaccine efficacy conditions.

~~ht!**Projected influenza seasonal incidence (per 100,000 population) up to 2029/30.** Each influenza season is topped out by a thicker stacked horizontal bar plot, corresponding to the simulated estimate under the high vaccine efficacy scenario; efficacies by strain were as follows - A(~~The impact of vaccine efficay on projected incidence is further exemplified by the stark contrast in predictions when comparing the pessimistic scenario (A(H1N1)pdm09: ~~73%; A(~~23.00%; A(H3N2): 0.00%; B/Victoria: 24.70%; B/Yamagata: 0.00%) to the optimistic scenario (A(H1N1)pdm09: ~~61%; B~~73.00%; A(H3N2): 61.00% B/~~Victoria: 92%; B~~Victoria:

92.00%; B/Yamagata: 92%. Thin bars correspond to simulation runs where vaccine efficacy against each strain were randomly sampled from the empirical distribution (totalling 1Yamagata: 92.00%). Under pessimistic vaccine efficacy conditions, both influenza types underwent vast fluctuations in incidence from one influenza season to the next. Type A was more severe in odd numbered years (e.g. 2028/29). Similarly, in even numbered years (e.g. 2029/30) type B incidence exceeded type A incidence. For the pessimistic vaccine efficacy scenario (compared to the other fixed vaccine efficacy scenarios) we also observe the strongest re-emergence of the alternating behaviour between dominant influenza A subtypes (Fig. 7(b)).

Maintaining a high vaccine efficacy, both influenza types had consistent incidence estimates. Whilst combined influenza B incidence was low, ranging approximately between 600-1700 cases per 100,000 replicates). The left side depicts the cumulative total of ILI GP consultations attributable to type A influenza per 100each season, seasonal influenza A incidence frequently reached 25,000-30,000 population (red shading denoting the A(cases per 100,000. Although we, once more, predict periodic behaviour for the portion of influenza A viruses in circulation ascribed to the subtypes A(H1N1)pdm09 subtype, orange shading the A(and A(H3N2), under these optimistic vaccine efficacy conditions it was influenza seasons ending in odd numbered years having a relatively even split between the two subtypes; whereas for influenza seasons ending in even numbered years the A(H3N2) subtype dominated A(H1N1)subtype)pdm09 (Fig. 7(c)). The right side stacked horizontal bars present similar data for type B influenza (cyan shading denoting the B/Victoria lineage, dark blue shading the B/Yamagata lineage). For all forward simulated seasons, vaccine uptake matched that of the 2017/18 influenza season.

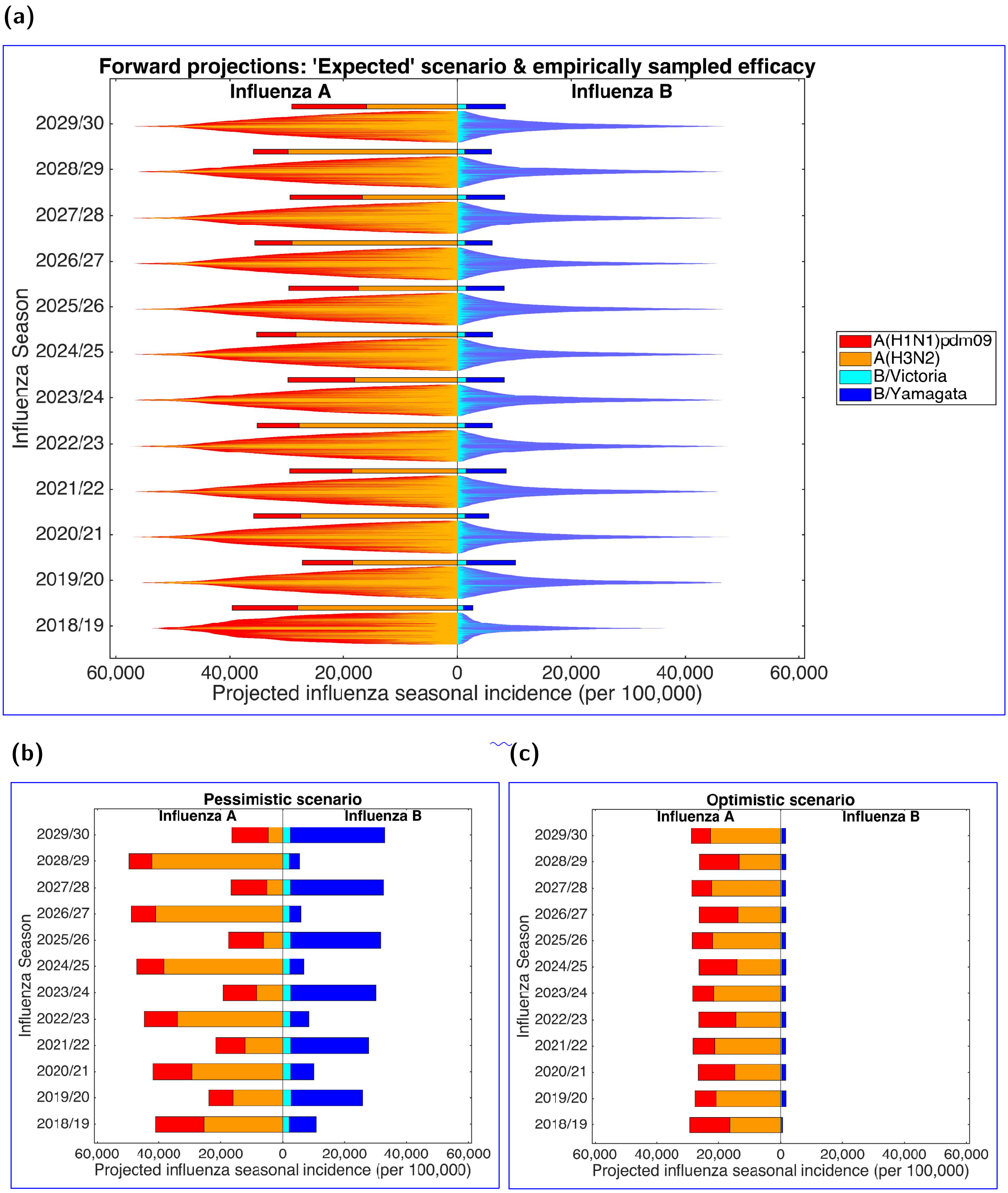


**Fig. 7:** **Projected influenza seasonal incidence (per 100,000 population) up to 2029/30.** For all scenarios, in forward simulated seasons vaccine uptake matched that of the 2017/18 influenza season. In all panels, the left side depicts the seasonal incidence of type A influenza per 100,000 population (red shading denoting the A(H1N1)pdm09 subtype, orange shading the A(H3N2) subtype). The right side stacked horizontal bars present projected seasonal incidence of type B influenza per 100,000 population (cyan shading denoting the B/Victoria lineage, dark blue shading the B/Yamagata lineage).**(a)** Each influenza season is topped out by a thicker stacked horizontal bar plot, corresponding to the simulated estimate under the 'expected' vaccine efficacy scenario; efficacies by strain were as follows - A(H1N1)pdm09: 55.25%; A(H3N2): 30.45%; B/Victoria: 55.90%; B/Yamagata: 52.60%. Thin bars correspond to simulation runs where vaccine efficacy against each strain were randomly sampled from the empirical distribution (totalling 1,000 replicates). Simulations are arranged such that the largest epidemics are in the centre of the bar. **(b)** Projected incidence under a pessimistic vaccine efficacy scenario; efficacies by strain were as follows - A(H1N1)pdm09: 23.00%; A(H3N2): 0.00%; B/Victoria: 24.70%; B/Yamagata: 0.00%. **(c)** Projected incidence under an optimistic vaccine efficacy scenario; efficacies by strain were as follows - A(H1N1)pdm09: 73.00%; A(H3N2): 61.00%; B/Victoria: 92.00%; B/Yamagata: 92.00%.

### Discussion

Understanding the processes that drive seasonal influenza transmission patterns is an important public-health question, given the economic cost and burden to human health afflicted by the condition [1, 2]. During recent years there has been marked progress in synthesising real-world data to inform mathematical model parameterisation. Yet, prior studies have tended to model each season and each strain circulating within that season independently ~~[10–13]~~[10, 12, 13]. Thus, the presence of strain and immunity interactions have been omitted. Using data subsequent to the 2009 pandemic for England, we have developed a dynamic multi-strain SEIR-type transmission model for seasonal influenza, explicitly incorporating immunity propagation mechanisms between seasons. ~~Our model is fitted to seasonal-level data on strain competition, with~~

With a view to minimising the number of independent parameters~~. Predictions from this model show substantial agreement with~~, we fit a parsimonious mechanistic model to seasonal-level data on strain competition. In spite of the multi-strain complexity and time scale of the study period (six influenza seasons), predictions from the model attain a strong qualitative resemblance (in terms of strain composition and overall quantity of GP consultations) to the empirical subtype data from England~~, and are able to predict the main qualitative features~~. We attribute much of the discrepancies to the homogeneous way in which we have treated immunity propagation, as in practise this is driven by complex patterns of waning and cross immunity as well as somewhat irregular genetic drift.

Inferred transmissibility of type A influenza strains exceeding those of type B, which reflects type A being the predominant class of influenza virus in circulation over the studied time period [26]. Moreover, concentrating on the parameters relevant to immunity propagation, we uncover evidence against vaccination stimulating similar long-term immunity responses as for natural infection. These conclusions corroborate ~~findings of broad immune response (against antigenically drifted strains) fading after repeated vaccination [48], plus rapid waning of~~ previous immunological studies showing that infection with influenza virus can induce broader and longer-lasting protection than vaccination [37, 49, 50], and authenticate prior work signalling that vaccine-mediated immunity rapidly wanes [51]. There are also indications that prior natural infection boosts vaccine responses against antigenically drifted strains, whereas prior vaccination does not [52]. The clinically observed impact of prior infection for enhancing vaccine efficacy was long-lasting, which may be used to instruct further model refinements.

**Writing - original draft:** Edward M. Hill.

**Writing - review & editing:** Edward M. Hill, Matt J. Keeling, Stavros Petrou, Simon de Lusignan, Ivelina Yonova.

### Financial disclosure

EMH, SP and MK are supported by the National Institute for Health Research [Policy Research Programme, Infectious Disease Dynamic Modelling in Health Protection, grant number 027/0089]. The funders had no role in study design, data collection and analysis, decision to publish, or preparation of the manuscript.

### Data availability

The GP consultation data contain confidential information, with public data deposition non-permissible for socioeconomic reasons. The GP consultation data resides with the RCGP Research and Surveillance Centre and are available via the RCGP RSC website (www.rcgp.org.uk/rsc). All other raw data utilised in this study are publicly available; relevant references and data repositories are stated within the main manuscript and ~~supporting information~~Supporting Information. Code and processed data used for the study is available at https://github.com/EdMHill/SeasonalFluImmunityPr[illegible]

### Competing interests

The authors declare that they have no competing interests.
